# Does charging for corrections in the bioscience literature disincentivize pre-publication handling of problematic image data? An ImageTwin-AI study

**DOI:** 10.64898/2026.01.16.700000

**Authors:** Paul S. Brookes

## Abstract

The Committee on Publication Ethics (COPE) recommends that publishers do not charge for corrections to published papers. Until late 2024 the *Journal of Cancer* levied a charge on authors (50% of the original article processing charge, APC) for publication of a correction. Herein, it was hypothesized this could disincentivize the discovery and removal of problematic data prior to publication, since post-publication discovery and correction would generate additional revenue. The correction charge policy at *J. Cancer* was rescinded in 2025, permitting a test of the hypothesis by comparing the prevalence of problematic image data in the journal before and after the policy change. Recently developed artificial intelligence (AI) tools afford the ability to screen scientific publications for problematic image data. As such, the 2024-2025 output of *J. Cancer* was analyzed using ImageTwin-AI, followed by human verification and annotation of identified problems. Of 754 papers analyzed, 510 contained image data. Of these, 95 (18.6 %) showed evidence of inappropriate image manipulation, with 19 papers (3.7 %) having images that overlapped with unrelated papers. The prevalence of papers with problem images was 20.3% in 2024, and 15.9% in 2025, suggesting only a modest impact of the policy change on pre-publication handling of such problems.

## Introduction

For most of the modern scientific era, publication of scientific results was limited to a handful of specialist publishing companies, whose revenues originated from journal subscription fees paid by libraries. Limited library budgets restricted the number of journals an institution could subscribe to, such that libraries and librarians played an unseen yet important role in determining the size of the publishing market and indirectly policing the quality of journal content.

The explosive growth of digital open access (OA) publishing ^[1,2]^, while removing global accessibility barriers to knowledge, has been accompanied by an unanticipated proliferation of commercial journals, funded by an *author pays* financial model (i.e., article processing charges, APCs). While journal quality is hard to measure, it is generally regarded that growth of pay-to-publish journals has been accompanied by a decline in quality. This is embodied by the concept of *paper-mills*, in which authors (often with no laboratories) pay a fee to a 3^rd^ party broker who prepares a paper using images and other data stolen from online resources, submits it and handles the peer review process ^[3]^. Several paper-mill networks have been uncovered, causing publishers and journals to retract thousands of papers *en-masse* ^[4]^.

The prevalence of research misconduct has been measured by several methods, with estimates ranging from 1-4% based on self-reporting ^[5-7]^, rising to >20% if the question is framed differently ^[8]^. Post-hoc studies on published papers estimate the prevalence of inappropriate image manipulation at 16% ^[9]^ or 2-4% ^[10]^, while paper-mill products have been estimated to comprise 3% of the literature ^[11]^. While misconduct is the reason that most scientific papers get retracted, such retractions only represent 0.02% of the literature ^[12]^, suggesting a mismatch between the amount of misconduct occurring, and the degree to which the published results of misconduct are appropriately addressed. This has led one prominent scientific investigator to claim that 1 in 7 of all scientific papers is fake ^[13]^, Furthermore, although I have shown that internet publicity of integrity issues in papers leads to enhanced levels of corrective action ^[14]^, and databases of retracted and problematic papers can be readily incorporated into bibliographic workflows ^[15,16]^, nevertheless most retracted papers continue to be cited ^[17]^.

The relationship between journal editorial policies and the prevalence of problem data in the scientific literature has not been extensively studied. Although contrary to the guidelines of the Committee on Publication Ethics (COPE) ^[18]^, some journals levy an additional charge on authors for the issuance of an erratum or correction to a published paper. Such fees yield additional revenue for any problems discovered post-publication, leading to the current hypothesis that such fees may lower incentives to discover and remove problems during pre-publication. This hypothesis was tested by analyzing papers from the *Journal of Cancer* (Ivyspring Publishing), an OA journal that follows COPE guidelines ^[19]^, but nevertheless levied a charge of 50% of the original APC for corrections/errata until the end of 2024 ^[20]^. The rescinding of this policy in 2025 affords a simple pre-/ post-comparison.

The advent of consumer-grade machine learning (ML) and artificial intelligence (AI) computing has led to the widespread availability of online tools to screen scientific papers for problems that may be suggestive of inappropriate image manipulation ^[21-23]^. Herein, the 2024– 2025 content from *J. Cancer* was screened using ImageTwin-AI. Within the study cohort, 18.6% of papers containing any image data showed evidence of inappropriate image manipulation. The fraction of problematic papers was slightly lower in 2025 (15.9%) vs. 2024 (20.3%), suggesting that adoption of COPE guidelines on charging for corrections was not associated with a large improvement in the discovery and removal of problem images pre-publication. As such the hypothesis is rejected.

Notably, 3.7% of papers contained images traceable to papers in other journals, which would have been impossible to discover manually. This highlights the value of AI tools for paper screening, and suggests that editorial workflows should engage in more rigorous screening of papers prior to publication.

## Methods

The origins and pipeline for this study are shown in Figure 1. The 2024 and 2025 scientific content of the *Journal of Cancer* was downloaded from the journal website ^[24]^ in early 2025 and early 2026, respectively. Only papers reporting original scientific results were studied, with exclusion of review articles, editorials and other forms of journal content. A total of 754 papers were analyzed.

**Figure 1.**
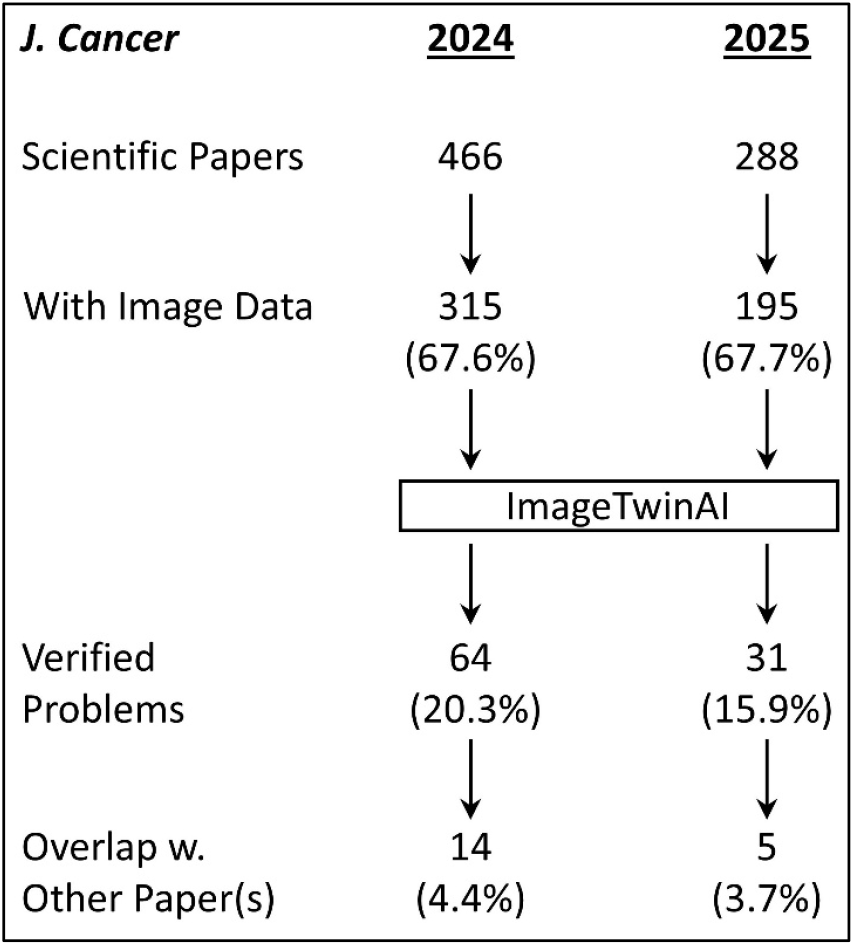
Outline and flowchart for the current study

Since the ImageTwin-AI screen is image-based, papers containing only graphs, charts, tables or other types of numerical or computer-generated data were also excluded from the analysis. 510 papers (67.6%) contained image data in the following formats: (i) western blots; (ii) fluorescent microscopy images; (iii) light microscopy images of cells; (iv) light microscopy images of tissue cross sections (histology); (v) gross anatomy or images of whole tissues; (vi) flow cytometry data; (vii) photographs.

Each PDF paper was uploaded to ImageTwin-AI and subjected to screening, to look for common image manipulations including but not limited to: (i) image re-use or duplication; (ii) reuse with rotation, transposition, flipping, resizing, recoloration, altered brightness or contrast; (iii) reuse within a figure, across multiple figures in a paper, or across multiple papers; (iv) splicing of western blots. Problems flagged by ImageTwin were verified manually, with compilation into report files and annotation using PowerPoint software. Although ImageTwin offers a new capability to detect AI-generated images, this feature was not employed.

## Results

Papers which contained images taken from public databases (e.g. Human Protein Atlas ^[25]^ or Genome Expression Profiling Interactive Analysis ^[26]^) with acknowledgement or citation were frequently tagged by ImageTwin as matching other papers, but such instances were not considered to be positives in this screen. A similar exclusion was applied to cases where the same loading control was used for western blots under the same experimental conditions, or the same control cell images were used across multiple experiments. Although such image reuse does not follow *best practices* for data presentation, it is nonetheless an accurate report of the scientific record.

Of 754 papers that contained image data and were screened, 95 contained evidence of inappropriate image manipulation, and contained 153 total problematic figures. Notably, 19 papers contained images that matched those found in other papers and other journals. Such examples would have been almost impossible to find manually, without the significant database of images contained within the ImageTwin-AI platform. These papers can be considered the most *suspicious* in terms of potential misconduct, since there can rarely be a legitimate explanation for re-using an image from another paper without citation.

Several examples of the types of problematic images identified by ImageTwin-AI and verified by human eye are shown in Figures 2-7. In addition, Supplemental files 1 and 2, and Supplemental Spreadsheet 1 contain full documentation of all the image data problems identified in this study.

**Figure 2.**
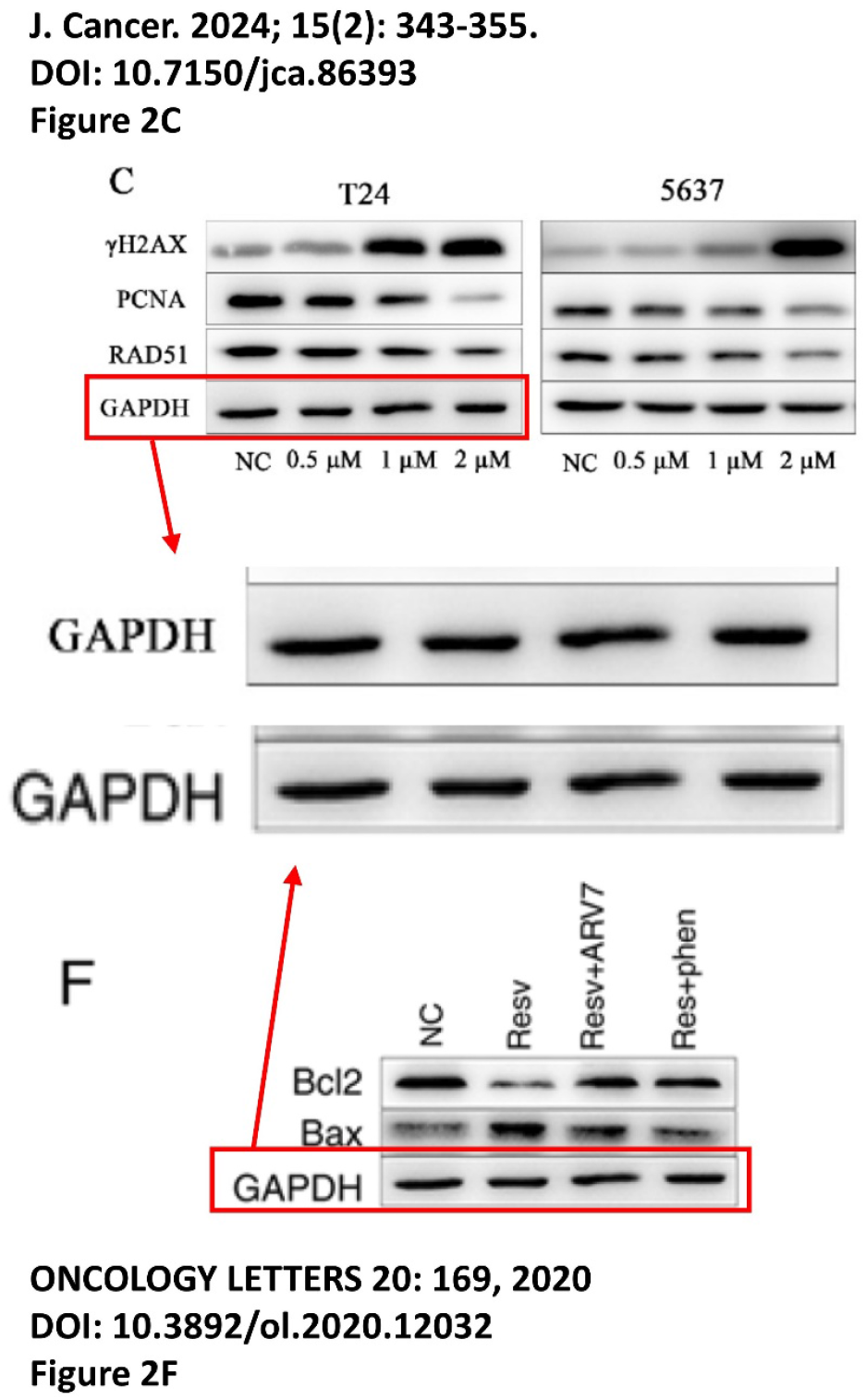
Example of apparent image duplication between western blots in two unrelated papers.

**Figure 3.**
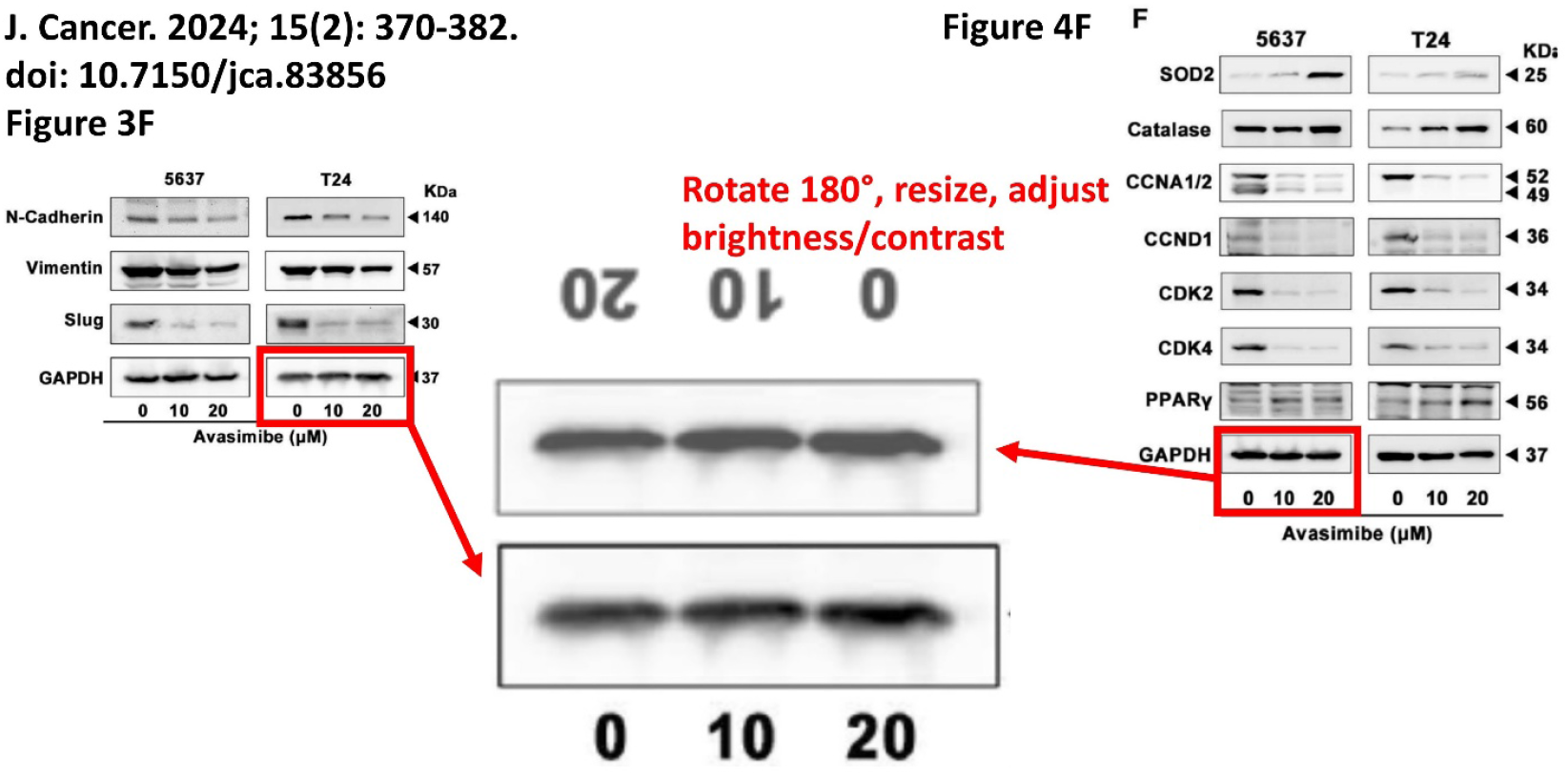
Example of apparent image duplication between western blots in two different figures in a paper, with 180° rotation, resizing, and adjustment of brightness/contrast.

**Figure 4.**
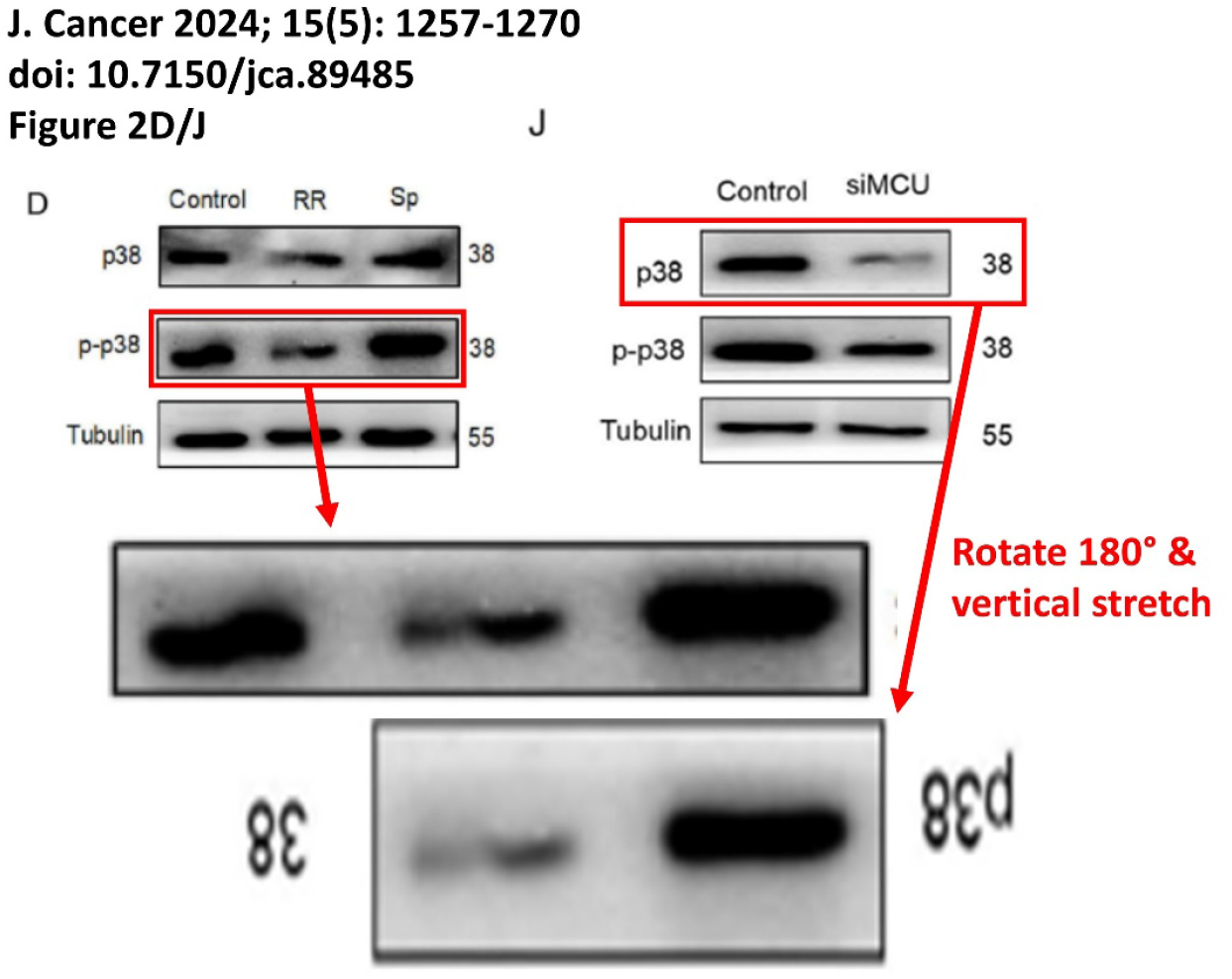
Example of apparent image duplication between western blots in two different figure panels in a paper, with 180° rotation and resizing.

**Figure 5.**
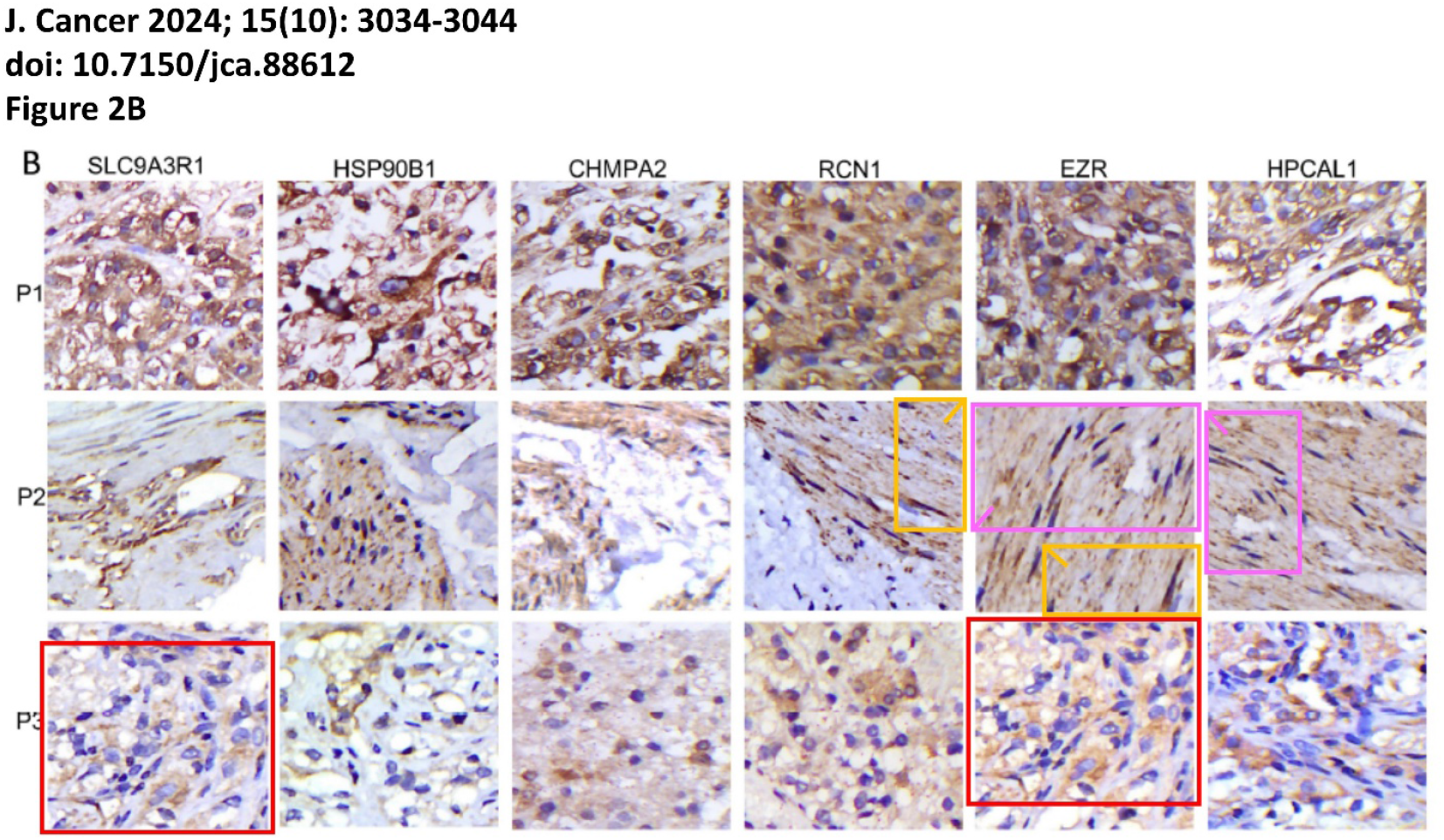
Example of multiple apparent image duplications between figure panels within a paper. Colored boxes show common features, with corner markers used to indicate apparent rotation of images.

**Figure 6.**
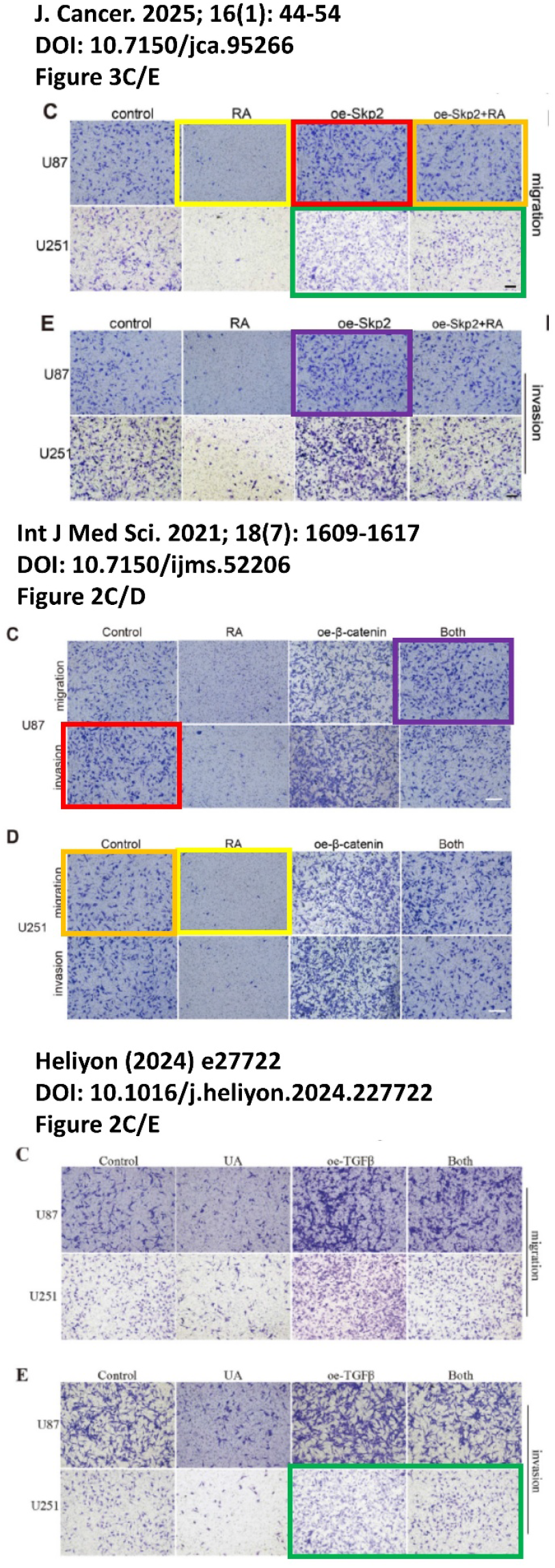
Example of multiple apparent image duplications between 3 separate papers. Colored boxes show common features.

**Figure 7.**
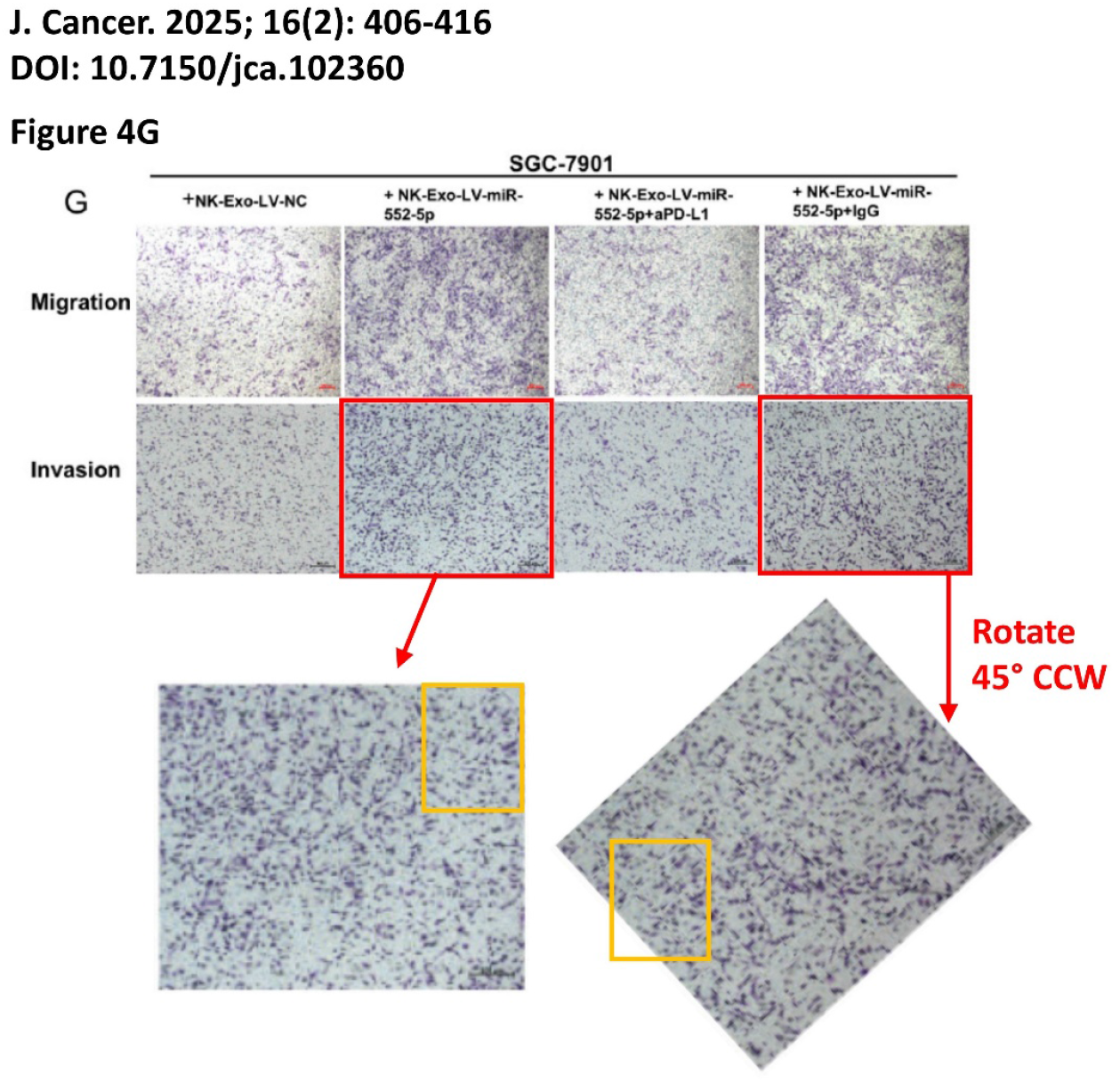
Example of apparent image duplica..ons between figure panels within a paper. Colored boxes show common features, with apparent rotation of images.

Addressing the hypothesis that the prevalence of problem data may be related to whether or not a journal charges a fee for publication of a correction, this does not appear to be the case. The fraction of papers with problem images was 20.3% in 2024 (while such a charge was in place) and 15.9% in 2025 (after the charge was removed), suggesting little if any relationship between this change in editorial policy and the rate at which problematic image data are addressed during pre-publication.

## Discussion

The main conclusions of this work are: (i) AI-assisted detection is a powerful and valuable tool for screening papers in the bioscience literature for problematic image data. It is uniquely capable of identifying shared images across otherwise unrelated papers. (ii) The imposition (or not) of a fee for correcting published work does not appear to have any relationship with the prevalence of problematic image data. It is unclear whether the small decline in prevalence between 2024 and 2025 can be attributed to the removal of the fee policy at *J. Cancer*.

The proportion of problematic papers identified in this study (18.6%) is in line with a previous finding of 16% ^[9]^, and is consistent with the general idea that 1 in 7 papers (14.3%) are problematic. These numbers are much greater than the 2-4% concluded in a 2016 study ^[10]^, which could be attributed to either large differences in methodology, or improvements in detection technology in the intervening decade. Notably these numbers are also much larger than the estimated prevalence of scientific misconduct (approx.’ 2 % based on self-reporting).

There are a number of caveats and limitations to this study that warrant discussion. Firstly, this is a relatively small study of 2 years’ of output from a single journal. It is therefore unclear whether the results are generalizable to the scientific literature at large. Secondly, image data of the type studied herein can be considered “low hanging fruit” for the detection of potential inappropriate manipulation. A significant portion (32.4%) of papers were excluded from this analysis because they did not contain any image data. Nevertheless, there is the possibility that the data within these papers (graphs, charts, etc.) could also be problematic. A number of approaches have been described for the forensic detection of problems in numerical data ^[27-30]^, although these often depend on having access to the original numbers used to generate figures.

Thirdly, while this study highlights the use of an AI platform for detection of problematic images, such systems are not without their own problems. For example, many generative AI and machine learning platforms use both the data input by users during the query process, along with user-feedback, to help train the model. This can create problems of data privacy and confidentiality, so for this reason most journals and funding agencies prohibit the use of AI in the peer review process. It is important to note that all of the papers studied herein were post-publication and available OA, so are already in the public domain and possibly already within the training sets of AI platforms. Thus, although tools such as ImageTwin-AI are useful for analyzing already published work, it is not clear whether they could be safely implemented during pre-publication workflows. Additionally, it is important to apply a *trust-but-verify* approach with all AI tools. Herein, all flagged images were subject to human verification and documentation, as shown in the Supplemental files.

Fourthly, there are several potential confounders for testing the hypothesis on the relationship between charging fees for corrections and the incentives for correcting papers prior to publication. The test was based on analysis of a single year of data before and after existence of the fee policy, and there is a notable drop in the total number of papers published from 2024 (466) to 2025 (288). It is possible this 38% drop reflects a larger number of papers being rejected due to problems being caught prior to publication. It is not known how the change in editorial policy was communicated to editors, or whether any additional tools or policies were in place between the two years. Nevertheless, the decrease in prevalence of problematic images from 20.3% to 15.9% (a 21.7% drop) is smaller than the decrease in publication volume, and suggests that a significant portion of problematic image data is still escaping pre-publication detection.

Lastly, it should be emphasized that none of the image data problems documented herein should be construed as implying that scientific misconduct has occurred. There may be reasonable explanations for occurrence of such problems, and the editors of *J. Cancer* are encouraged to engage with authors to resolve these issues.

## Supporting information

Supplemental file 1 (2024 content)

Supplemental file 2 (2025 content)

Spreadsheet with data

## Acknowledgements and Conflict of Interest

The author was an early beta-tester for the ImageTwin-AI platform, and as such acknowledges free access to the website and its tools. No funding was received for the work described herein.

## Data Availability

The complete data set and supplemental files are available at FigShare (DOI: 10.6084/m9.figshare.31079962).

