## Supplemental file 1 (2024 content) for "Does charging for corrections in the bioscience literature disincentivize pre-publication handling of problematic image data? An ImageTwin-AI study"

### Images dupl' from other journals

J. Cancer. 2024; 15(2): 343-355.
DOI: 10.7150/jca.86393
Figure 2C

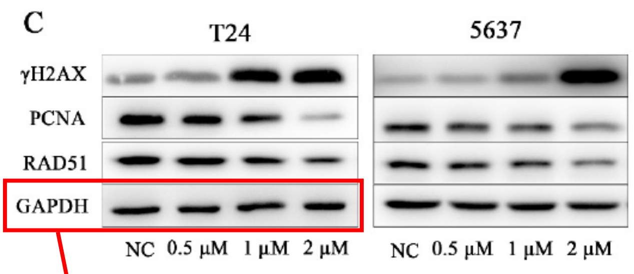


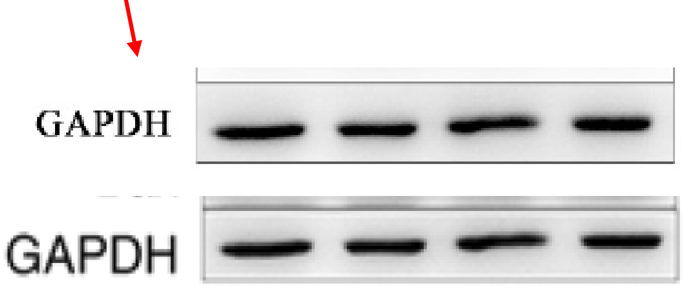


ONCOLOGY LETTERS 20: 169, 2020
DOI: 10.3892/ol.2020.12032
Figure 2F

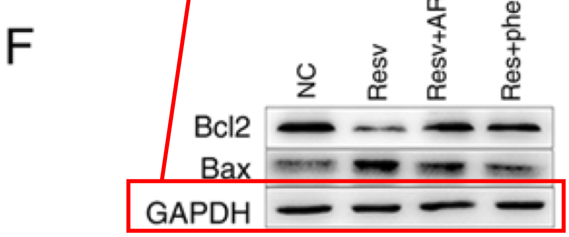

J. Cancer. 2024; 15(2): 356-369.
DOI: 10.7150/jca.83985
Figure 3A

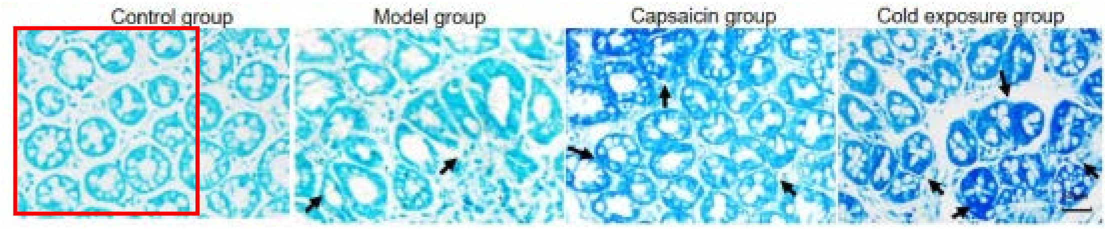


Evidence-Based Complementary and Alternative Medicine
Volume 2021, Article ID 5547578, 25 pages
DOI: 10.1155/2021/5547578
Figure 5

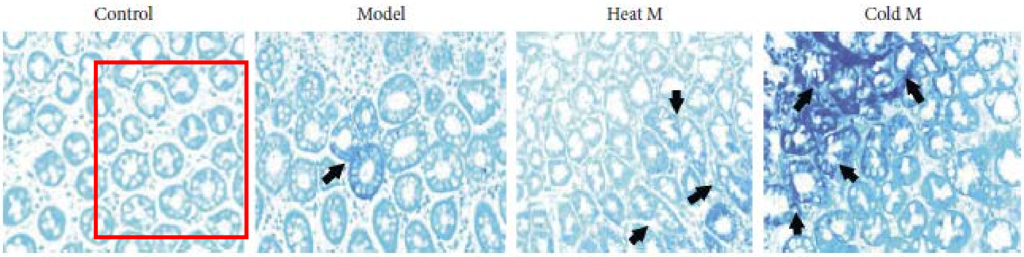

J. Cancer. 2024; 15(2): 356-369.
DOI: 10.7150/jca.83985
Figure 7A&B

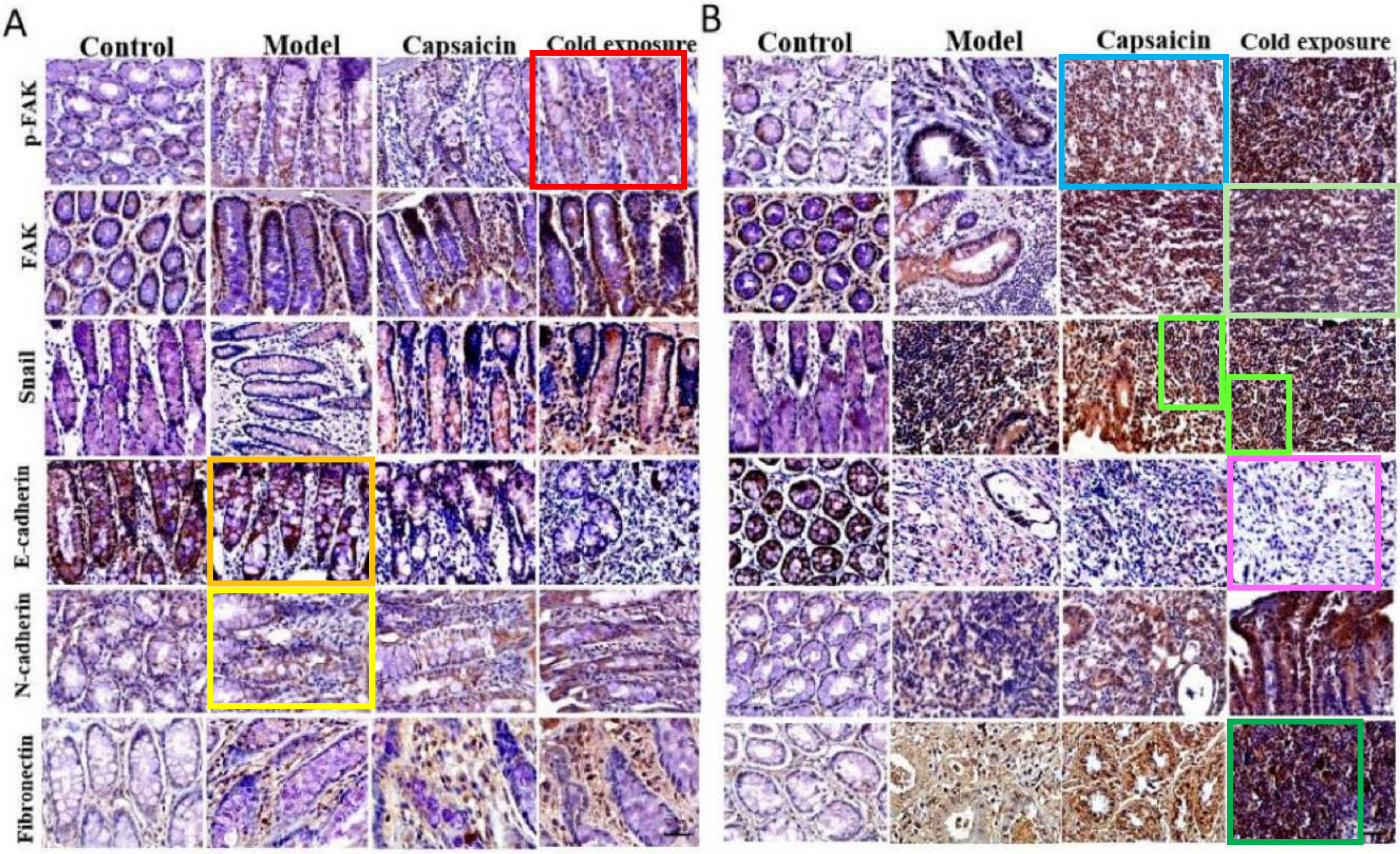


*World J Gastroenterol* 2022, 28(35): 5154-5174.
DOI: 10.3748/wjg.v28.i35.5154
Figure 5A

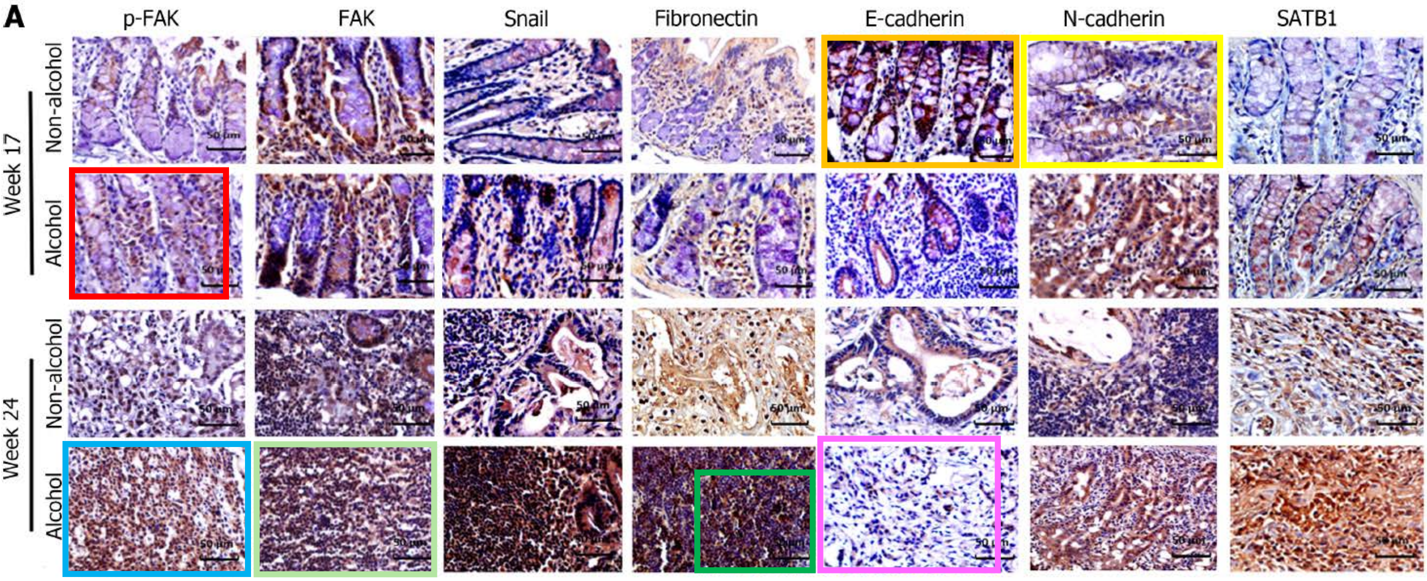

J. Cancer. 2024; 15(2): 356-369.
DOI: 10.7150/jca.83985
Figure 4 D&E vs. Figure 5A&B

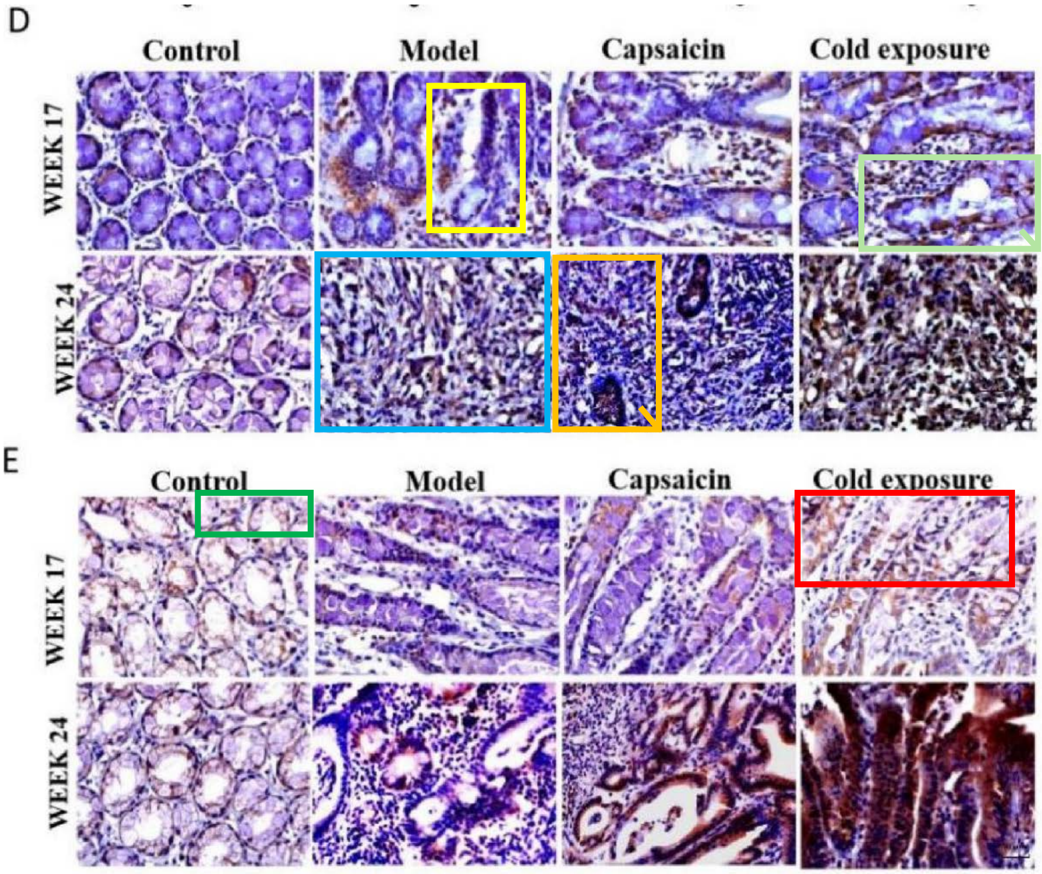


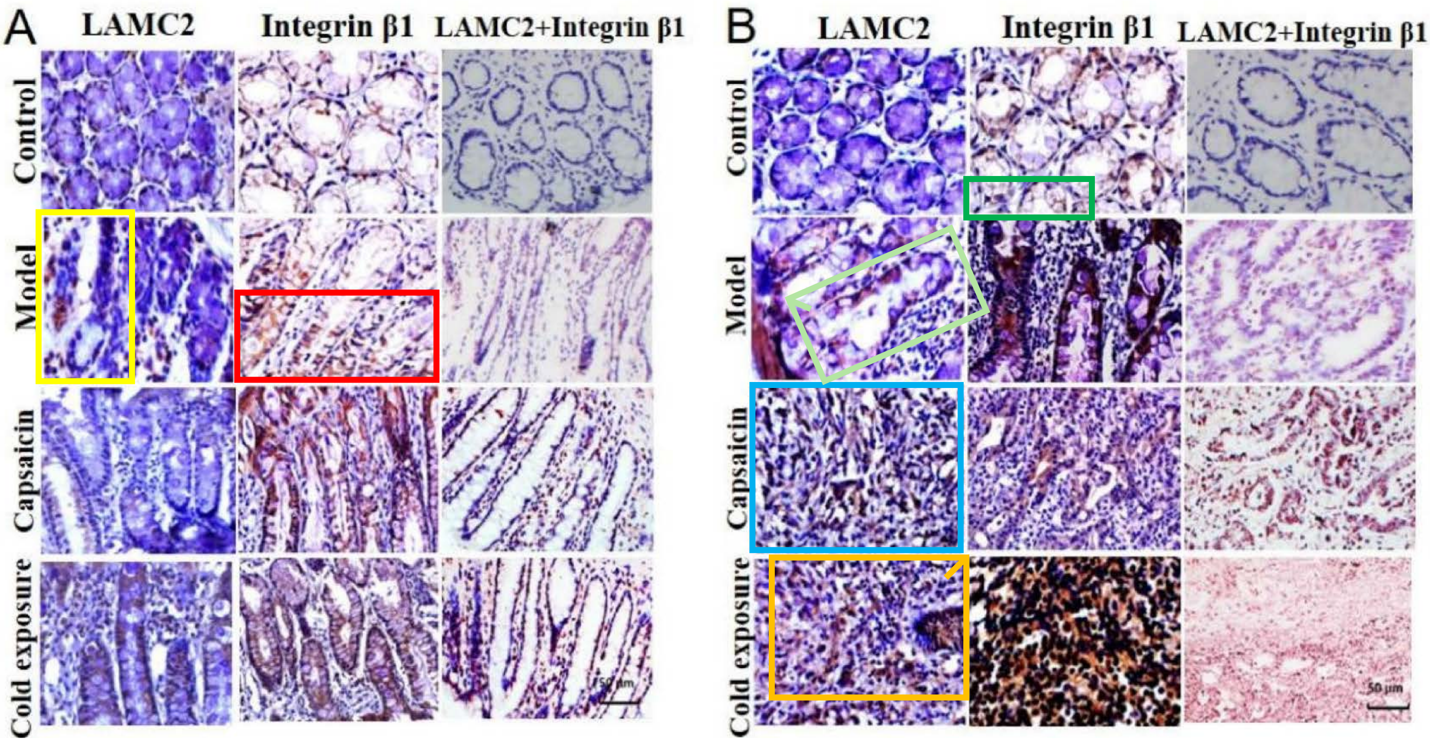

J. Cancer. 2024; 15(2): 356-369.
DOI: 10.7150/jca.83985
Figure 4B vs. 6A

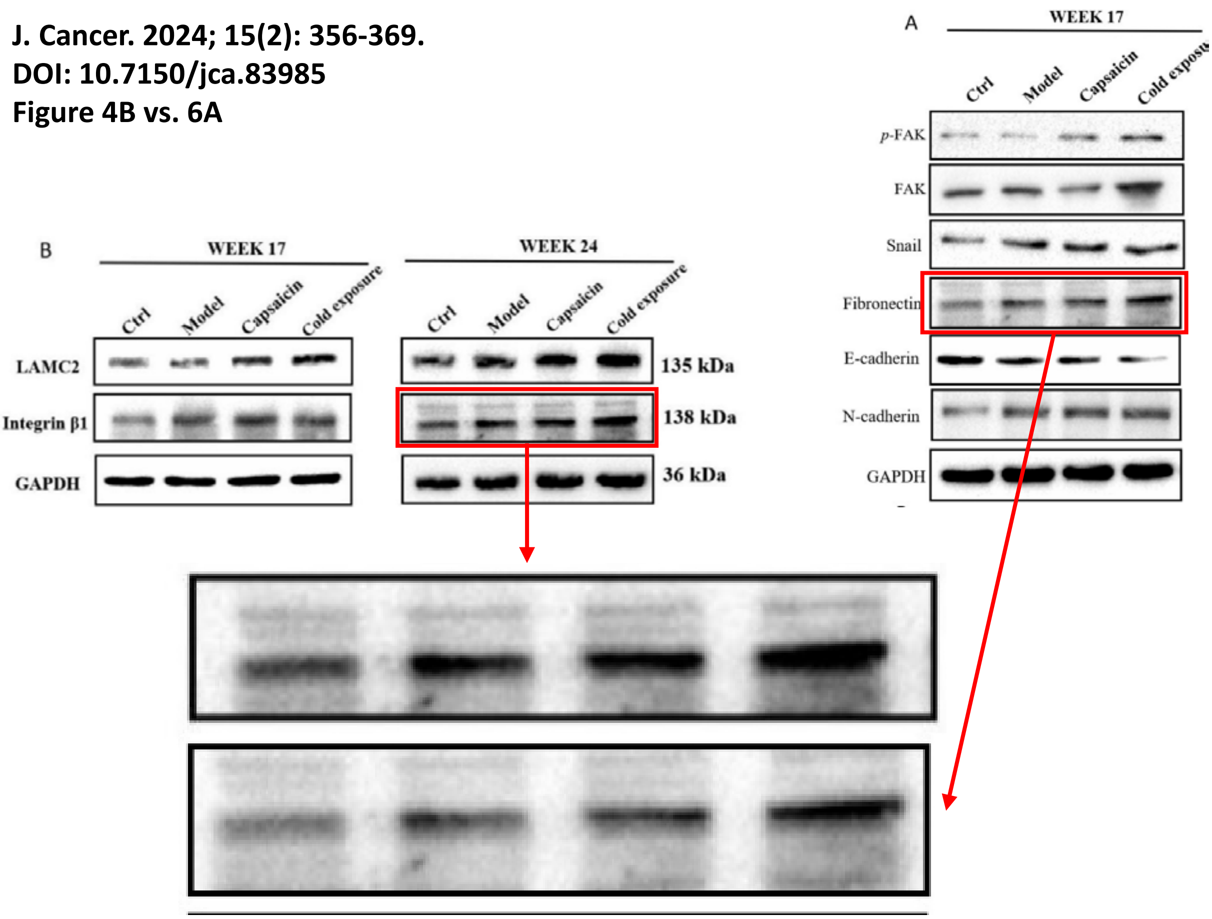

**J. Cancer. 2024; 15(4): 871-879.**
**doi: 10.7150/jca.90732**
**Figure 6**

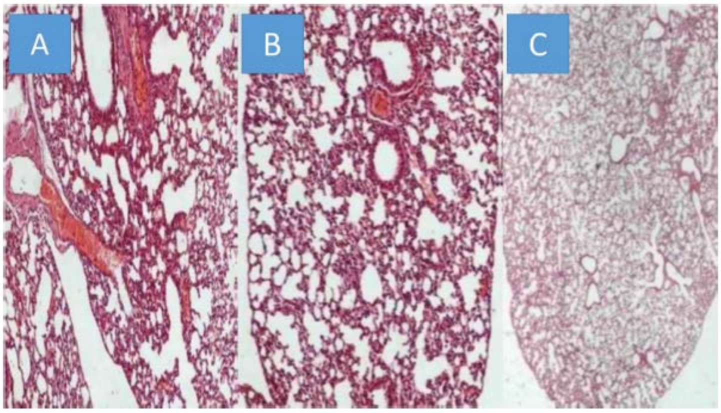

Figure 6. A) Paclitaxel group, B) Sotatercept group, and C) Iloprost group all x 200 magnification.

**J. Cancer. 2014; 5(5): 344-350.**
**doi: 10.7150/jca.9021**
**Figure 1**

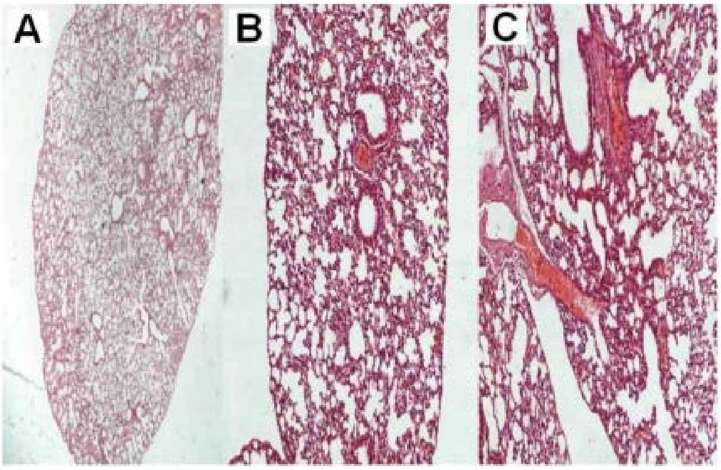

**J. Cancer. 2024; 15(4): 880-888.**
**doi: 10.7150/jca.90494**

**Figure 5**

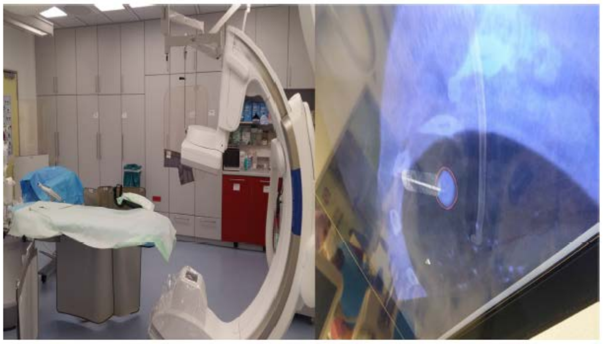

Figure 5. Cios Spin Siemens.

**Figure 8**

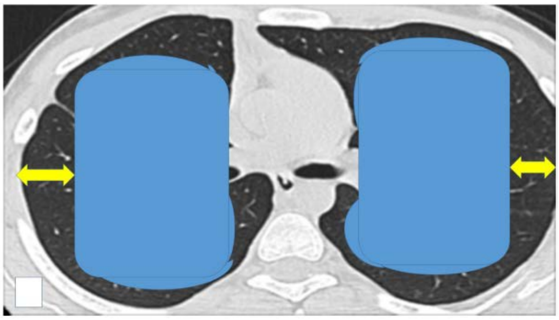

**J. Cancer. 2024; 15(4): 880-888.**
**doi: 10.7150/jca.90494**

**Figure 5**

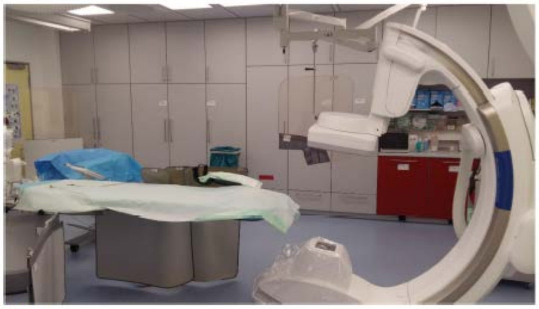

Figure 6. Dyna CT system Philips.

**Figure 7**

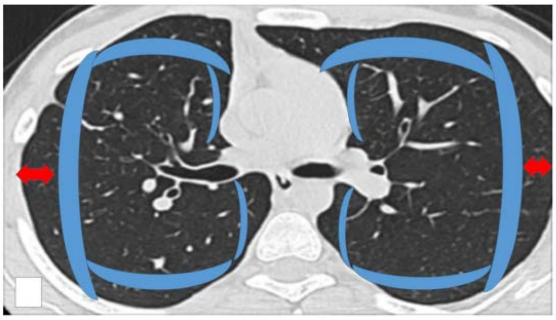

**J. Cancer. 2024; 15(4): 1115-1123.**
**doi: 10.7150/jca.91892**
**Figure 4A**

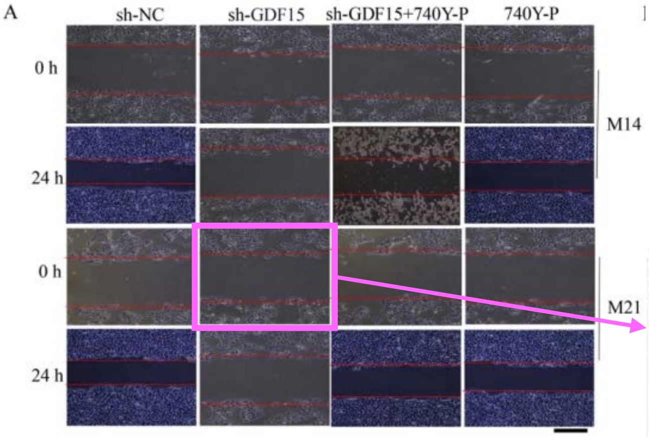


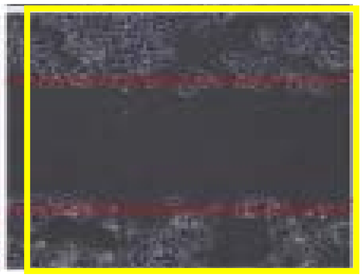

**BioMed Research International**
**Volume 2022, Article ID 2337447**
**DOI: 10.1155/2022/2337447**
**Figure 8A** ***RETRACTED***

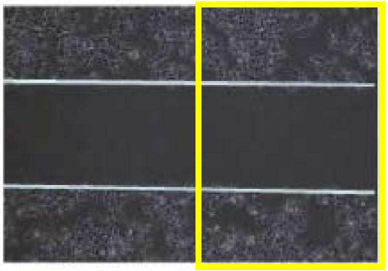

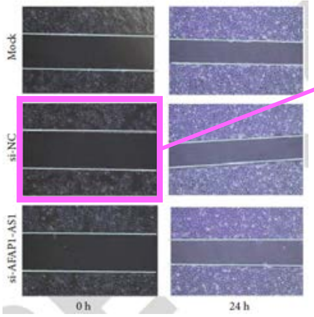

J. Cancer. 2024; 15(4): 1115-1123.
doi: 10.7150/jca.91892
Figure 3A

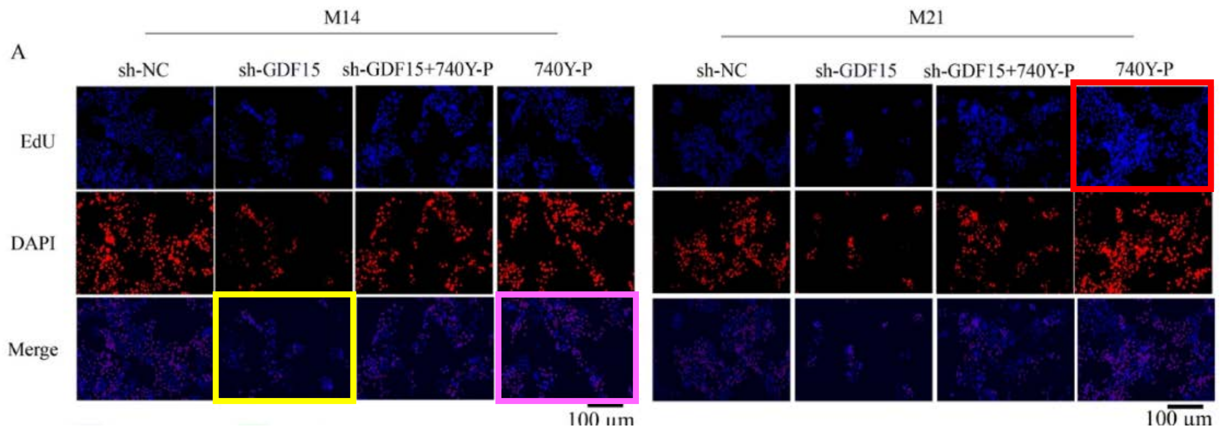


BioMed Research International
Volume 2022, Article ID 2337447
DOI: 10.1155/2022/2337447
Figure 6A *RETRACTED*

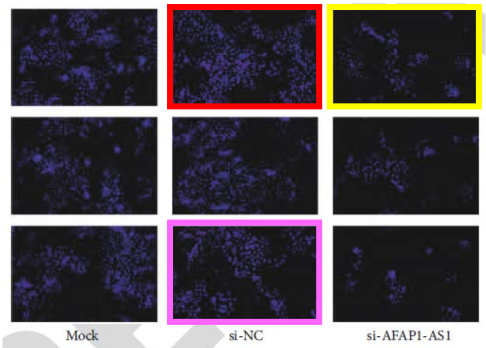

J. Cancer. 2024; 15(4): 1115-1123.
doi: 10.7150/jca.91892
Figure 4C

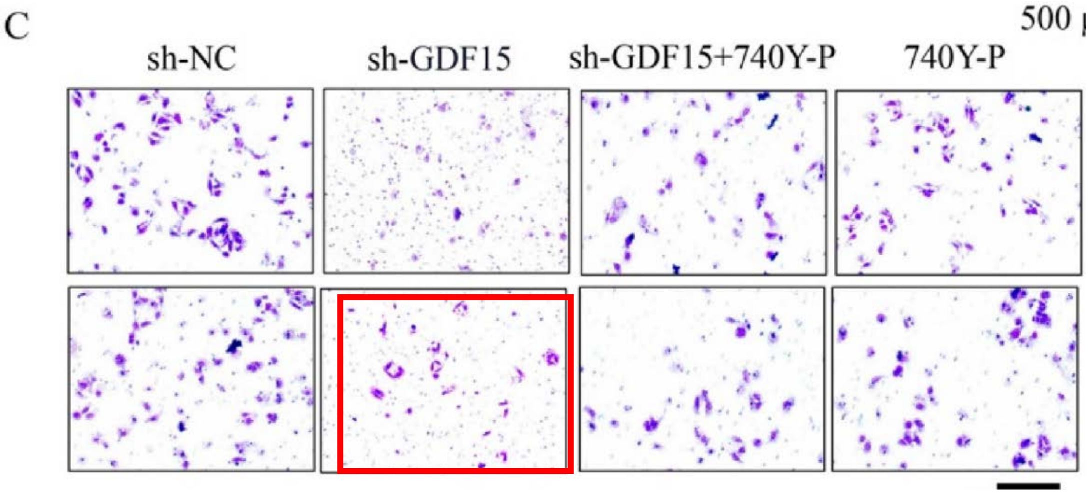


J. Cancer. 2021; 12(11): 3291-3298.
doi: 10.7150/jca.56218
Figure 2C

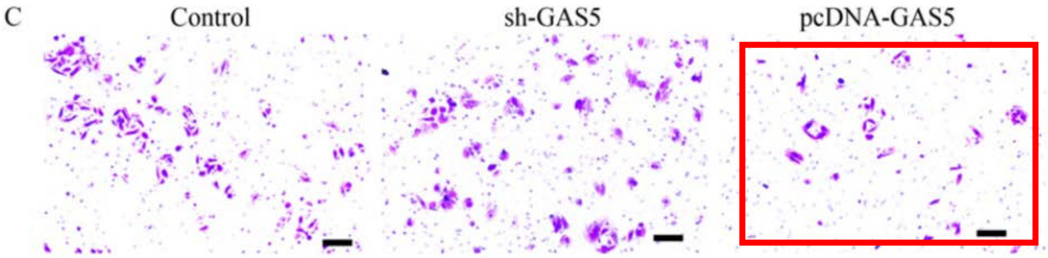


Open Medicine 2020; 15: 663-671
DOI: 10.1515/med-2020-0137
Figure 1C

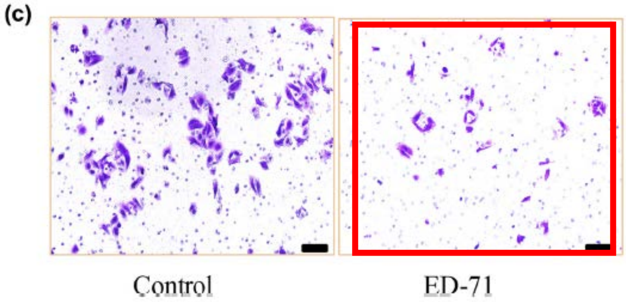

J. Cancer. 2024; 15(6): 1718-1730.
doi: 10.7150/jca.91184
Figure 2D

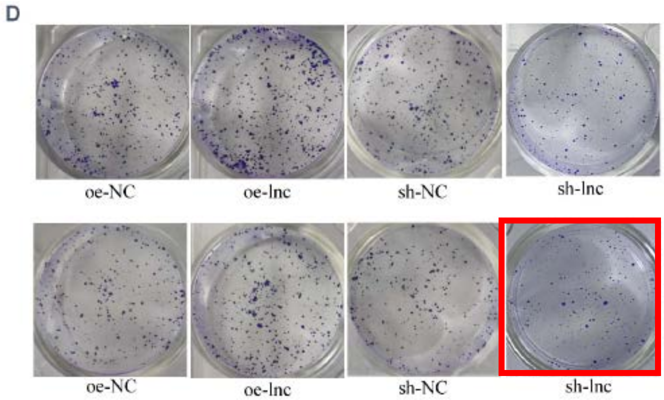


Journal of Oncology
Volume 2022, Article ID 4849174
DOI: 10.1155/2022/4849174
Figure 2A

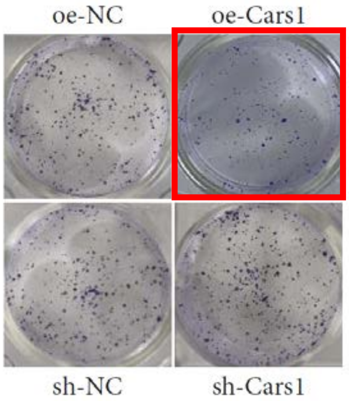

**J. Cancer. 2024; 15(7): 2074-2094.**
**DOI: 10.7150/jca.93398**
**Figure 7A**

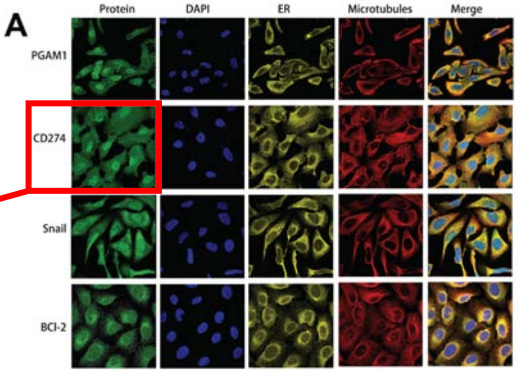


CD274

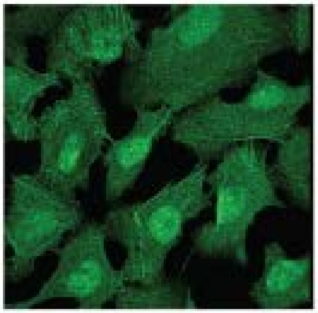

PD-L1

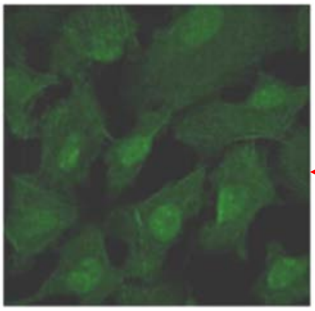

**Cells 2022, 11, 3958.**
**DOI: 10.3390/cells11243958**
**Figure 4E**

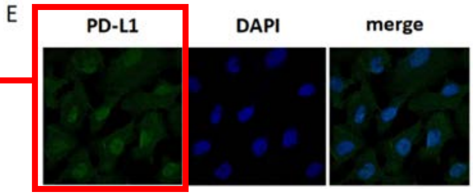

J. Cancer. 2024; 15(10): 2981-2993.
DOI: 10.7150/jca.89970
Figure 2B

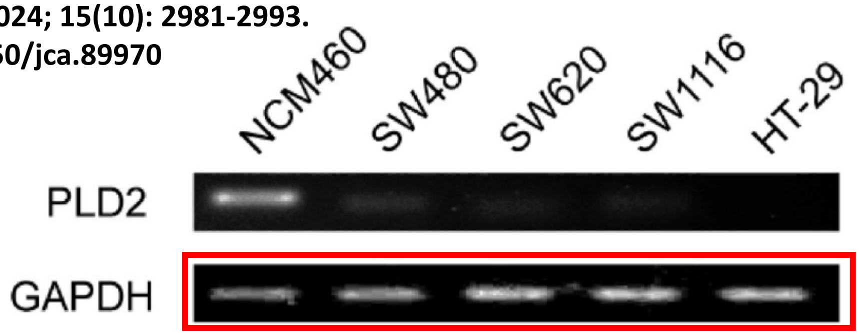


Transl. Oncol. 2016, 9(6): 512-520
DOI: 10.1016/j.tranon.2016.09.009
Figure 1B

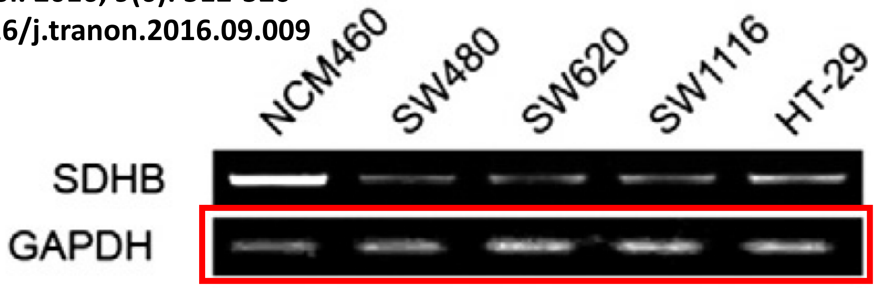

J. Cancer. 2024; 15(10): 3114-3127.
DOI: 10.7150/jca.93833

**Figure 1D**

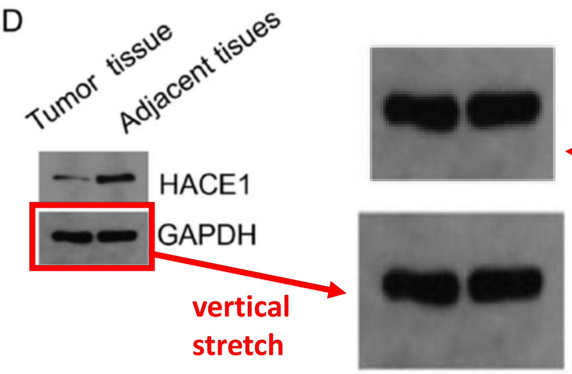


**Figure 4J**

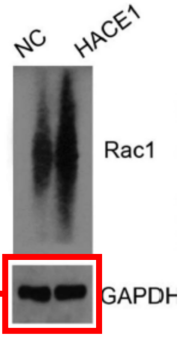


**Figure 6D**

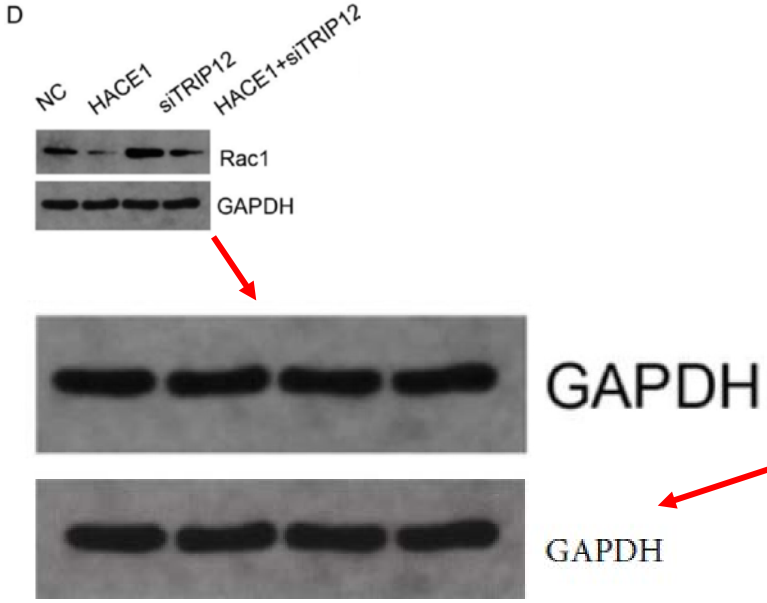


Mediators of Inflammation
Volume 2023, Article ID 8052579
DOI: 10.1155/2023/8052579

**Figure 8C**

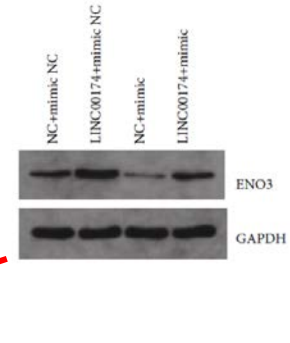

**J. Cancer. 2024; 15(12): 3995-4006.**
**DOI: 10.7150/jca.90087**
**Figure 3C**

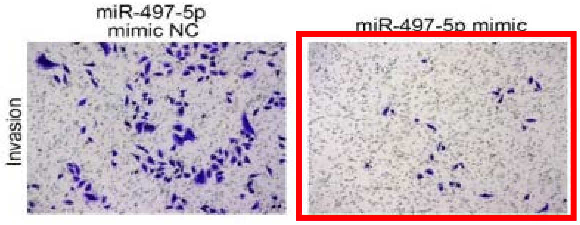


**Figure 5E**

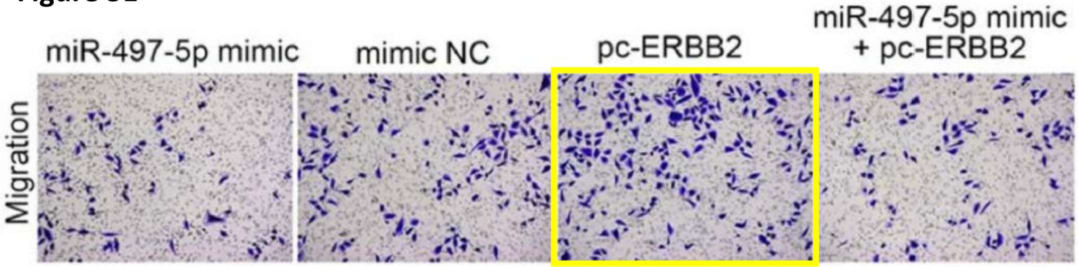


**Journal of Oncology**
**Volume 2022, Article ID 3966386.**
**DOI: 10.1155/2022/3966386**
**Figure 2B**

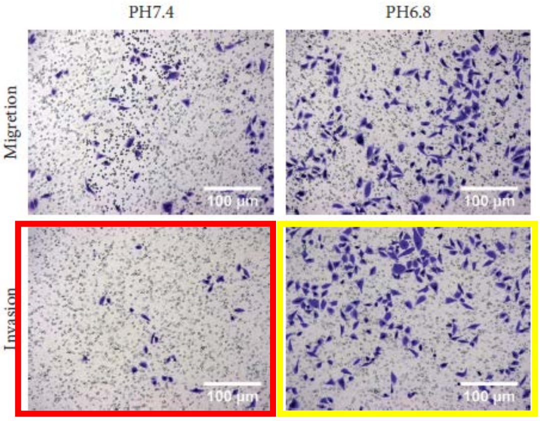

J. Cancer. 2024; 15(13): 4374-4385.
DOI: 10.7150/jca.95876
Figure 2D

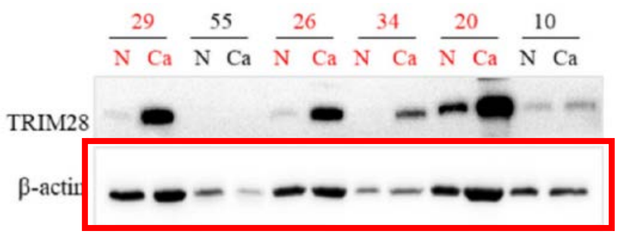


Front. Oncol. 2023 13:1158087.
DOI: 10.3389/fonc.2023.1158087
Figure 1A

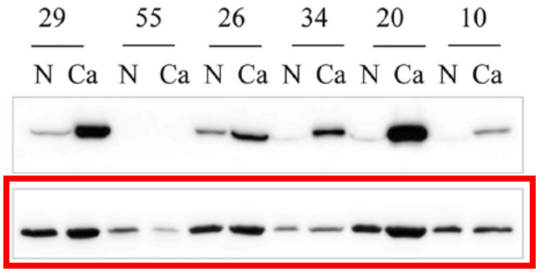

J. Cancer. 2024; 15(17): 5719-5728.
DOI: 10.7150/jca.99307

Figure 2A

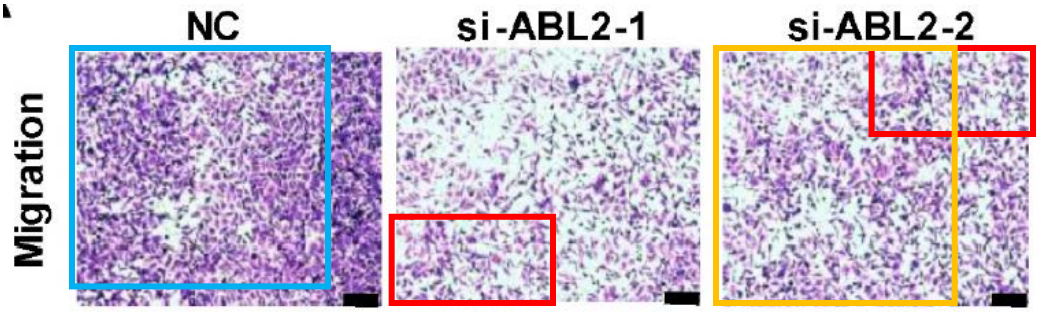


Figure 5B

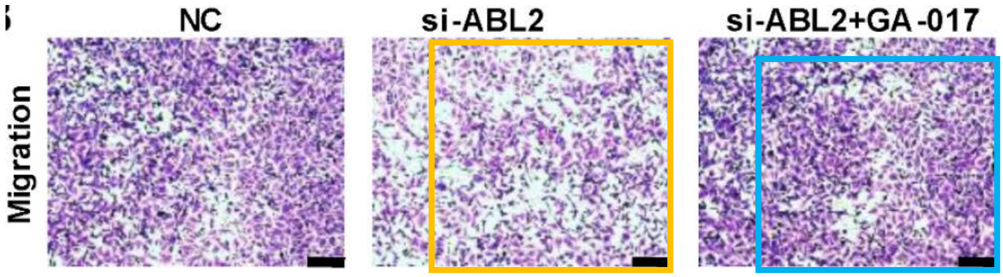


Figure 3D

Biology Direct (2023) 18:51
DOI: 10.1186/s13062-023-00402-9
Figure 2F

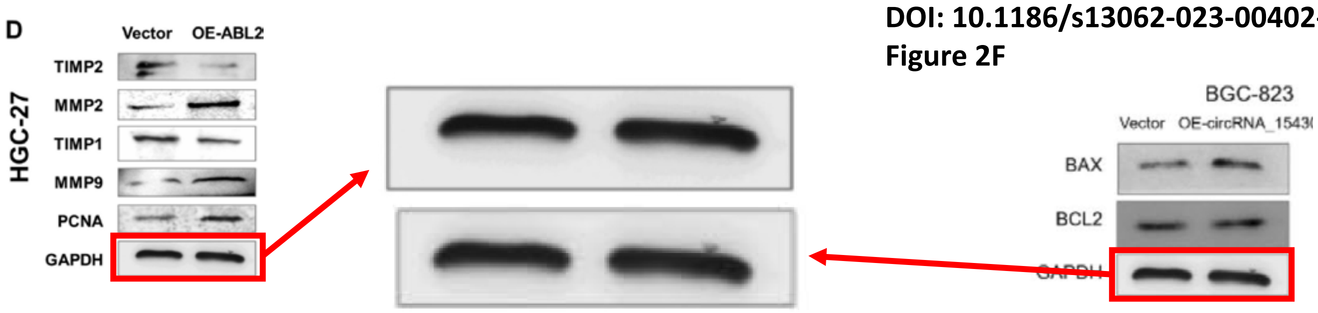

J. Cancer. 2024; 15(19): 6160-6176.
DOI: 10.7150/jca.894292
Figure 13-2

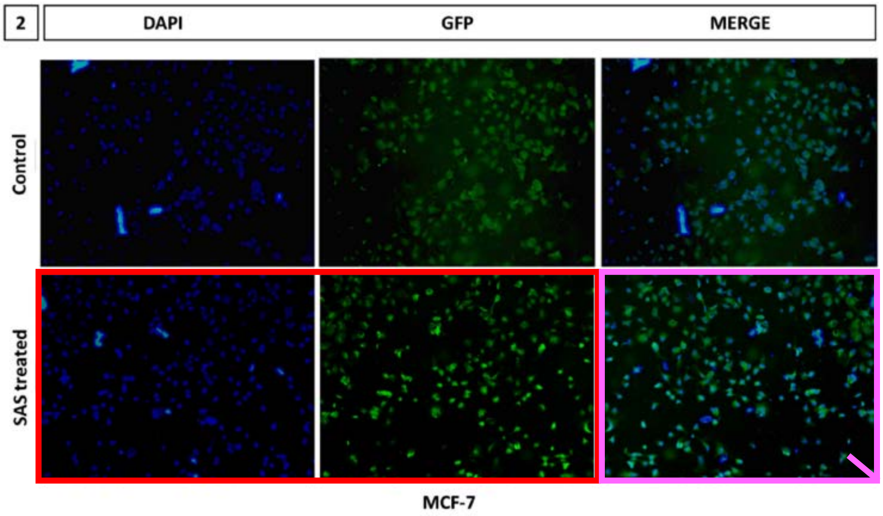


Front. Pharmacol. 2023, 14:1134758.
DOI: 10.3389/fphar.2023.1134758
Figure 11

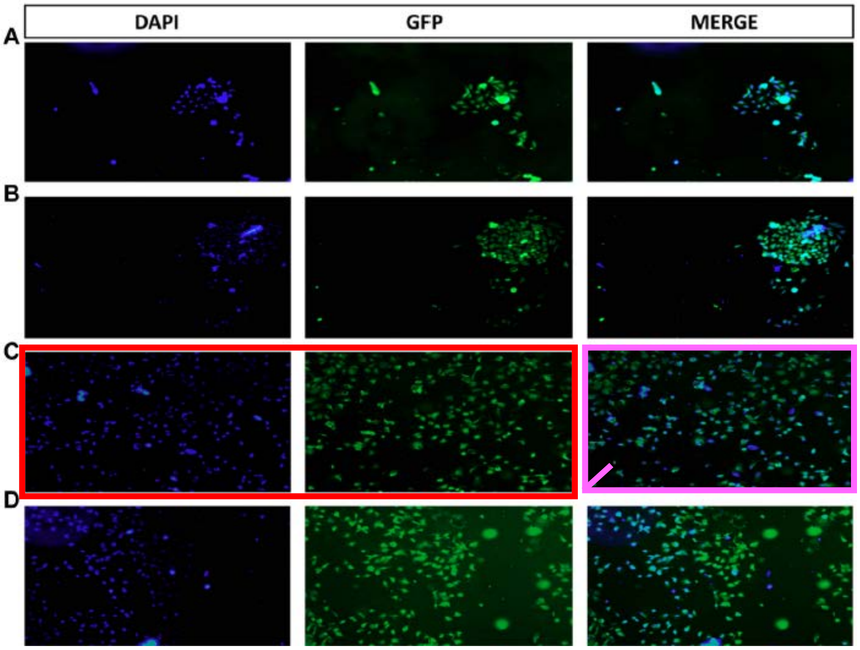

J. Cancer. 2024; 15(19): 6345-6354.
DOI: 10.7150/jca.100840
Figure 2

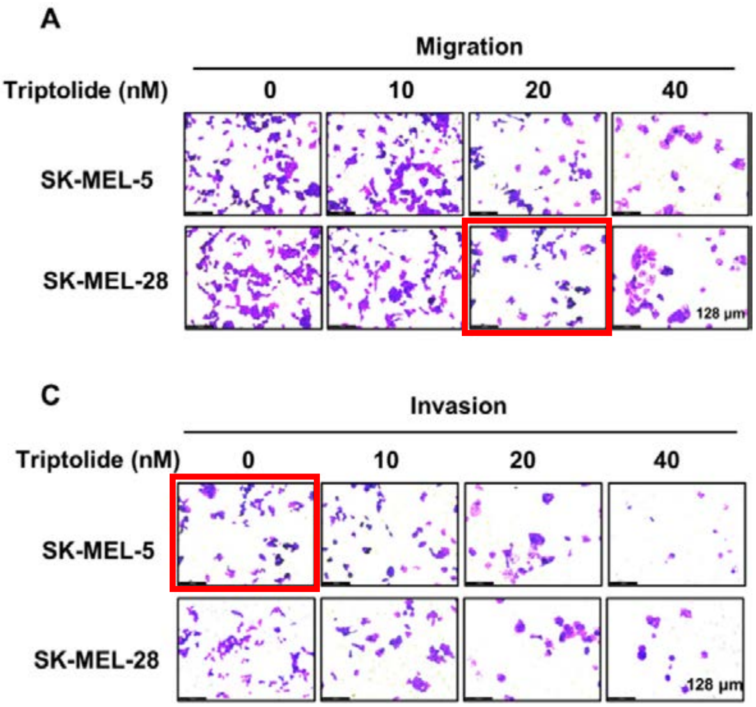


Figure 5C

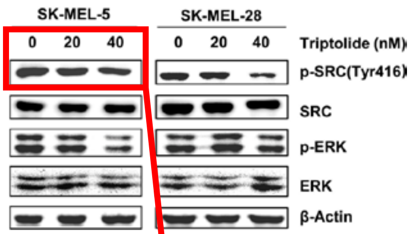


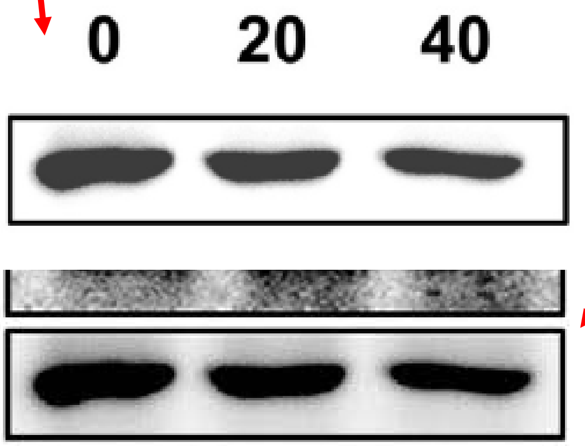


J. Ginseng Res. 46 (2022) 396e407
DOI: 10.1016/j.jgr.2021.07.004
Figure 4C

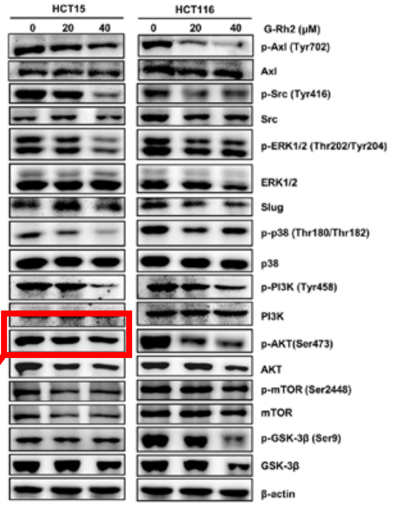

### Image problems within *J. Cancer* papers

J. Cancer. 2024; 15(1): 232-250.
doi: 10.7150/jca.87733
Figure 10B

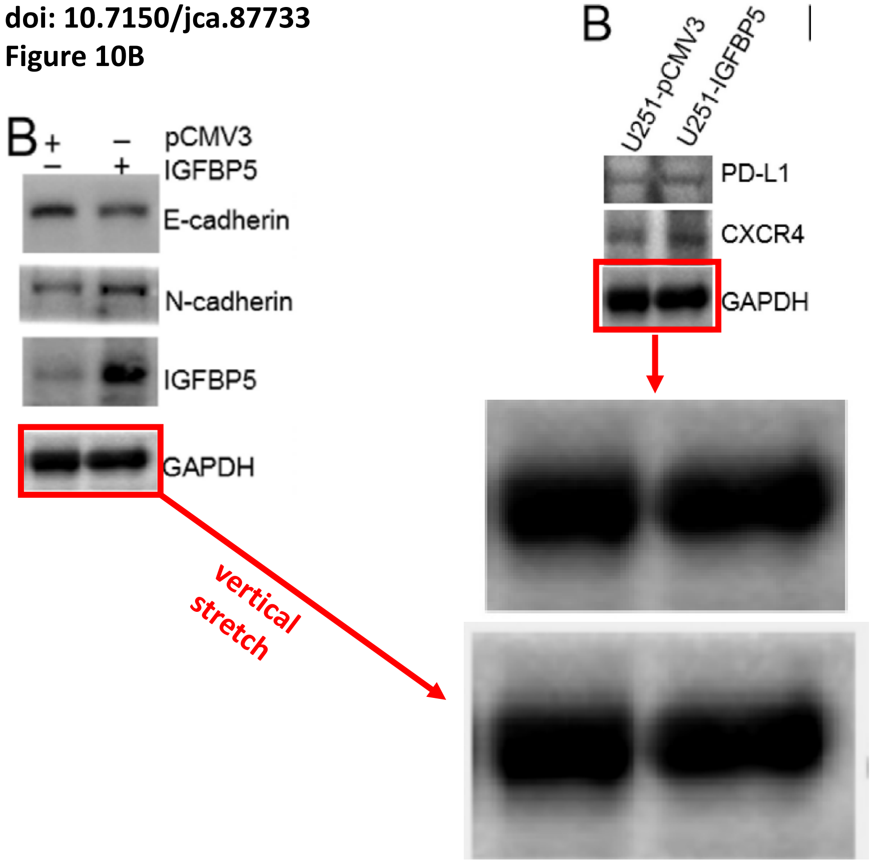

J. Cancer. 2024; 15(1): 251-274.
doi: 10.7150/jca.88397
Figure 13C

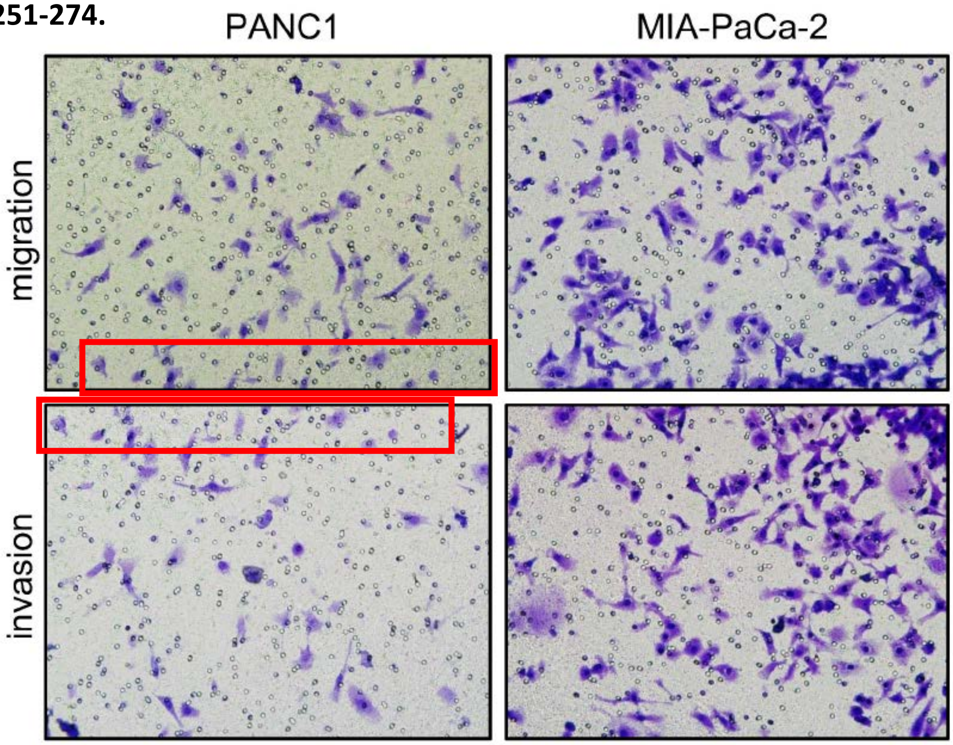

**J. Cancer. 2024; 15(2): 309-316.**
**doi: 10.7150/jca.89059**
**Figure 3B vs. 3D**

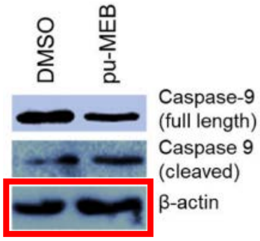


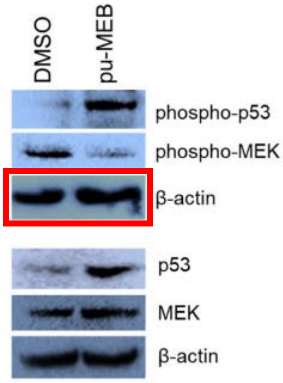

J. Cancer. 2024; 15(2): 370-382.
doi: 10.7150/jca.83856

Figure 3F

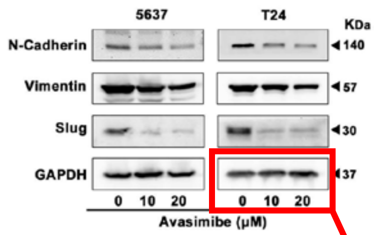


Figure 4F

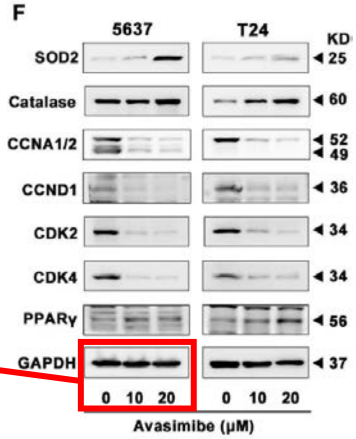


Rotate 180°, resize, adjust brightness/contrast

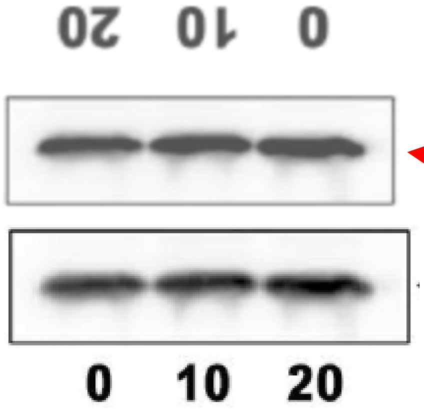

**J. Cancer. 2024; 15(2): 494-507.**
**doi: 10.7150/jca.89986**
**Figure 4A**

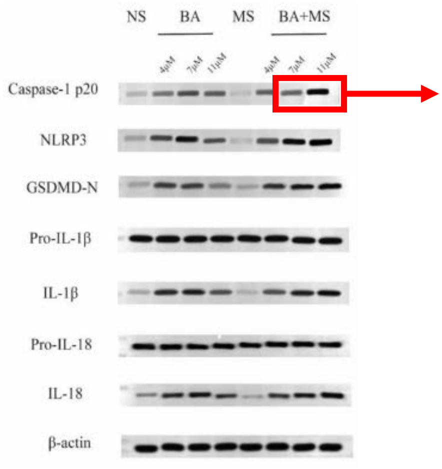


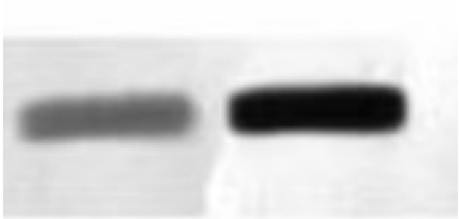

J. Cancer. 2024; 15(2): 545-559.
doi: 10.7150/jca.90236
Figure 1A

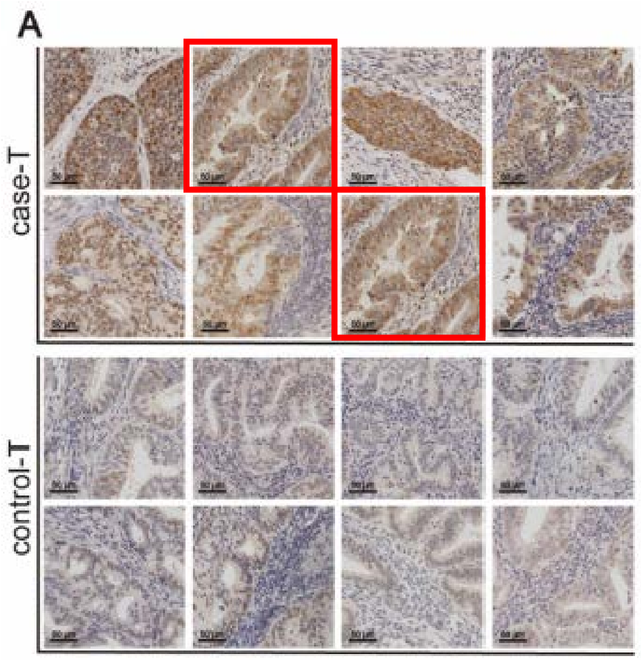

J. Cancer 2024; 15(3): 659-670.
doi: 10.7150/jca.88555

Figure 1D

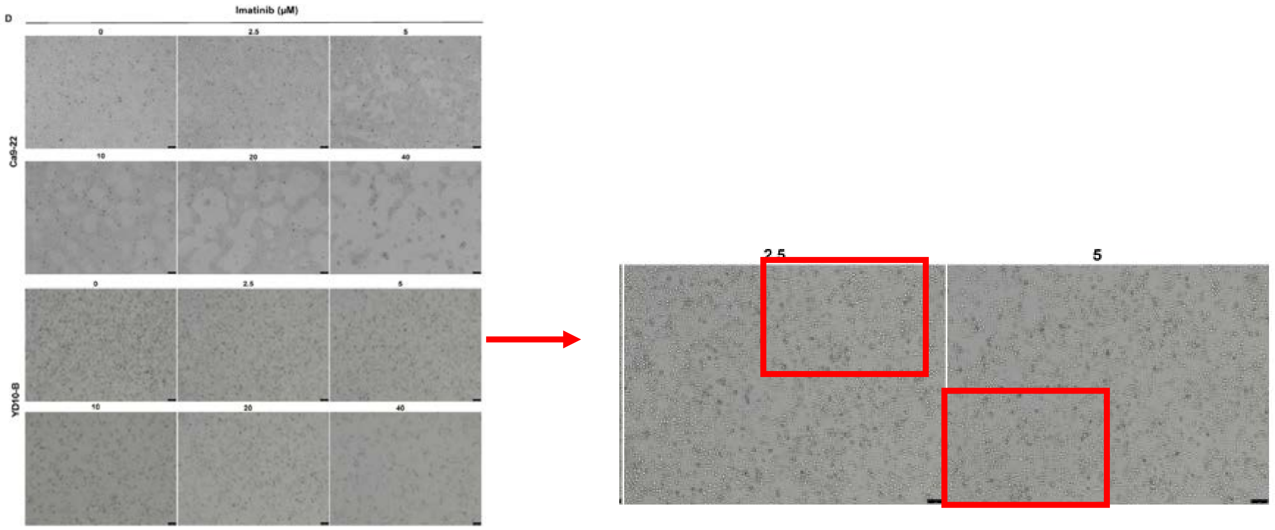


Figure 2A

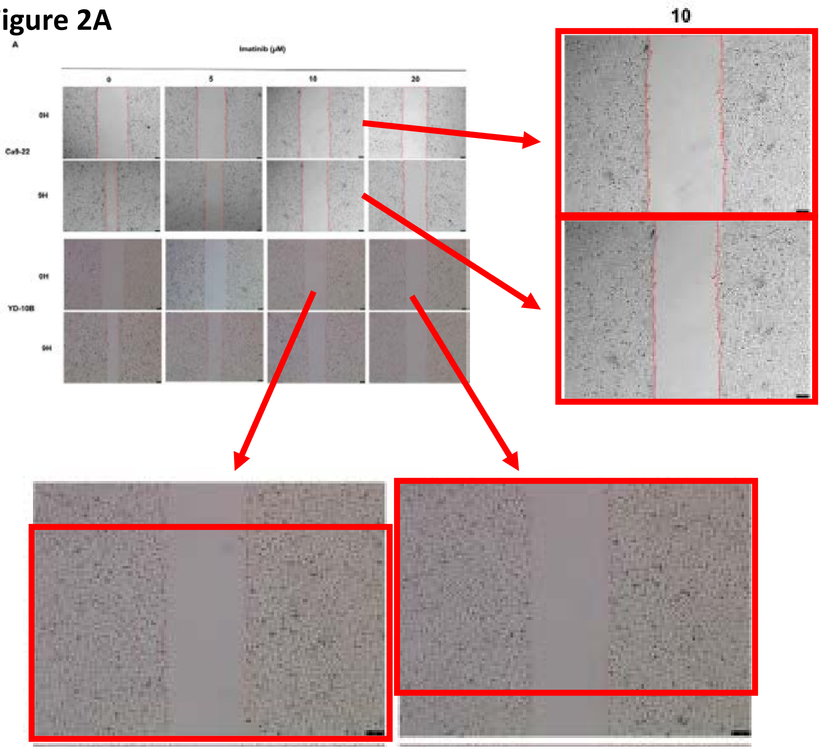

J. Cancer 2024; 15(3): 685-698
doi: 10.7150/jca.90574
Figure 2B/C

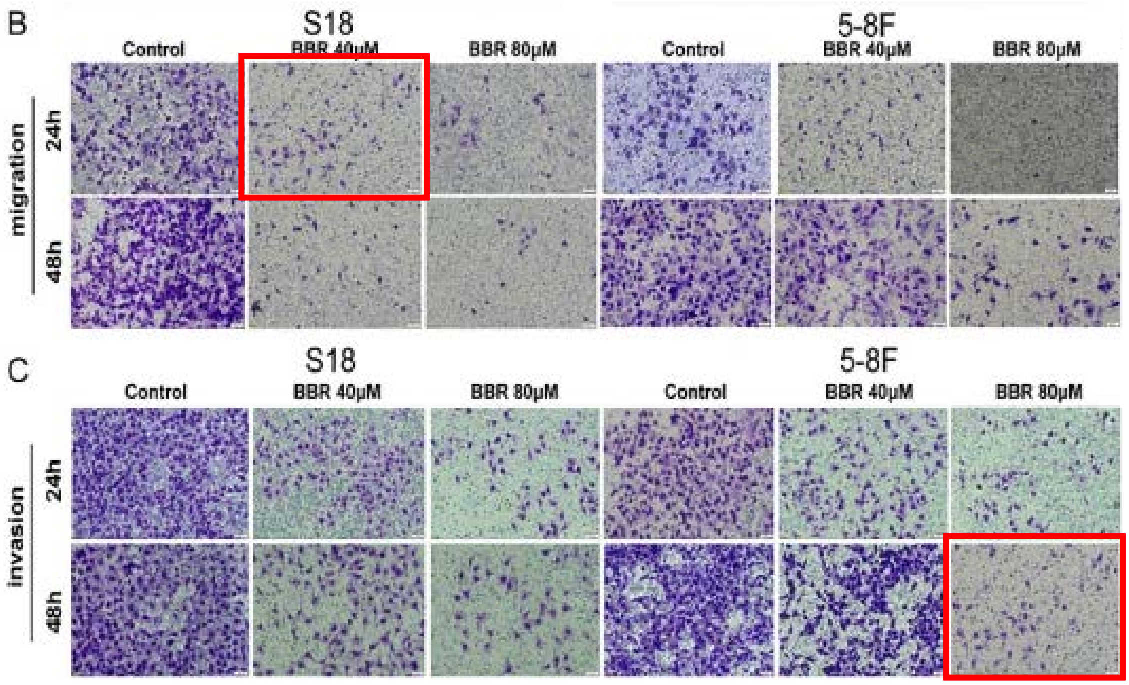

J. Cancer 2024; 15(3): 699-713
doi: 10.7150/jca.90371

Figure 3A

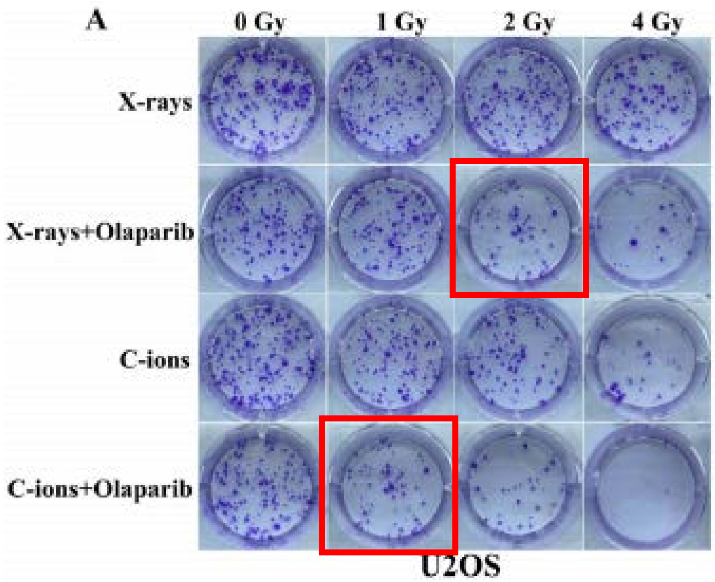


Figure 11A/B

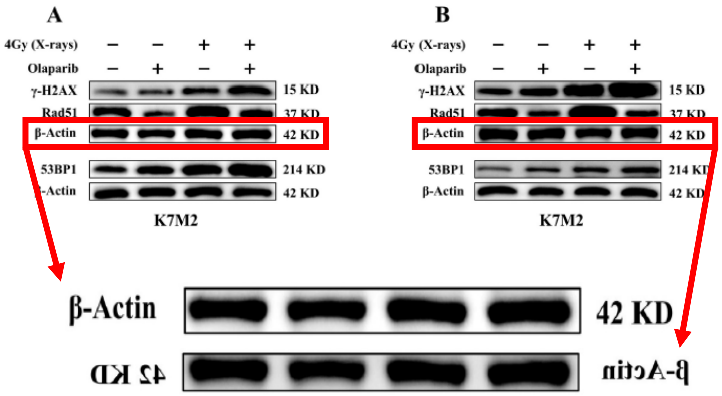

J. Cancer 2024; 15(3): 764-775
doi: 10.7150/jca.90051
Figure 2E/F

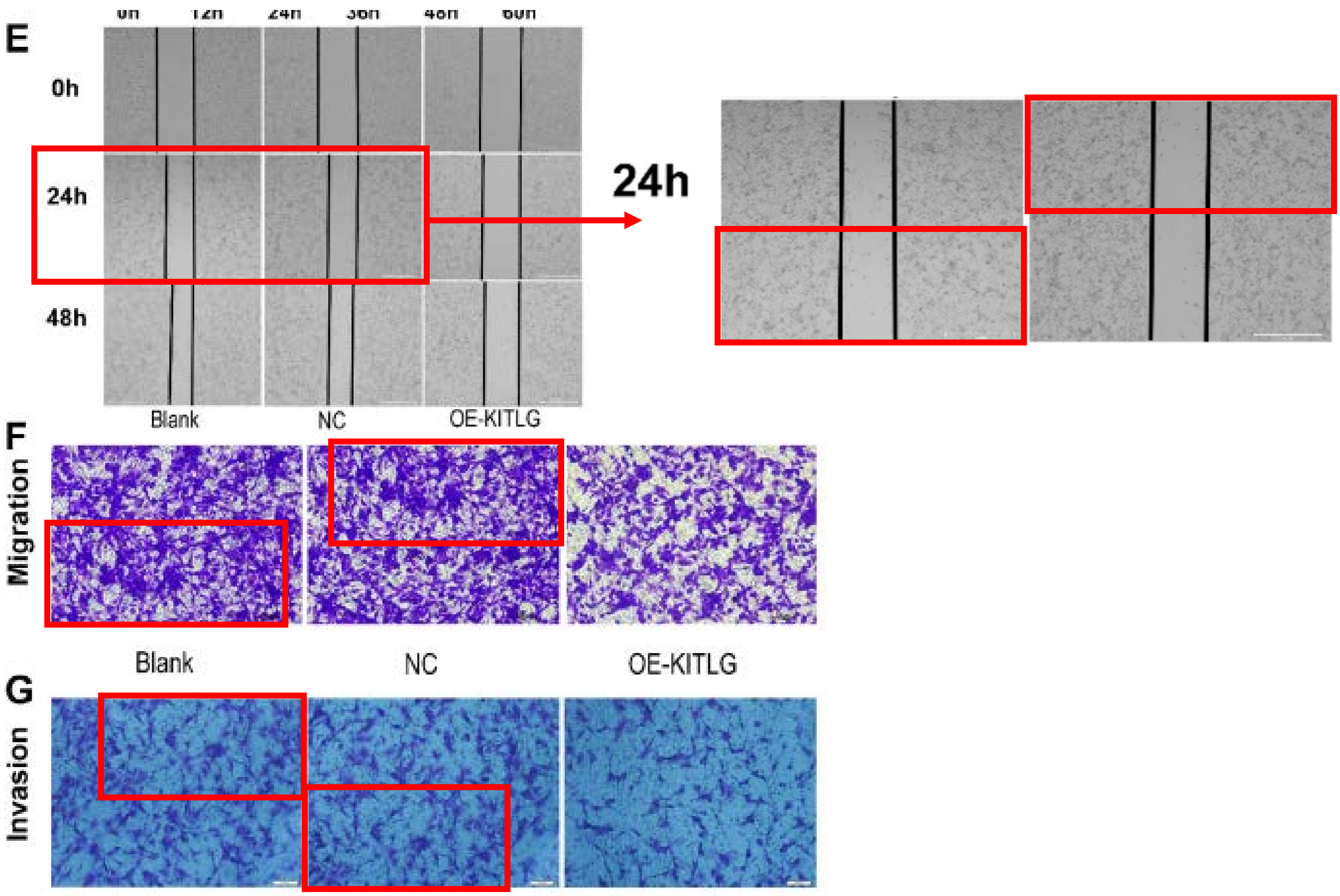

J. Cancer 2024; 15(4): 990-998
doi: 10.7150/jca.90321
Figure 4F

Blots on the right are just over-exposures of blots on the left

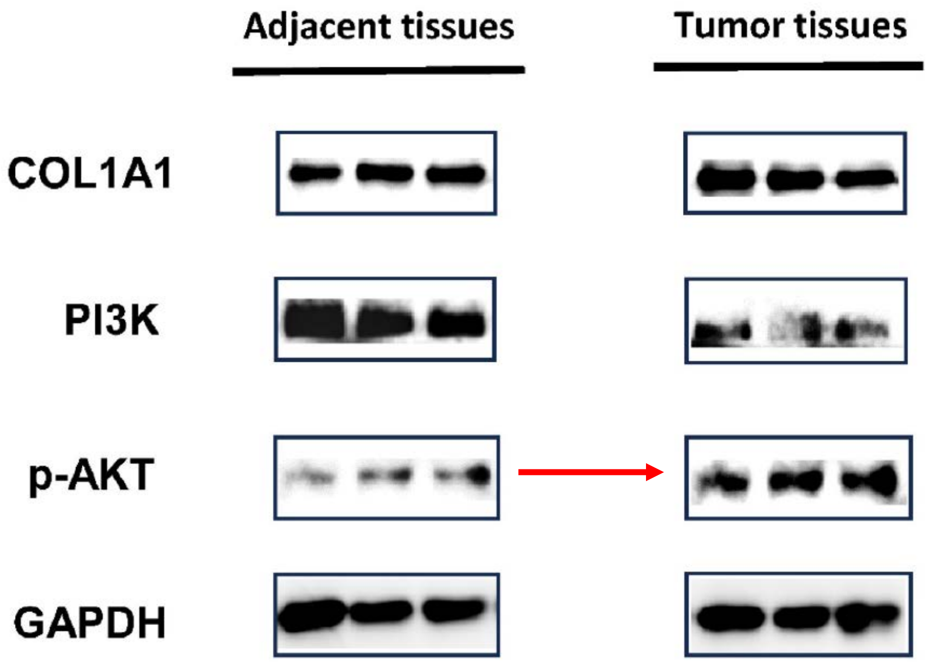

J. Cancer 2024; 15(4): 999-1008
doi: 10.7150/jca.90756

Figure 2C (overexpression of miR-135a)

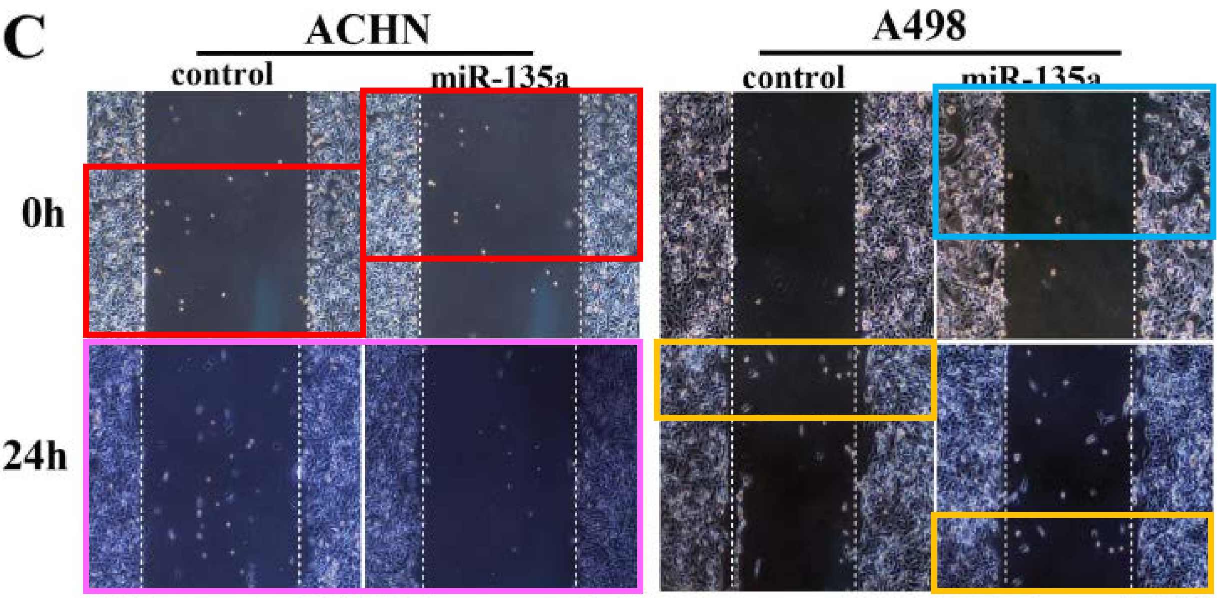


Figure 3C (inhibition of miR-135a)

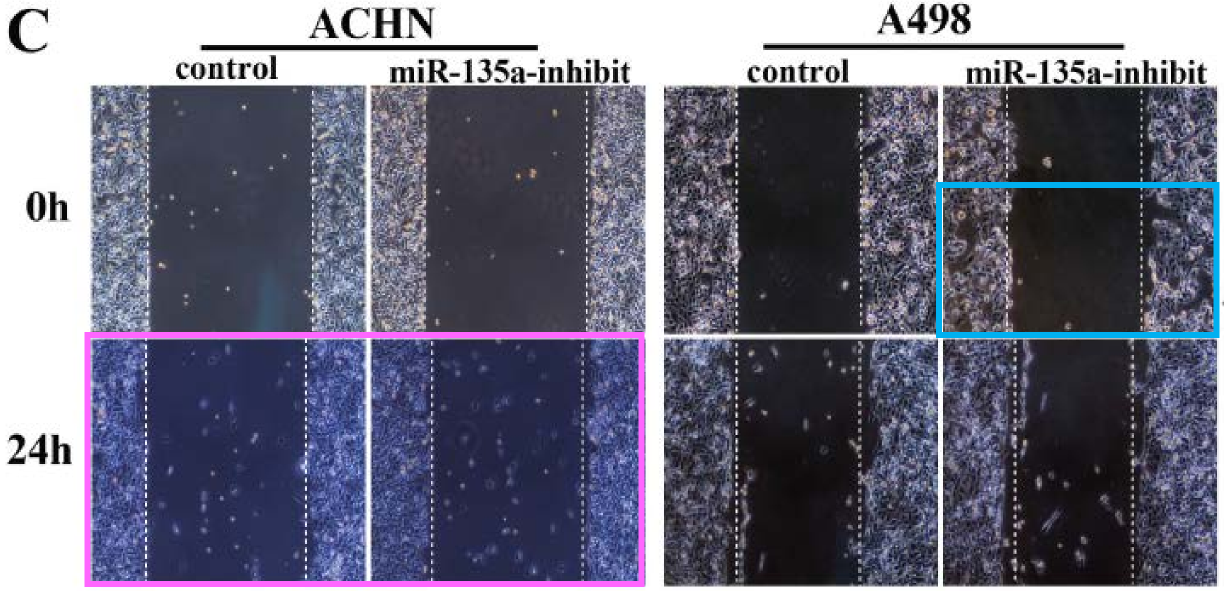

J. Cancer 2024; 15(4): 999-1008
doi: 10.7150/jca.90756
Figure 5

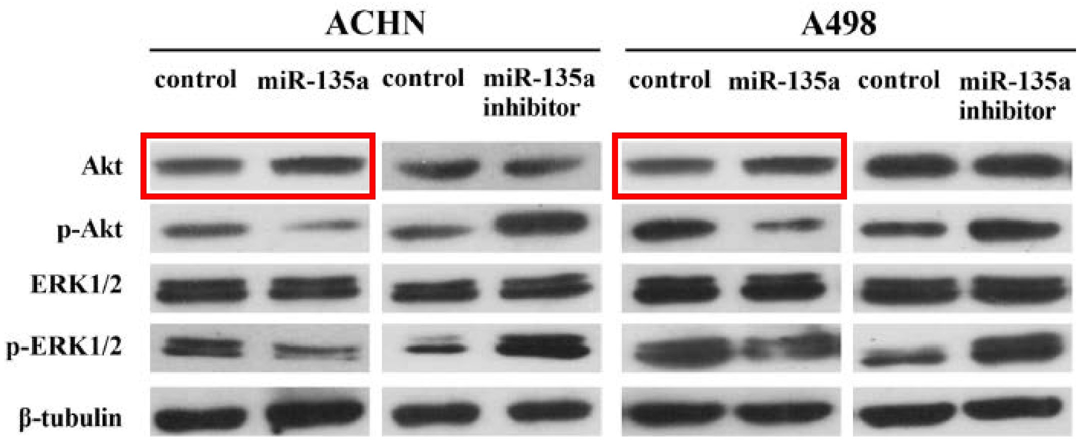

J. Cancer 2024; 15(4): 1124-1137
doi: 10.7150/jca.91360
Figure 5B/C

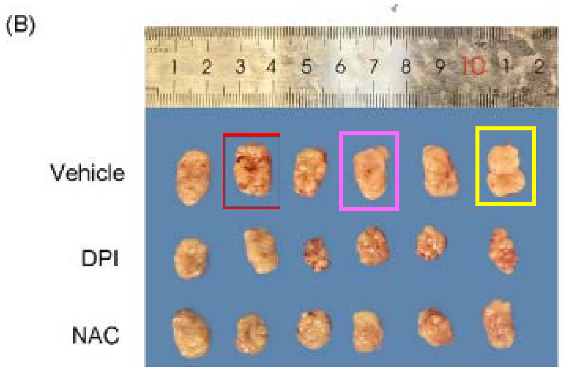


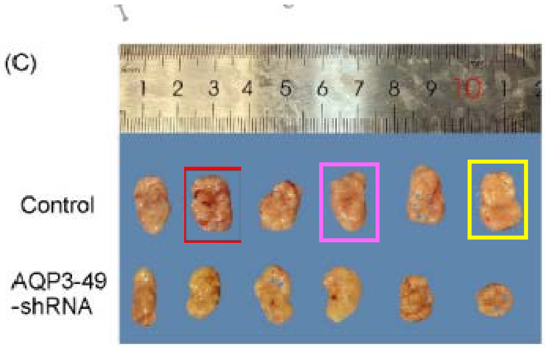


Also note "holey" appearance of many samples (blue background visible in holes)

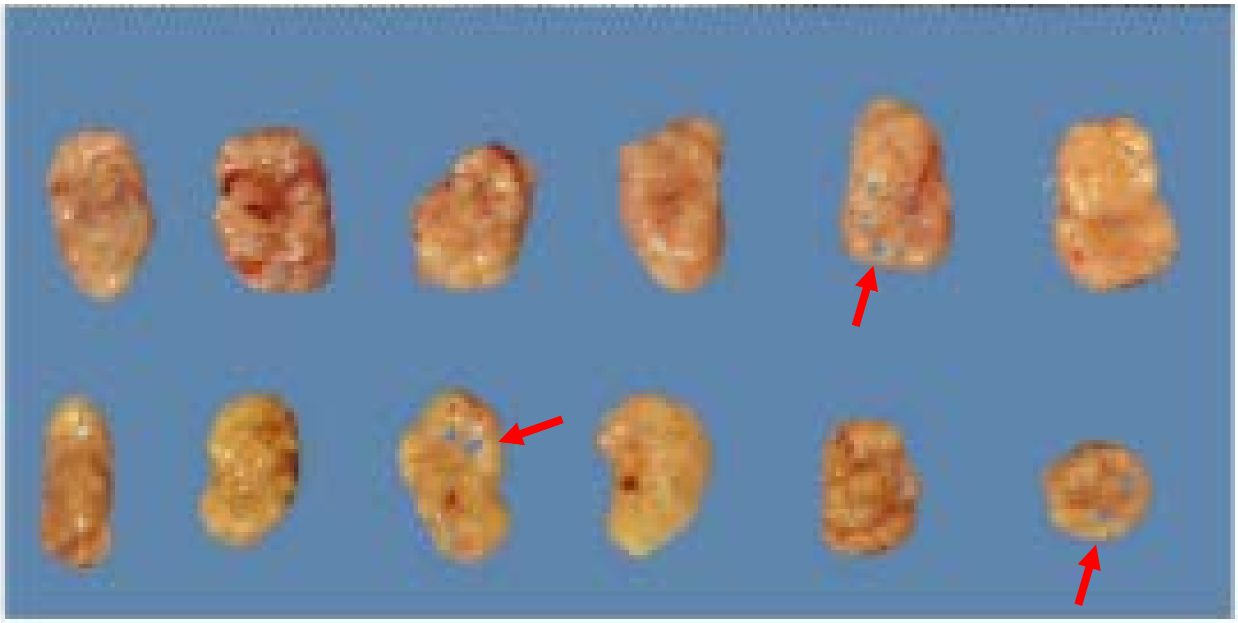

J. Cancer 2024; 15(5): 1138-1152
doi: 10.7150/jca.90422
Figures 1, 5A/B, 7B

Same GAPDH "loading control" used for 14 separate western blots

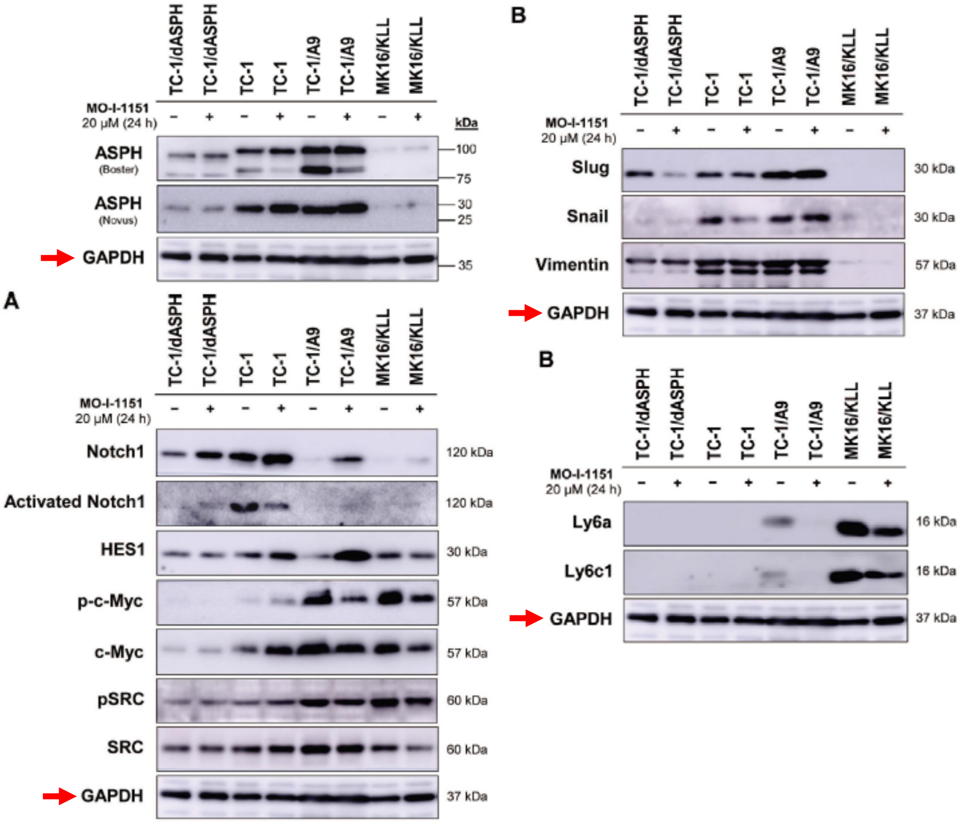

J. Cancer 2024; 15(5): 1257-1270
doi: 10.7150/jca.89485

Figure 2D/J

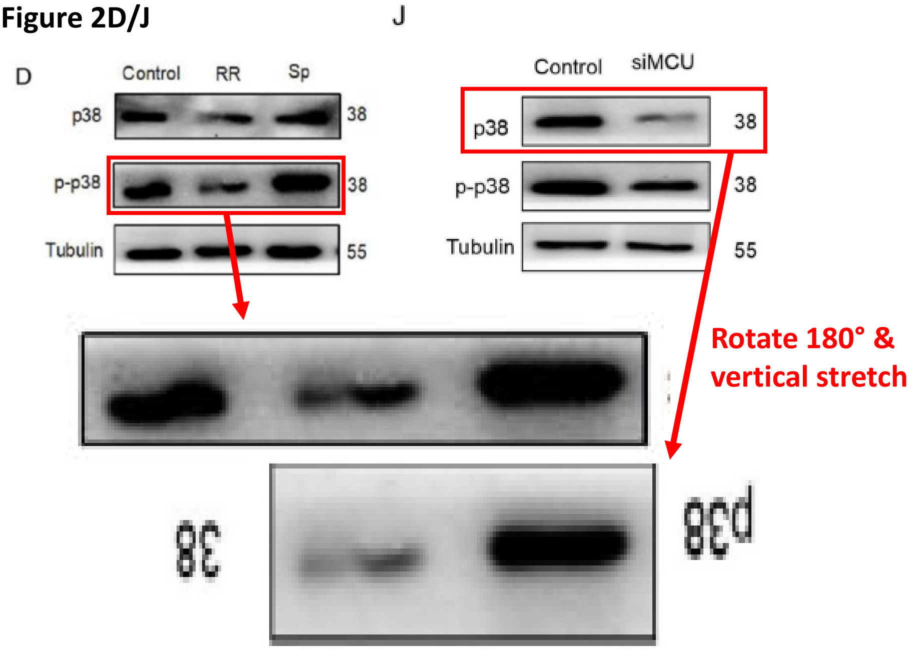


Figure 3A/J

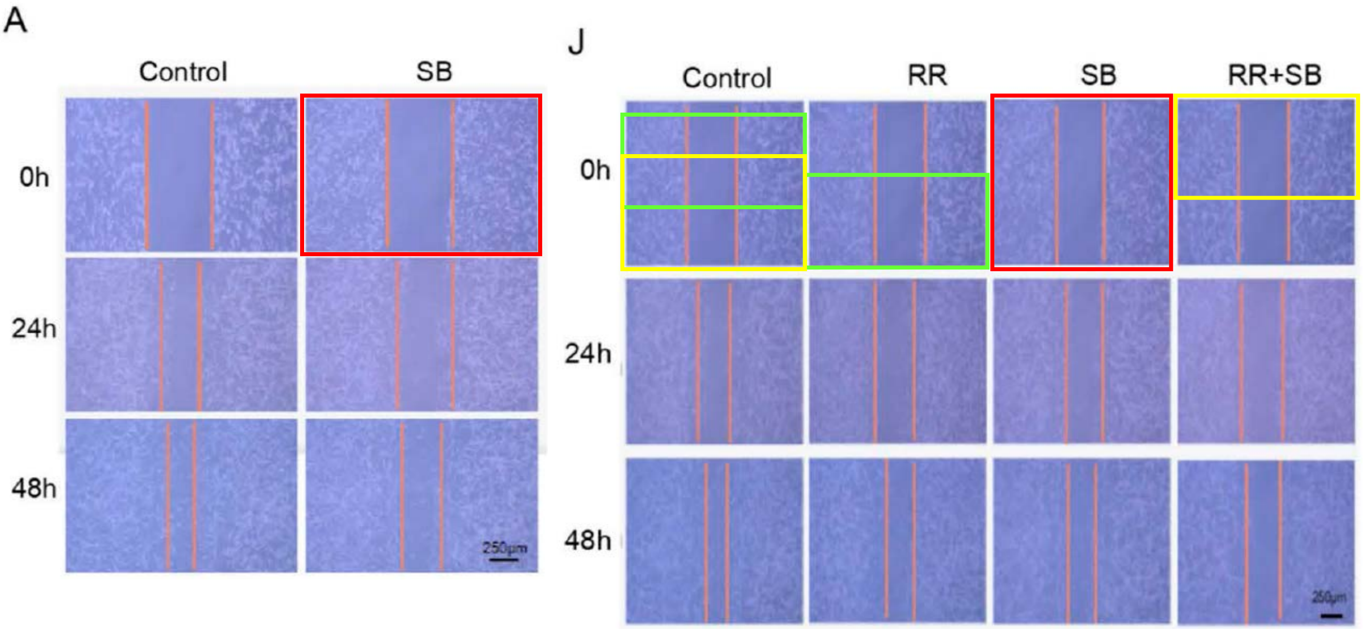

J. Cancer 2024; 15(5): 1257-1270
doi: 10.7150/jca.89485
Figure 4M/O

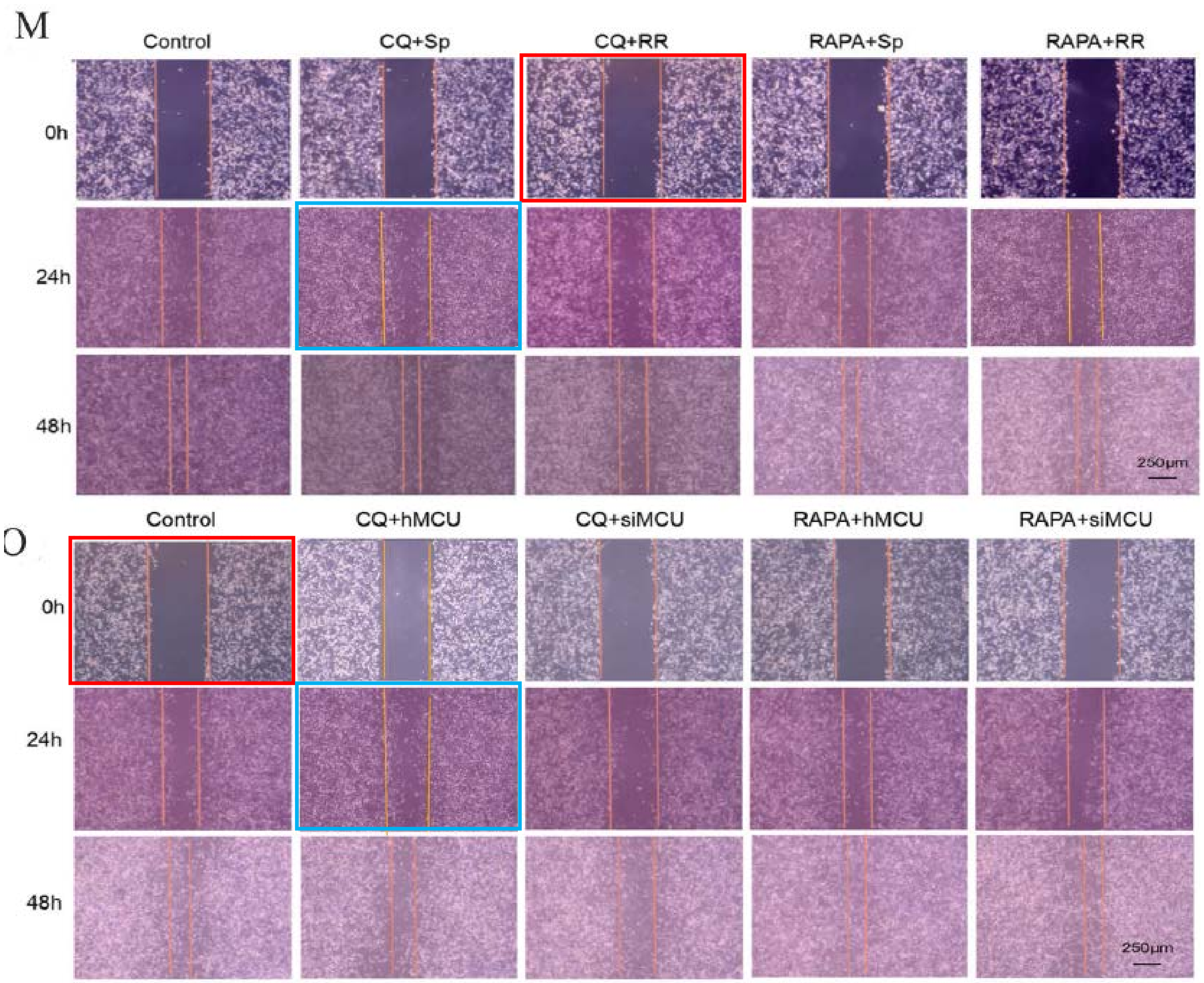


Figure 5I/M

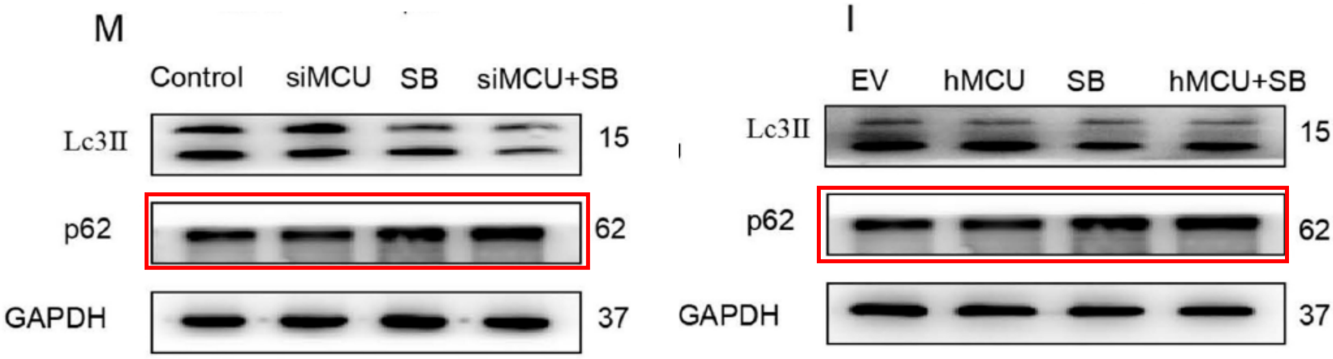

J. Cancer 2024; 15(5): 1378-1396
doi: 10.7150/jca.91798
Figure 9B

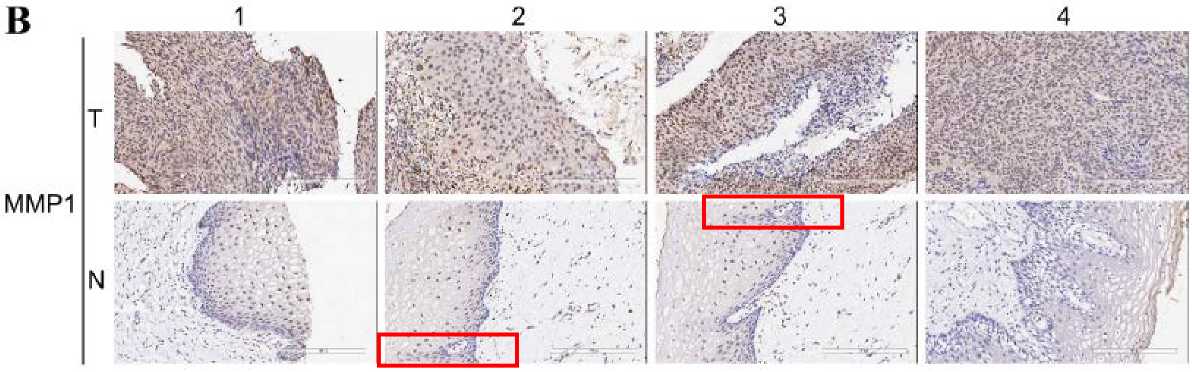

J. Cancer 2024; 15(5): 1462-1486
doi: 10.7150/jca.91944
Figure 11A

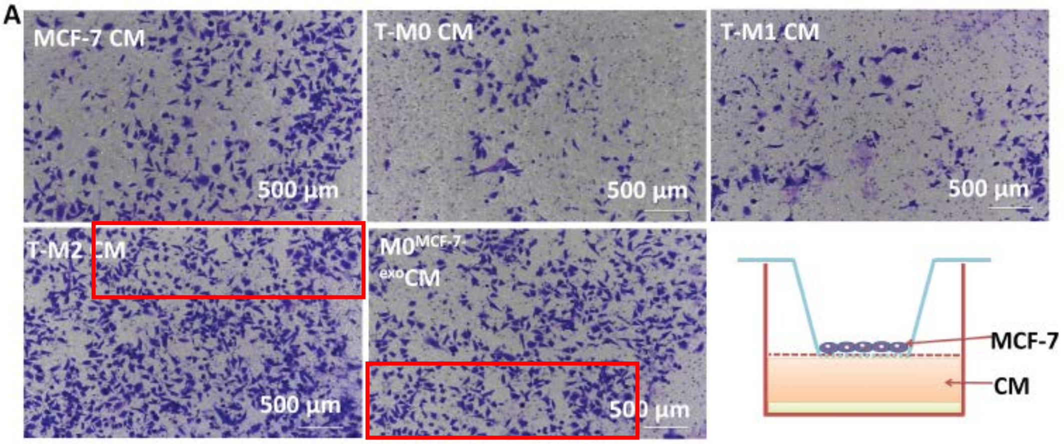

J. Cancer 2024; 15(7): 1870-1879
doi: 10.7150/jca.92723
Figures 1E/4B

Same GAPDH loading controls, different treatments

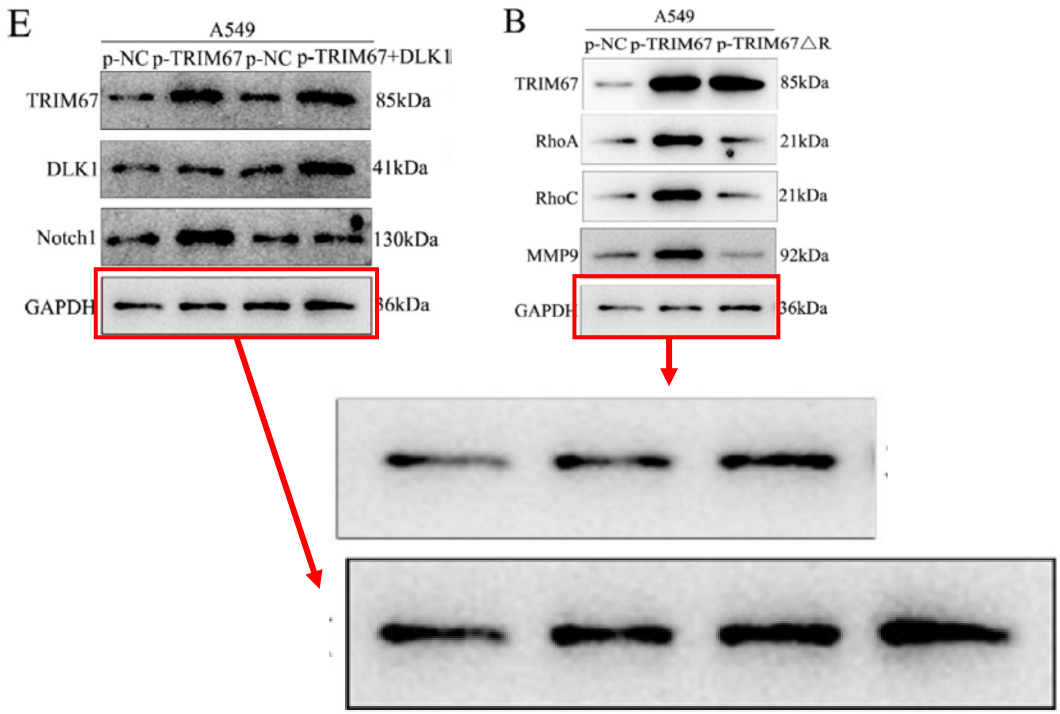

J. Cancer 2024; 15(7): 1966-1982
doi: 10.7150/jca.88887
Figure 11A

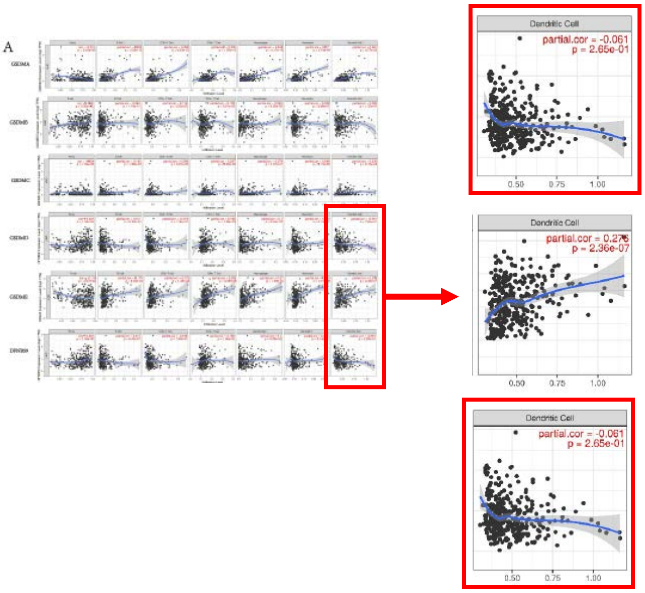

J. Cancer 2024; 15(8): 2110-2122
doi: 10.7150/jca.93373
Figure 3C/F

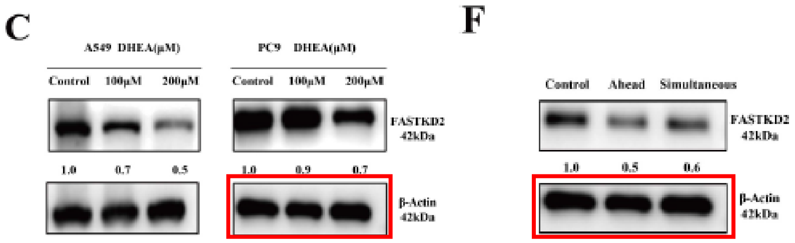

J. Cancer 2024; 15(8): 2340-2353
doi: 10.7150/jca.93890
Figure 4C

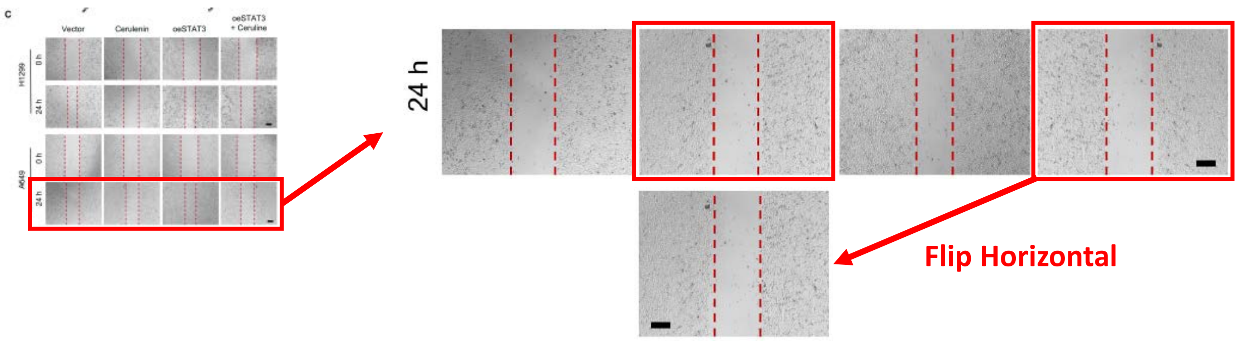

J. Cancer 2024; 15(8): 2361-2372
doi: 10.7150/jca.92087
Figure 3H

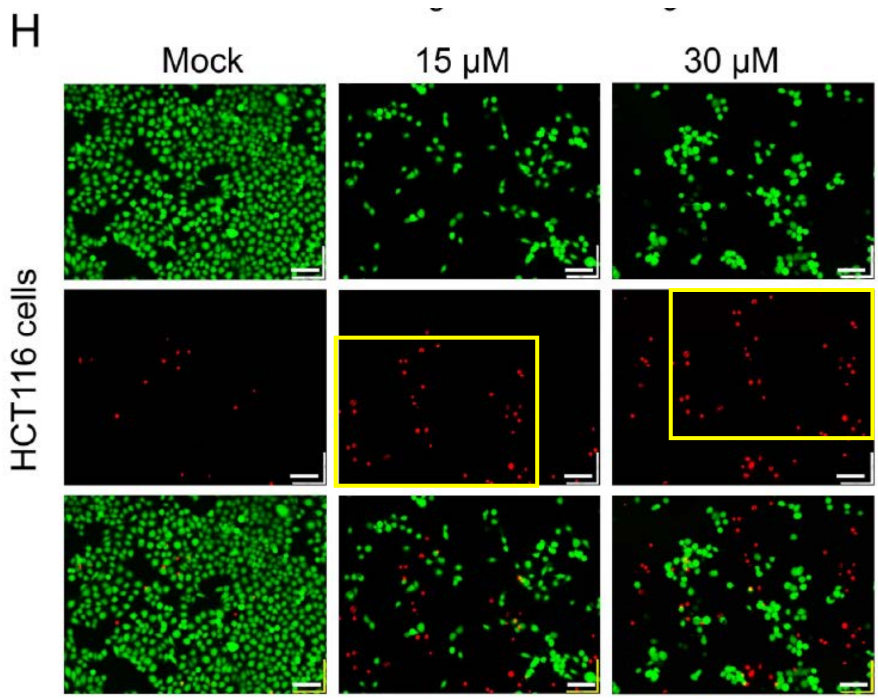


Figure 4A/B

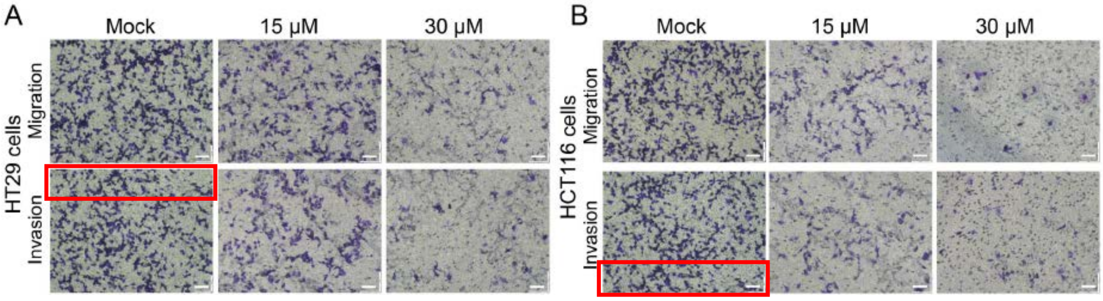

J. Cancer 2024; 15(9): 2448-2459
doi: 10.7150/jca.91566

Figures 5A (VEGF treated) & 7A (TGF treated)

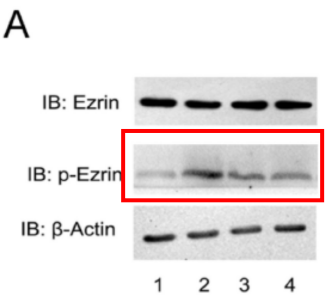


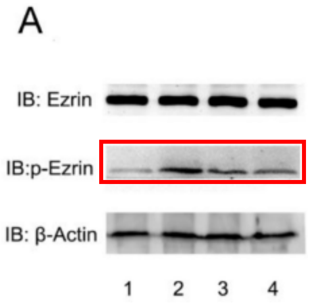


Figure 6B

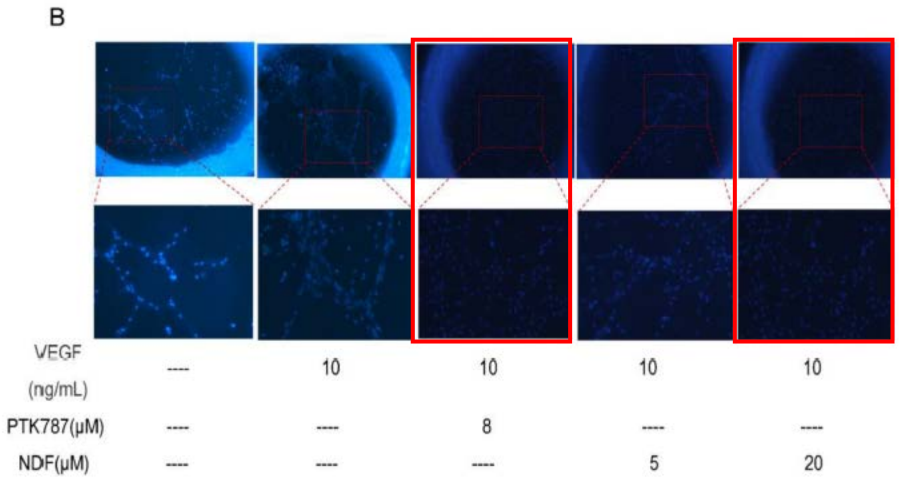


Figure 6C

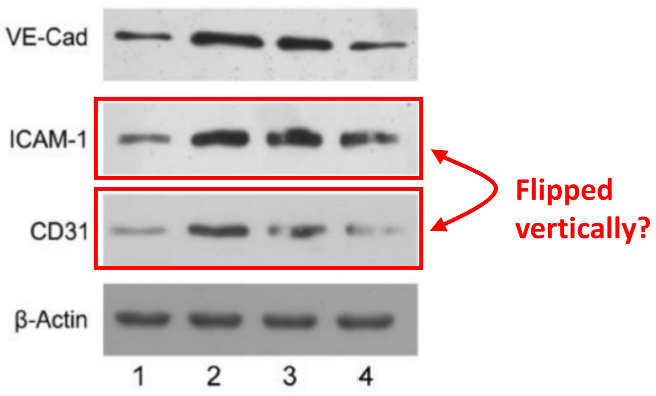

J. Cancer 2024; 15(9): 2561-2572
doi: 10.7150/jca.92386
Figures 1C & 6C

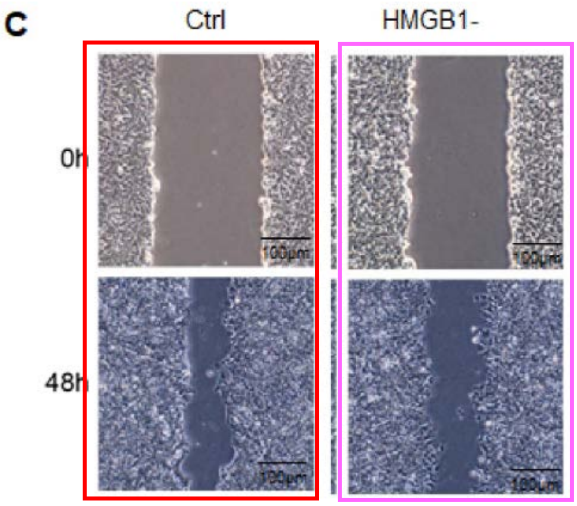


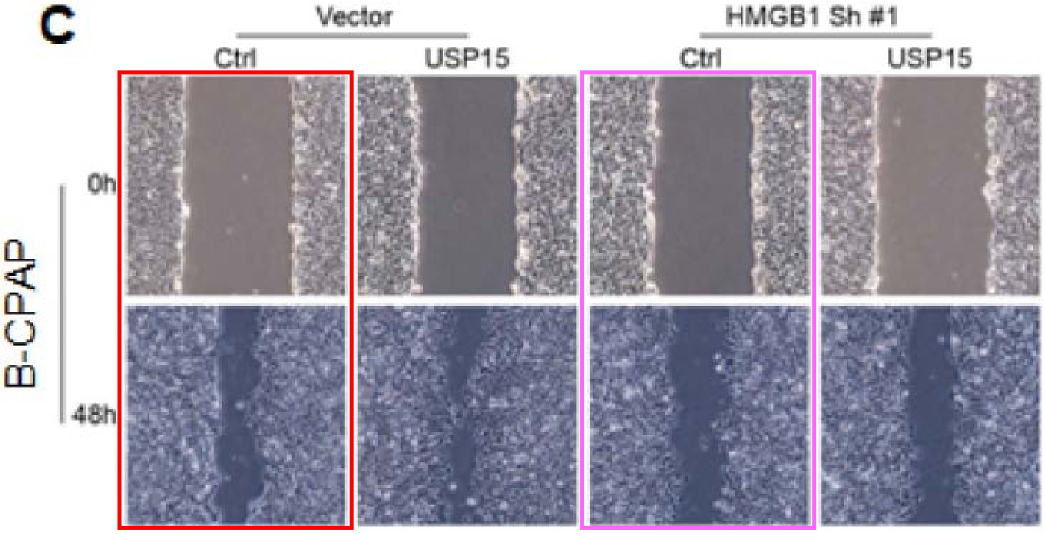

J. Cancer 2024; 15(9): 2601-2612
doi: 10.7150/jca.91955
Figures 2A-H vs. 5G/H

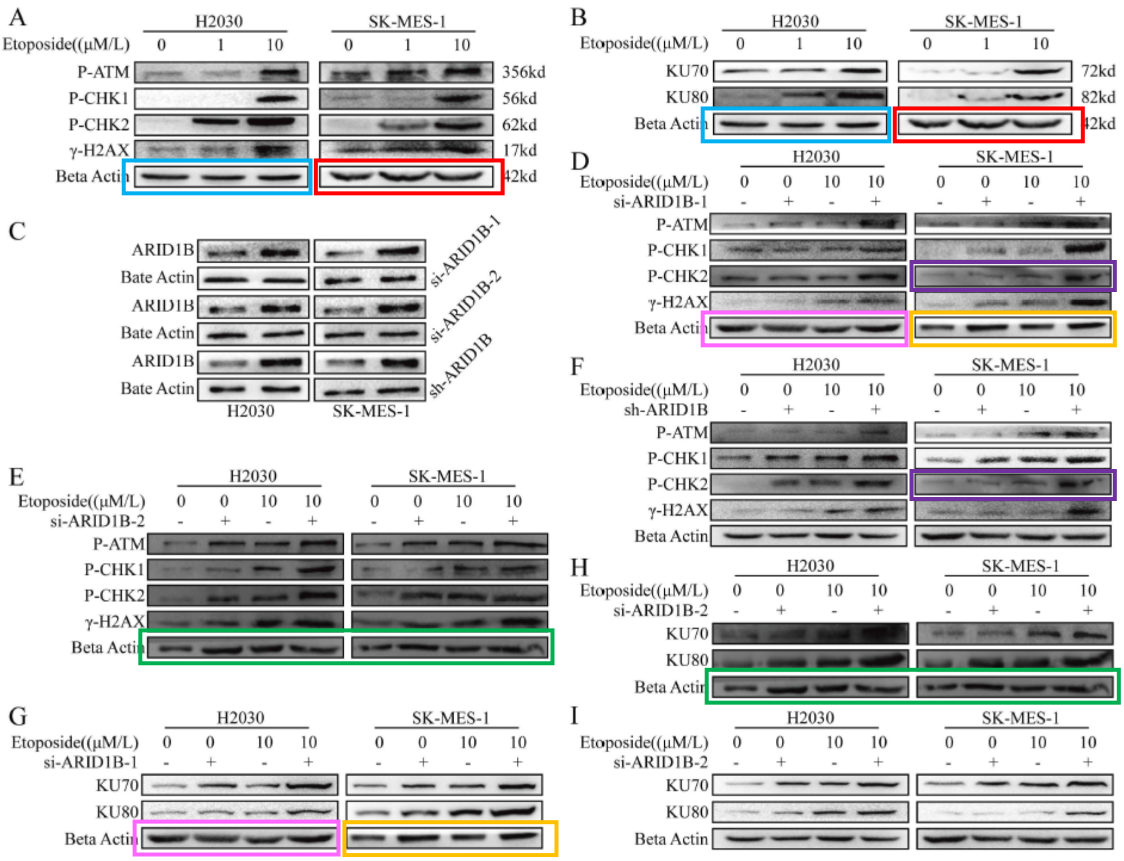


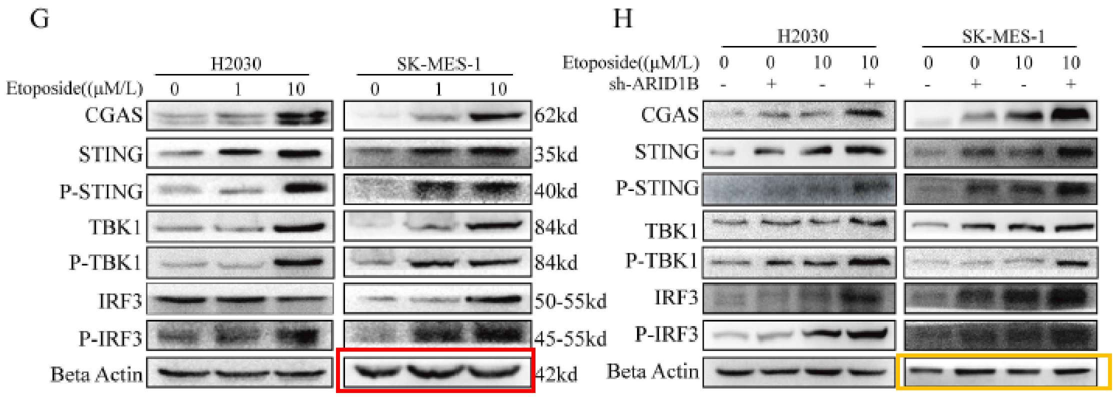

J. Cancer 2024; 15(10): 3034-3044
doi: 10.7150/jca.88612
Figure 2B

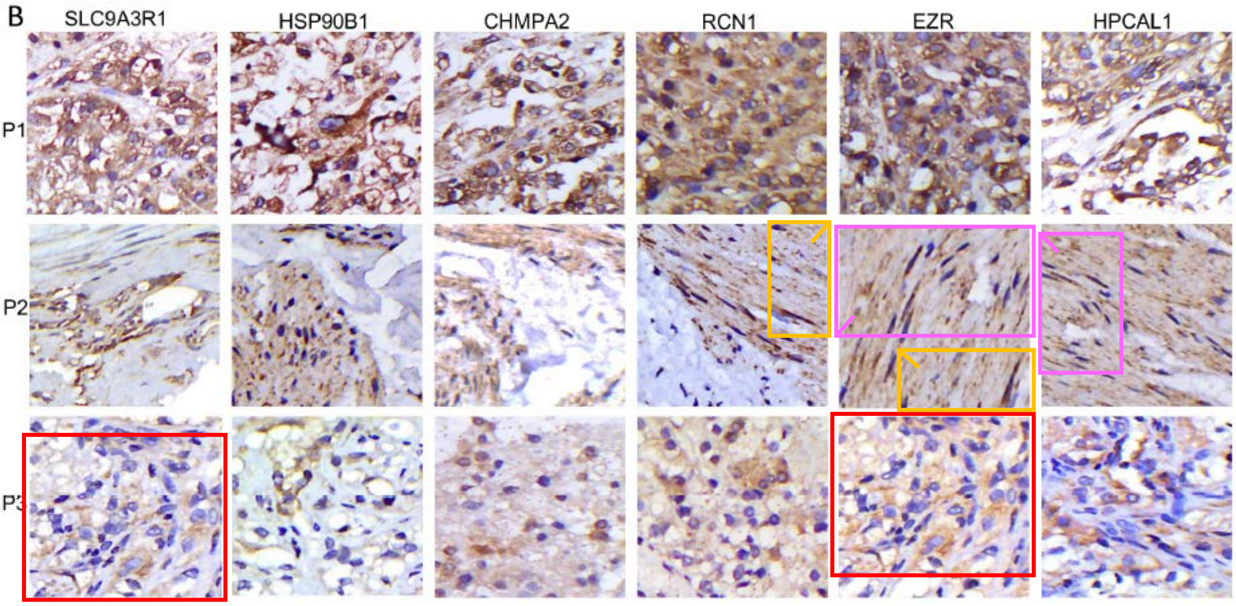

J. Cancer 2024; 15(10): 3183-3198
doi: 10.7150/jca.93789

Figure 5A

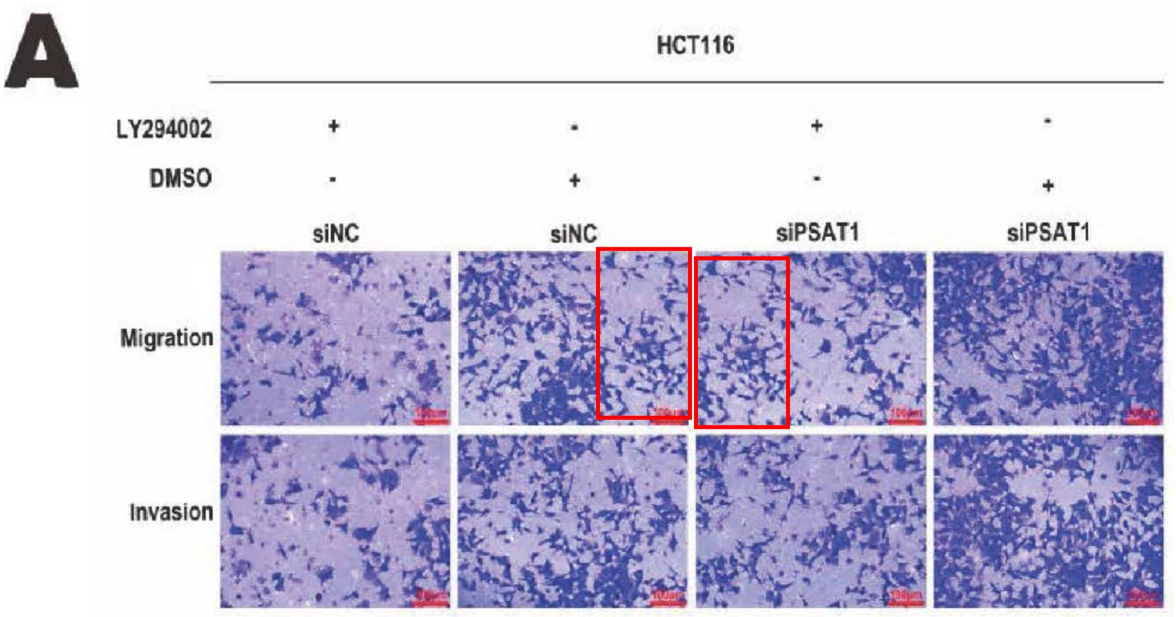


Figures 2D & 5B

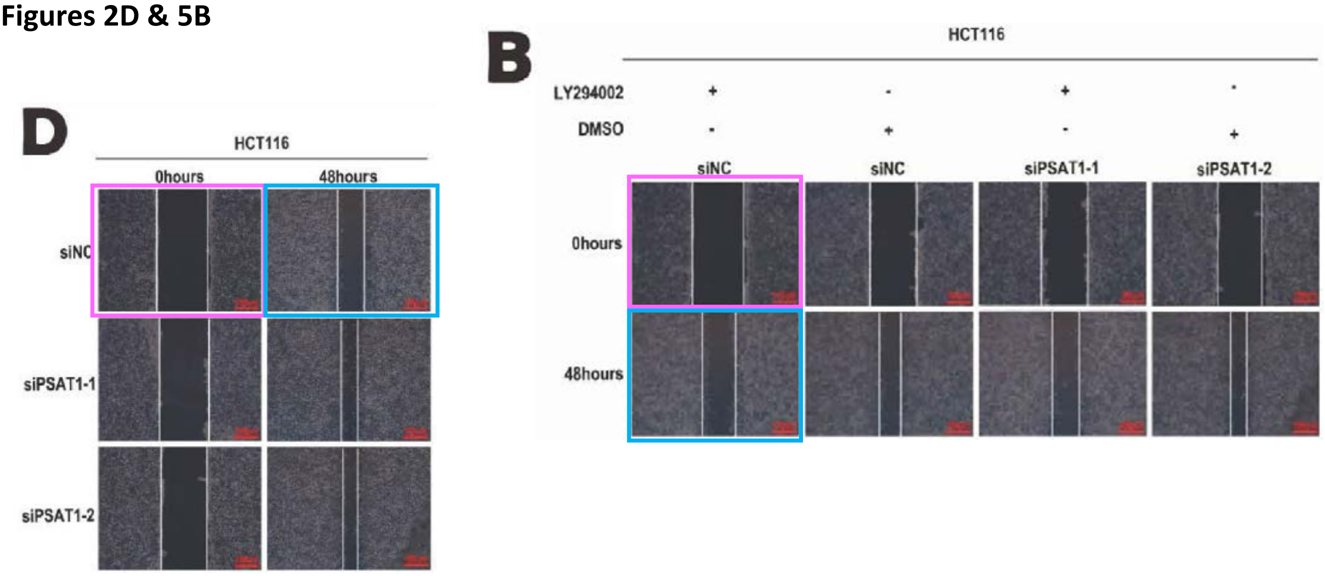

J. Cancer 2024; 15(11): 3242-3253
doi: 10.7150/jca.94343

Figure 1G/H

Same loading control (different exposure) used for different sets of patient samples

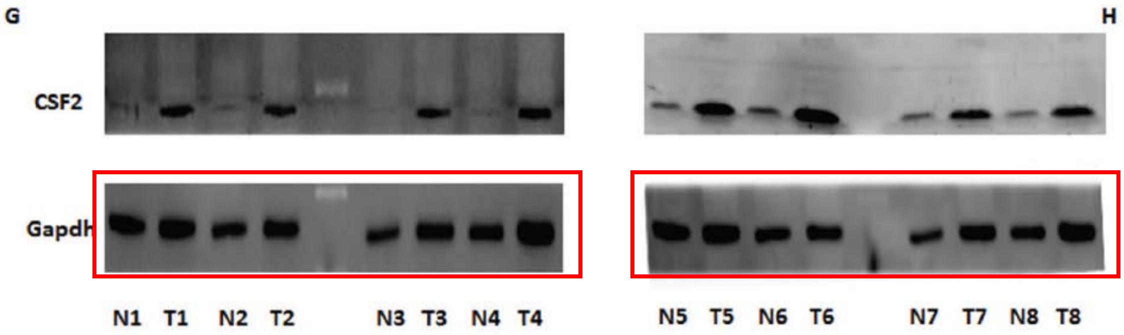


Figure 3G vs. 5F

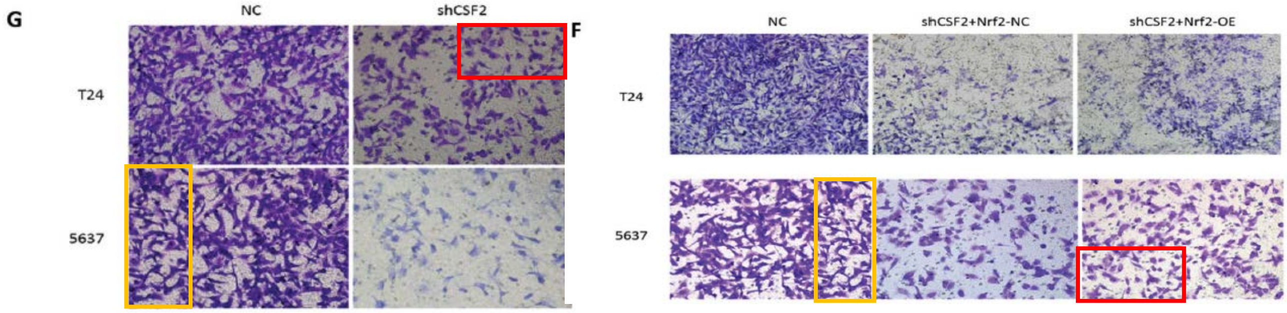


Figure 3C vs. 5E

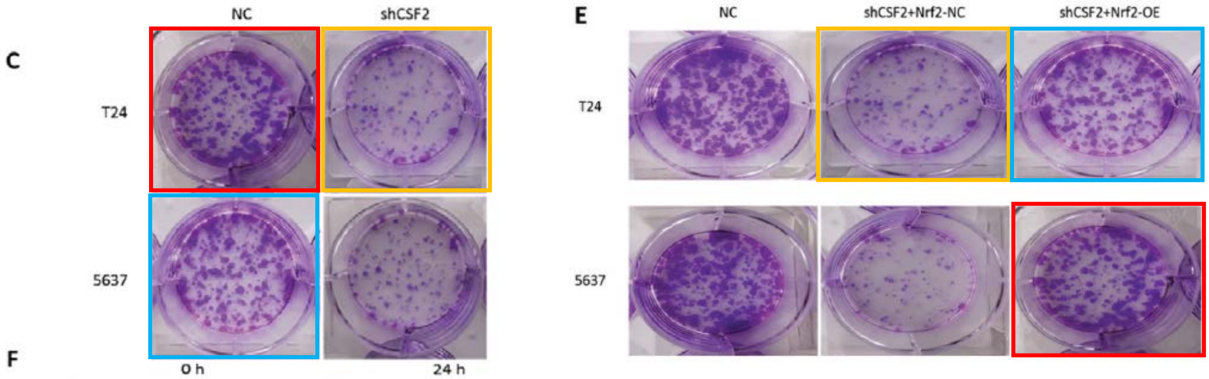

J. Cancer 2024; 15(11): 3297-3312
doi: 10.7150/jca.95549
Figure 3B vs. 5C

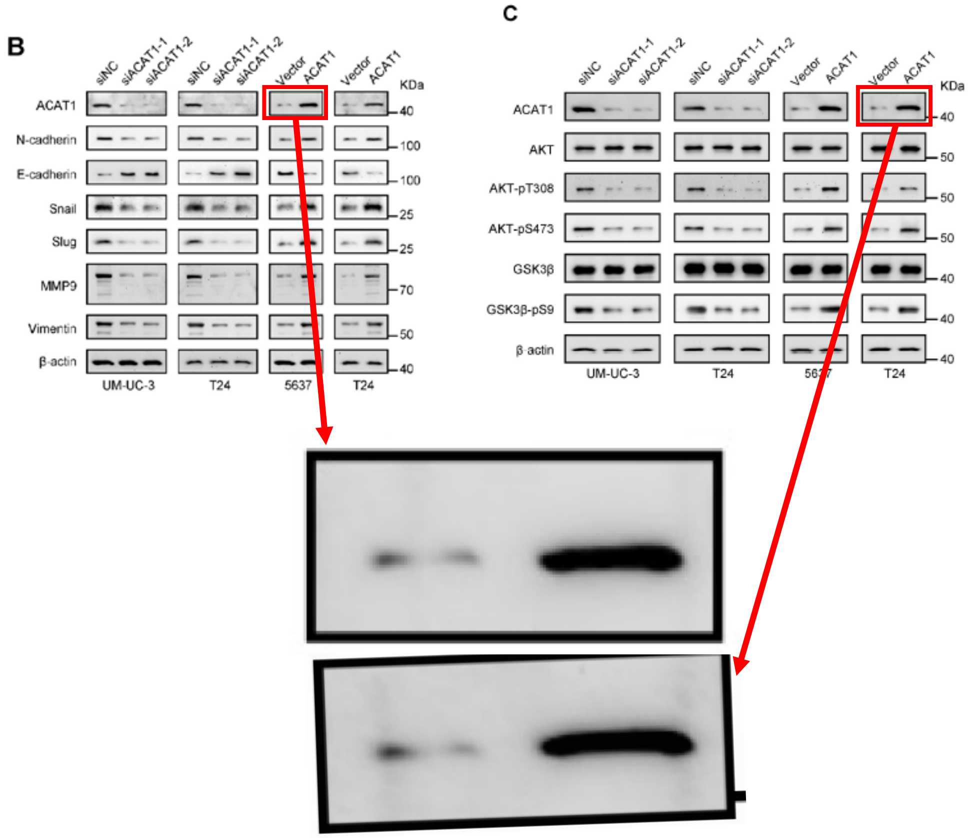

J. Cancer 2024; 15(11): 3466-3480
doi: 10.7150/jca.94452
Figure 4 vs. 6B

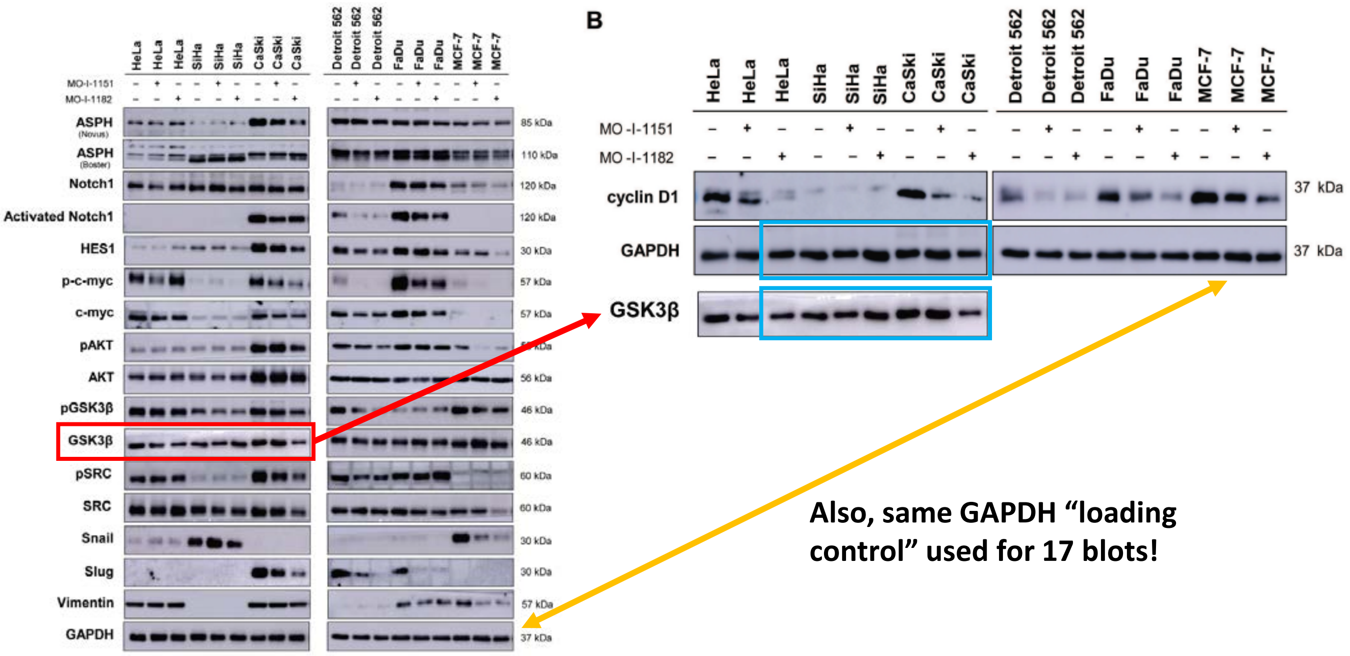

J. Cancer 2024; 15(11): 3481-3494
doi: 10.7150/jca.94685
Figure 8B

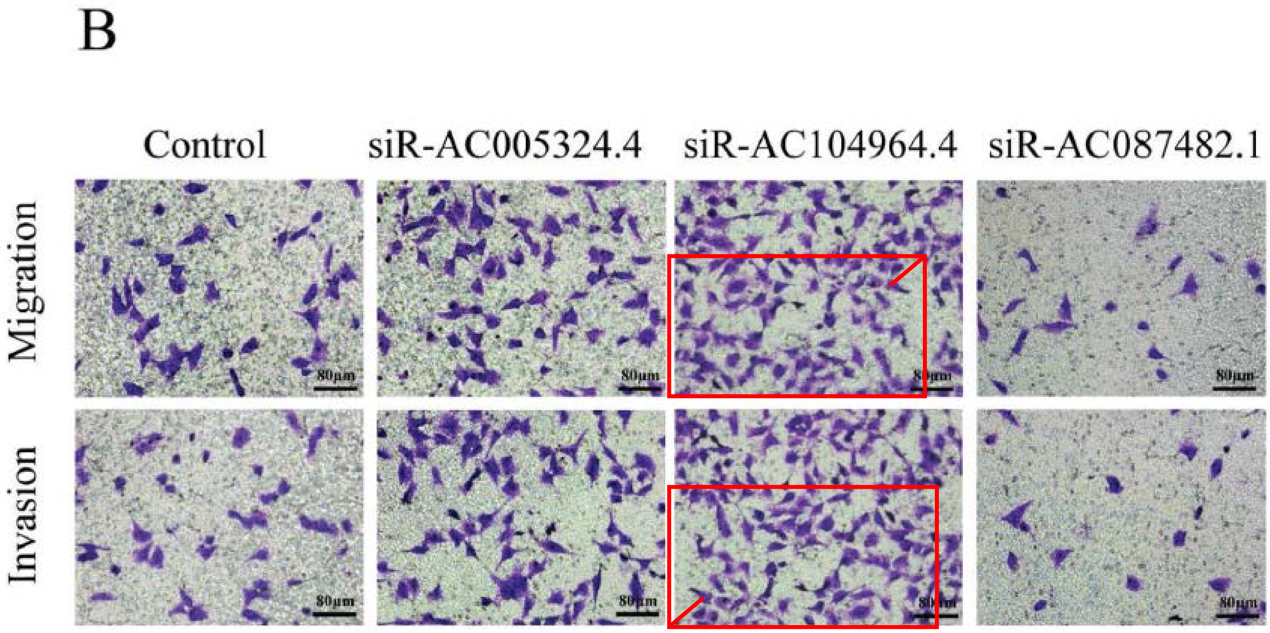

J. Cancer 2024; 15(12): 3645-3662
doi: 10.7150/jca.94902

Figure 10E

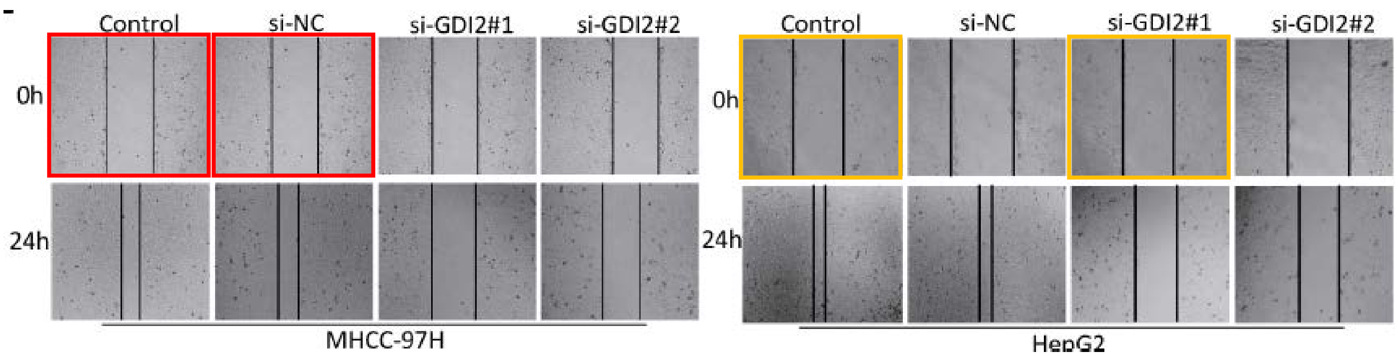


Figure 10H

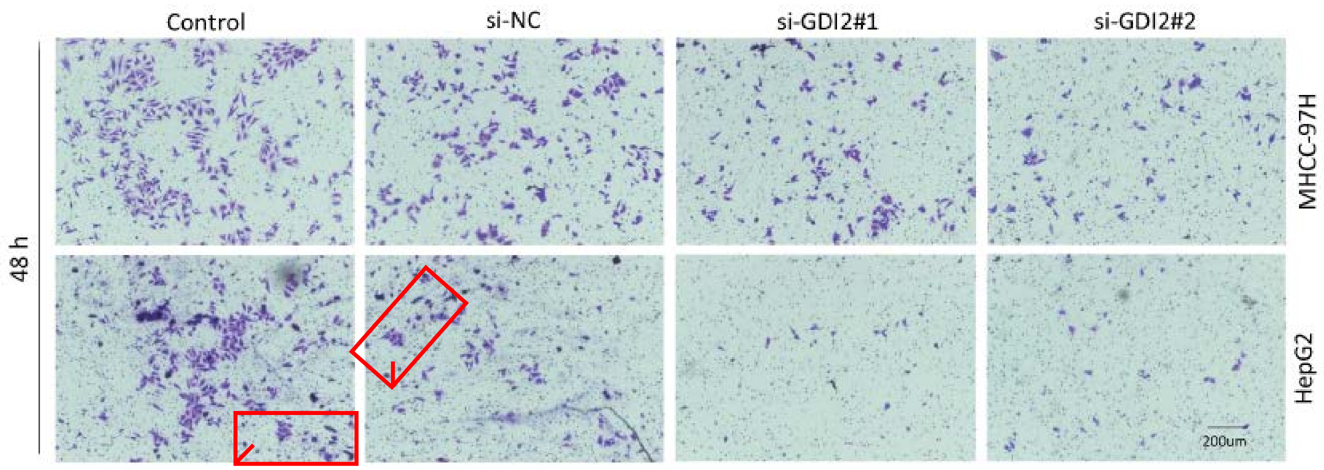

J. Cancer 2024; 15(12): 3913-3929
doi: 10.7150/jca.93208
Figure 4C

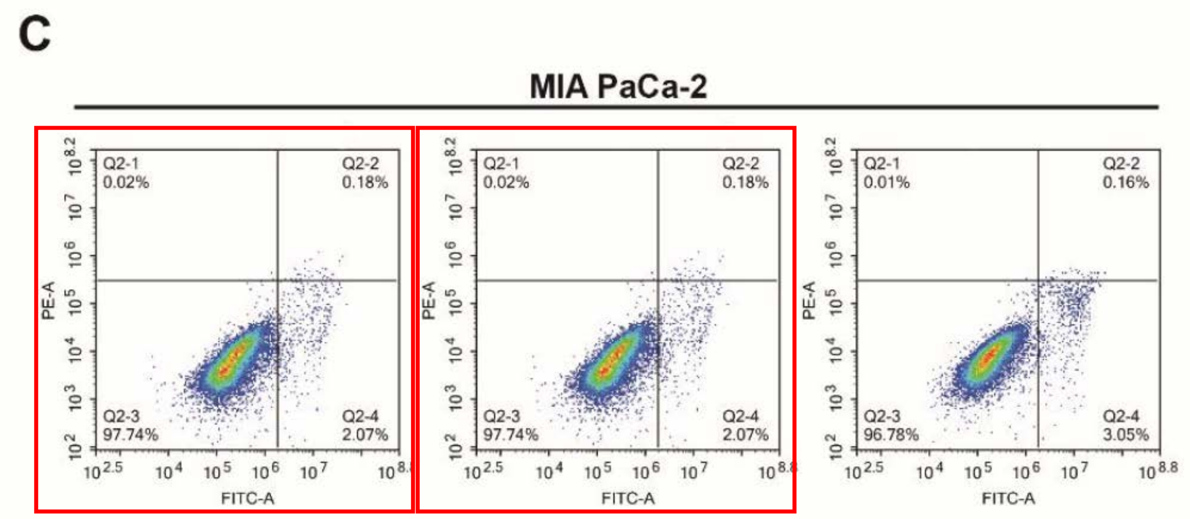

J. Cancer 2024; 15(13): 4097-4112
doi: 10.7150/jca.96306

#### Figure 1D

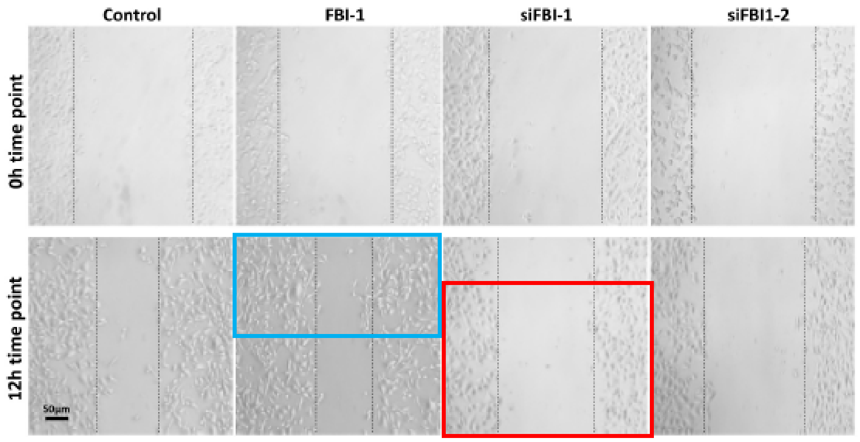


#### Figure 3E

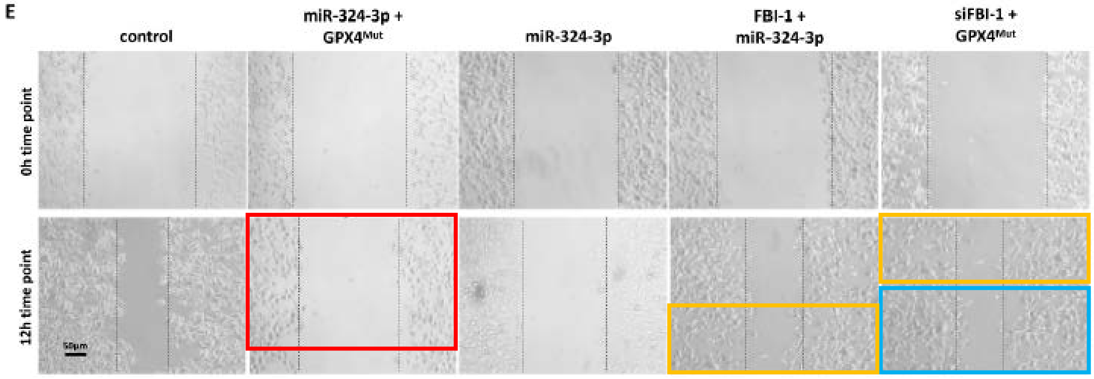

J. Cancer 2024; 15(14): 4448-4466
doi: 10.7150/jca.96500
**Figure 4B vs. 8C**

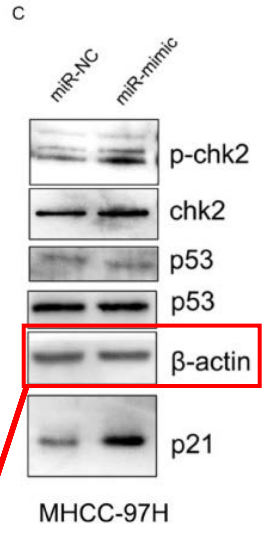


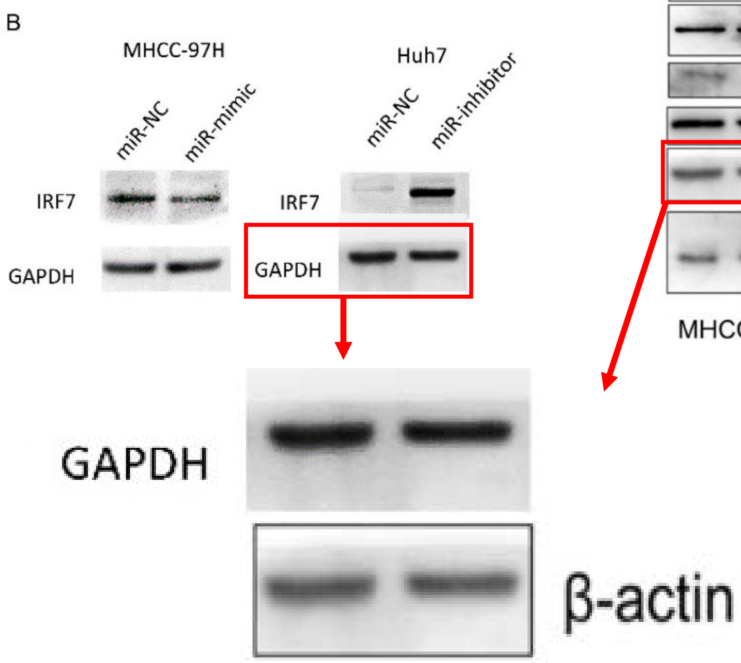


**Figure 4E vs. 6H**

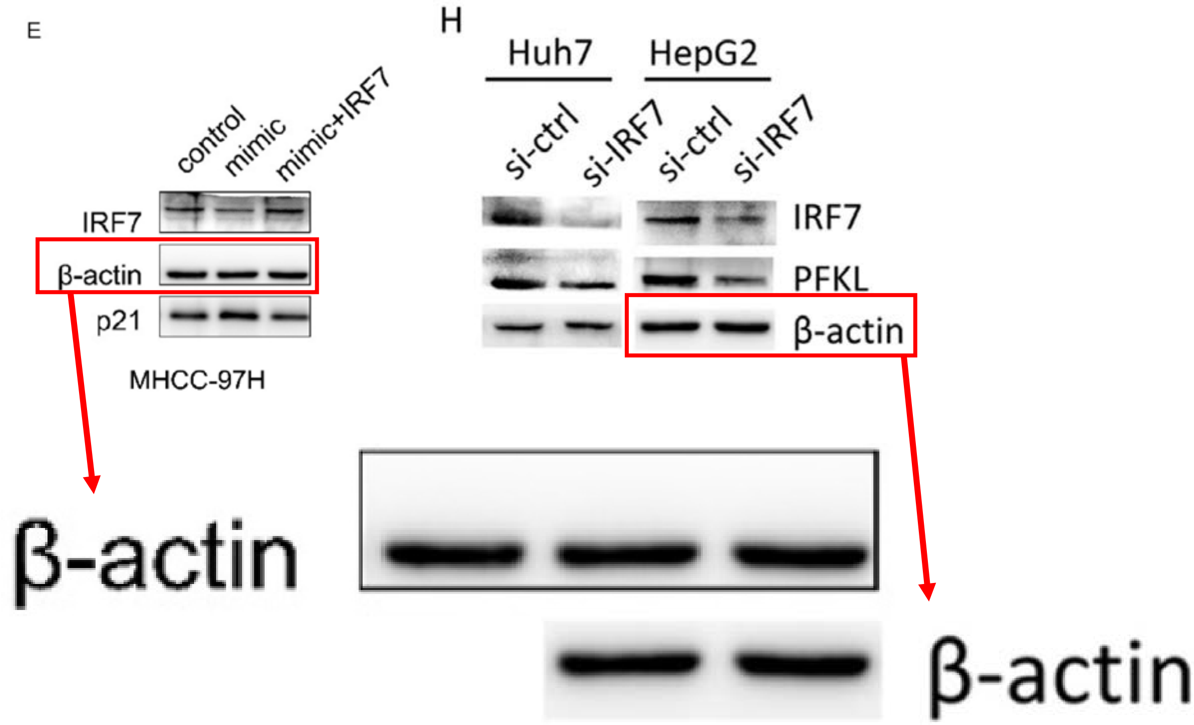

J. Cancer 2024; 15(14): 4490-4502
doi: 10.7150/jca.96185

Figure 4B

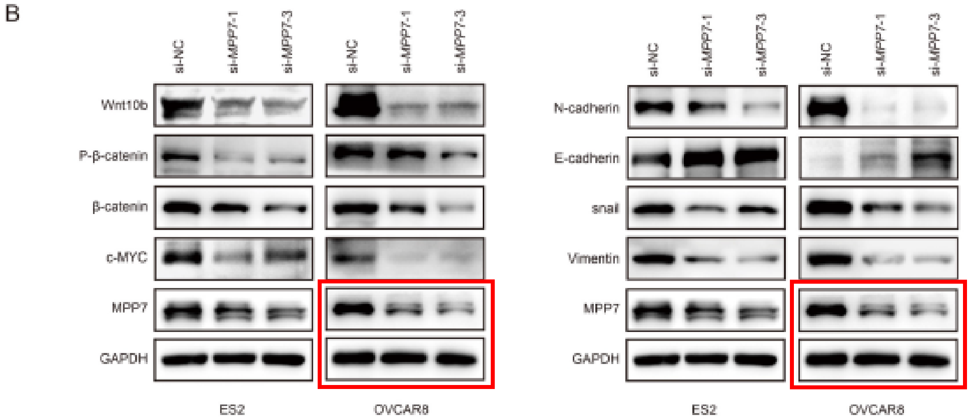


Figure 4C

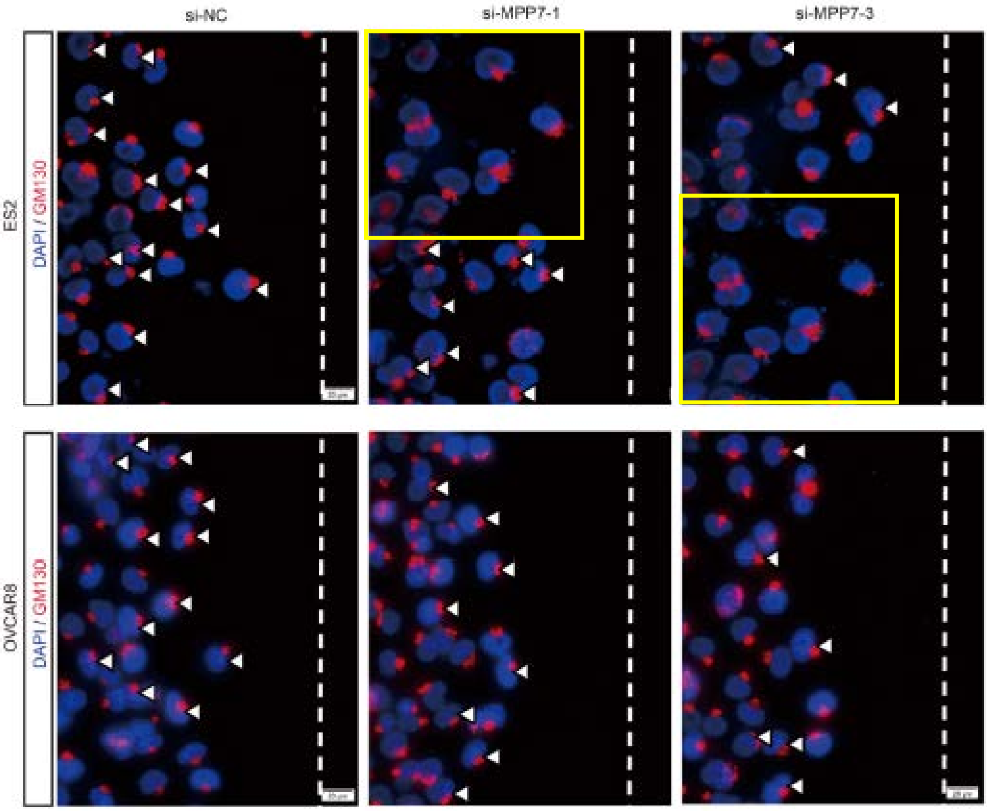

J. Cancer 2024; 15(14): 4534-4550
doi: 10.7150/jca.95407

**Figure 3A**

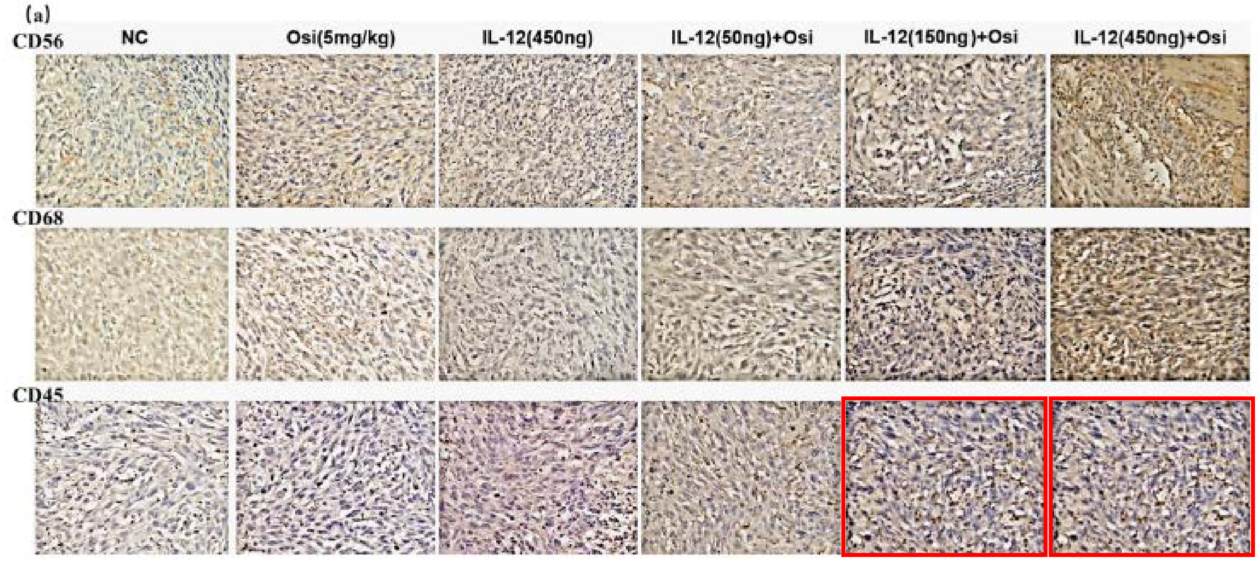


**Figure 7A/B vs. 8A/B**

**"loading controls" shared across 12 different blots**

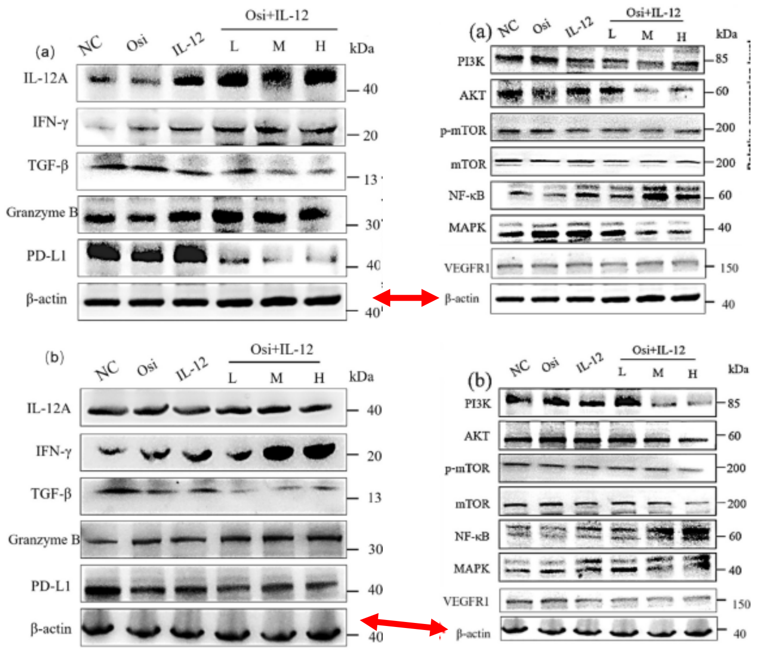

J. Cancer 2024; 15(14): 4801-4817
doi: 10.7150/jca.98075
Figure 2D

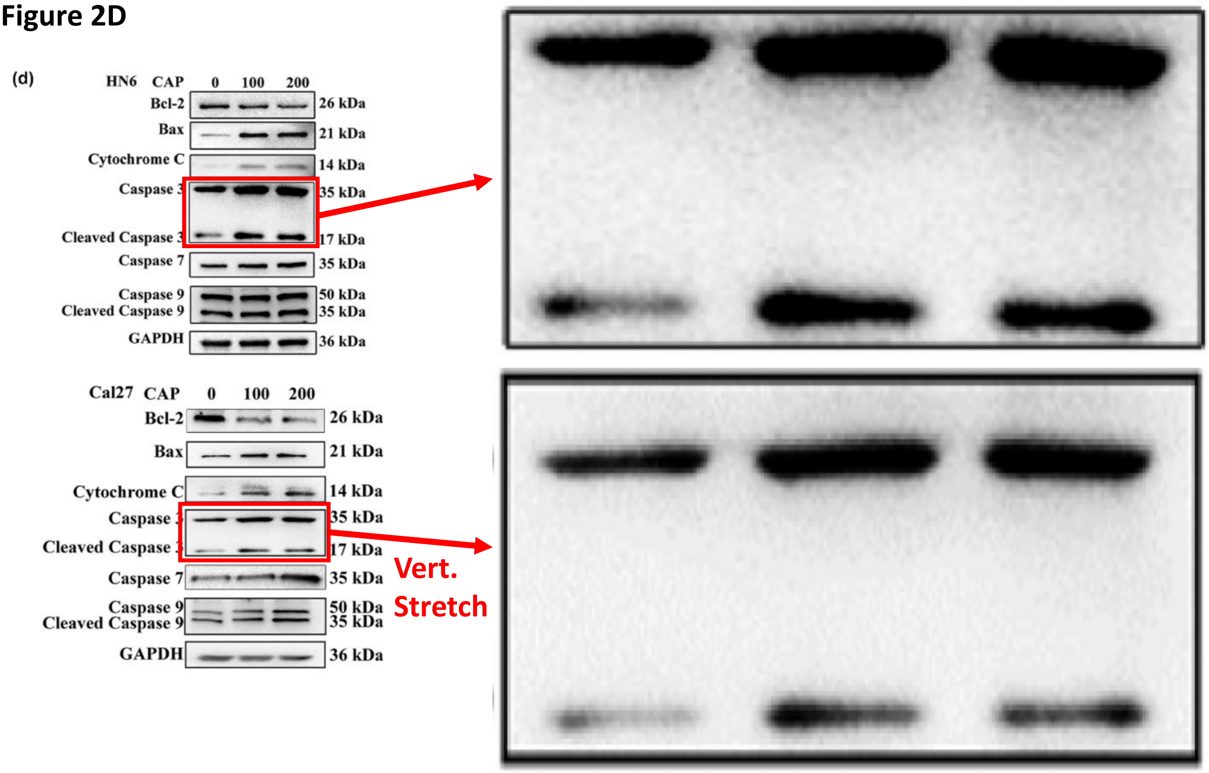

J. Cancer 2024; 15(15): 5007-5019
doi: 10.7150/jca.95816
Figure 3G vs. 6C

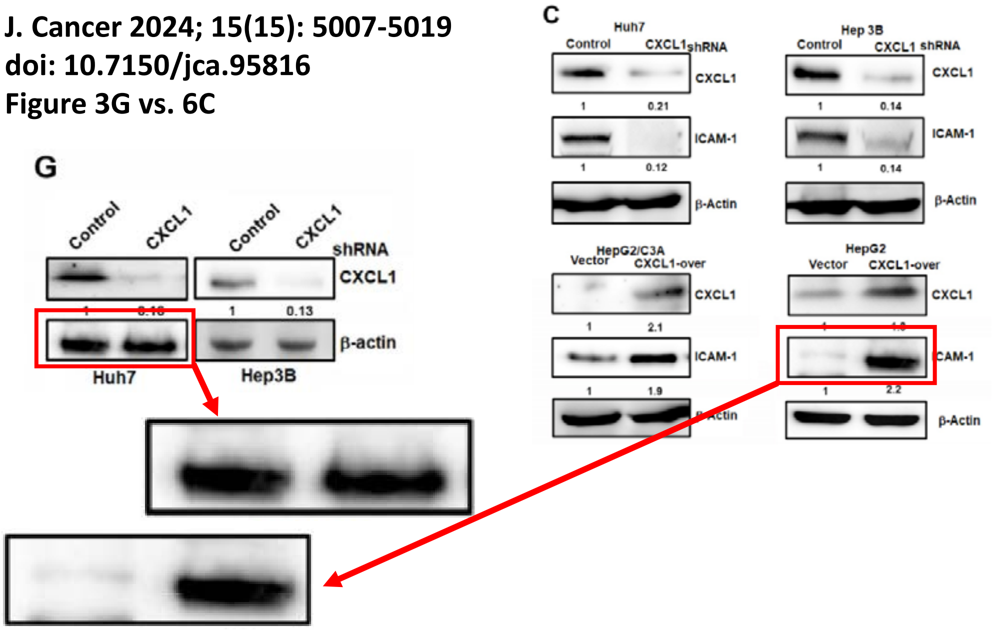

J. Cancer 2024; 15(16): 5149-5164
doi: 10.7150/jca.98238
Figure 4A vs. 6F

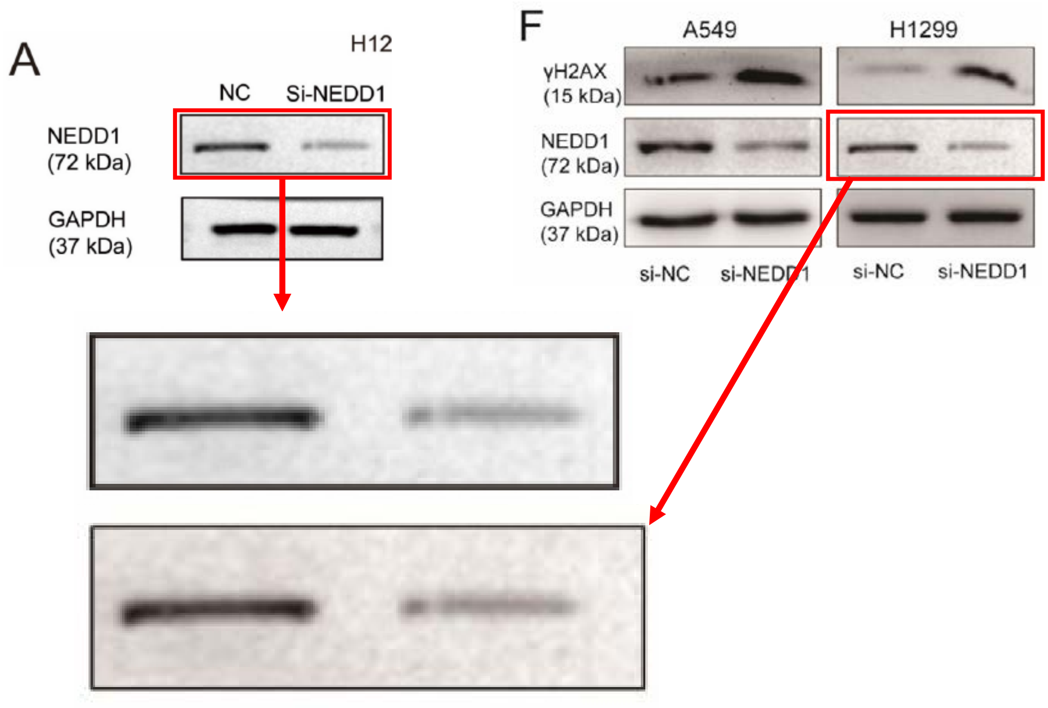

J. Cancer 2024; 15(16): 5288-5307
doi: 10.7150/jca.98738
Figure 1C

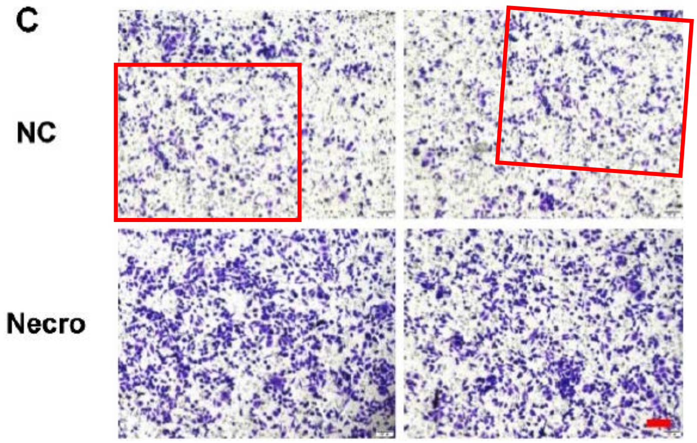


Figure 1D

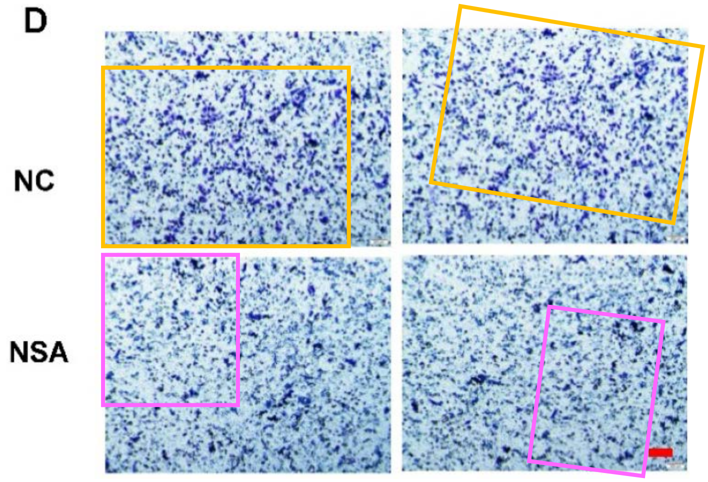

J. Cancer 2024; 15(16): 5308-5317
doi: 10.7150/jca.96001
Figure 2A/B

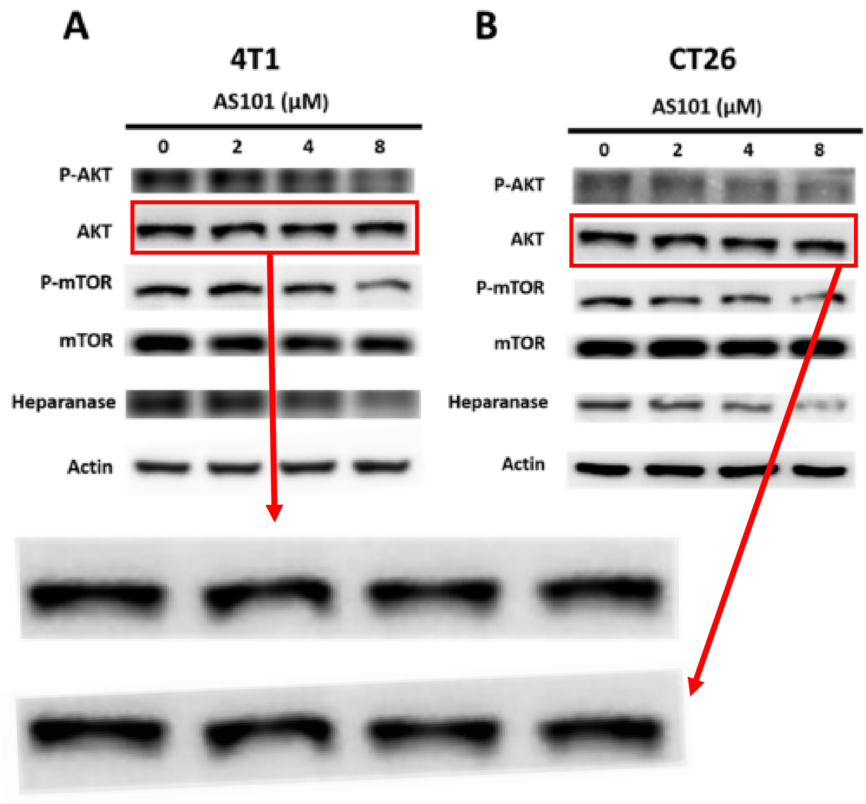

J. Cancer 2024; 15(16): 5351-5366
doi: 10.7150/jca.88767
Figure 7E

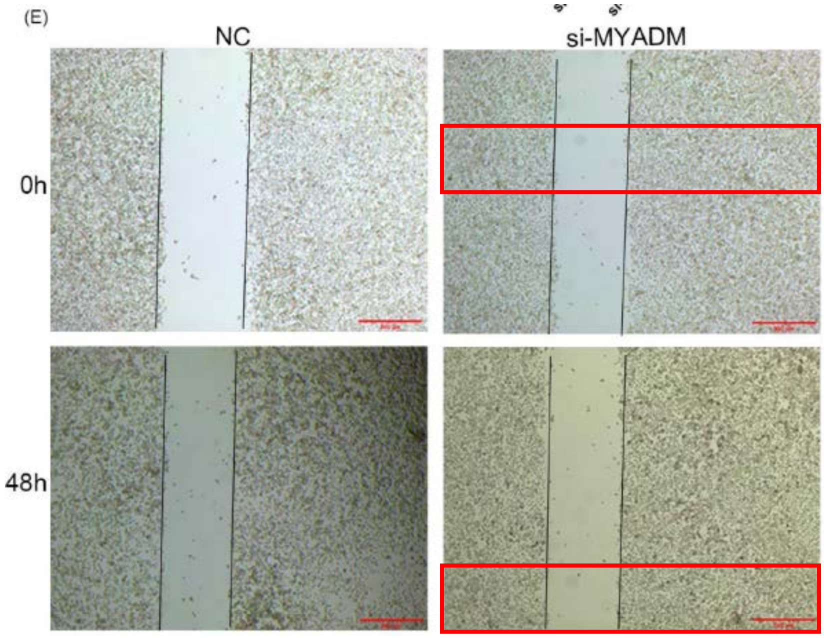

J. Cancer 2024; 15(17): 5691-5709
doi: 10.7150/jca.98240
Figure 12

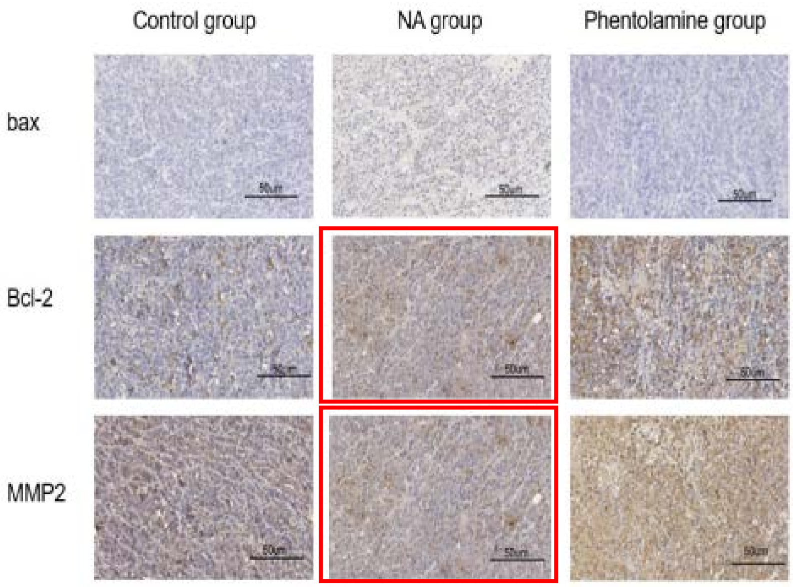

J. Cancer 2024; 15(17): 5810-5827
doi: 10.7150/jca.98915
Figure 3B

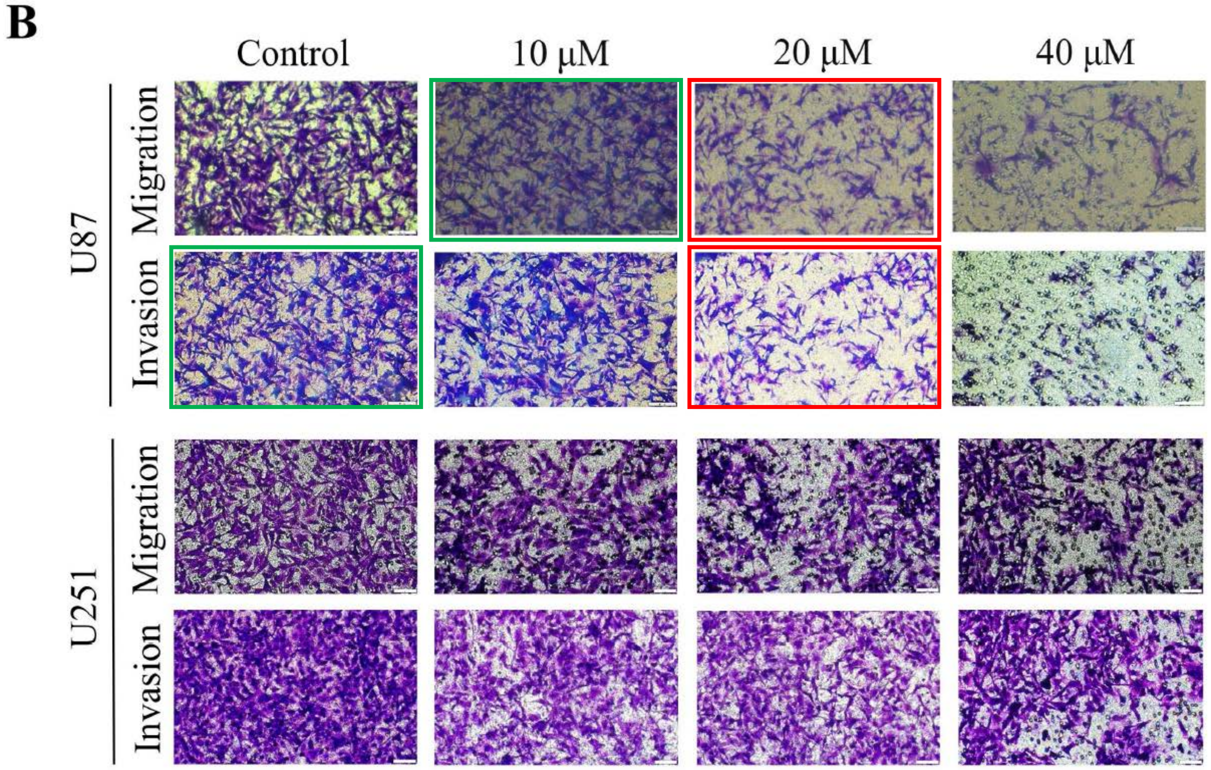

J. Cancer 2024; 15(18): 5876-5896
doi: 10.7150/jca.100706
Figure 4E vs. 10D

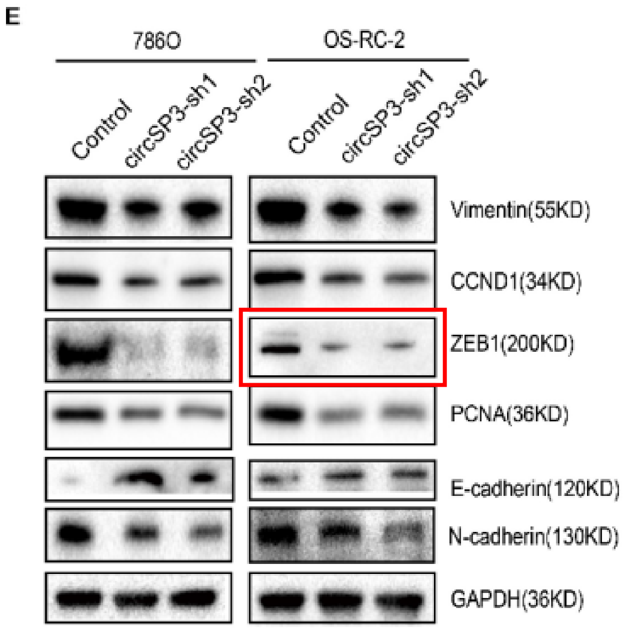


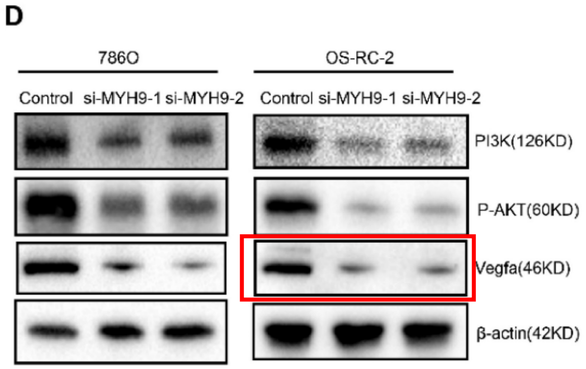


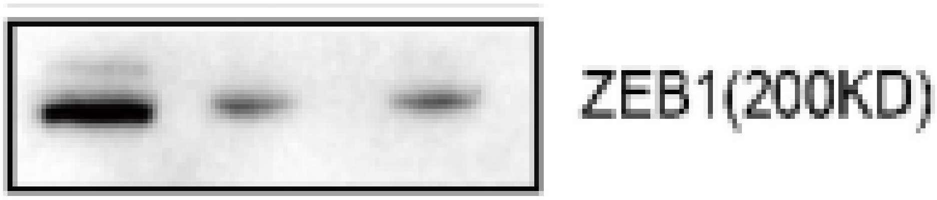


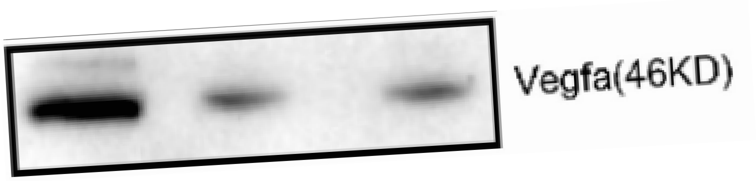

J. Cancer 2024; 15(18): 6710-6723
doi: 10.7150/jca.96162
Figure 7D

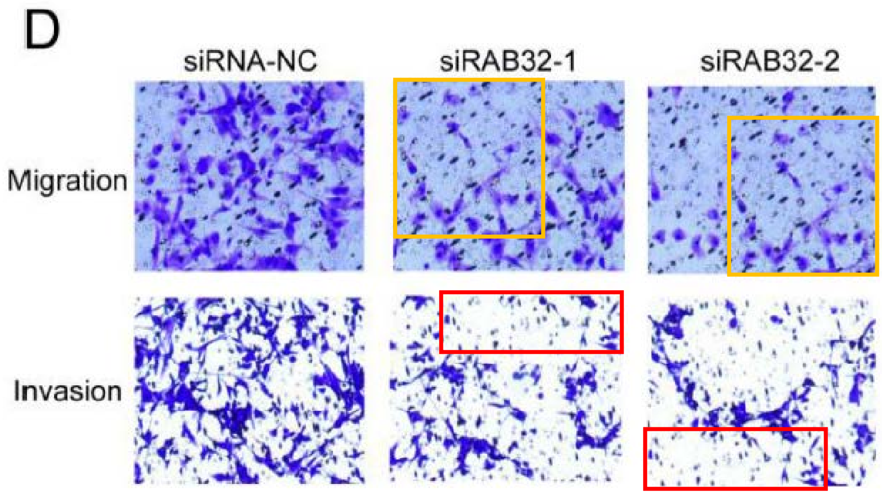
