## Supplemental file 2 (2025 content) for "Does charging for corrections in the bioscience literature disincentivize pre-publication handling of problematic image data? An ImageTwin-AI study"

### Images dupl’ from other journals

J. Cancer. 2025; 16(1): 12-33
DOI: 10.7150/jca.96848
Figure 3

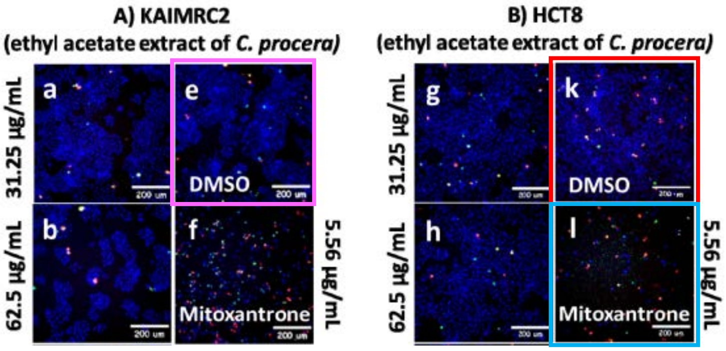


Saudi Pharm Jour. 2023; 21: 101794
DOI: 10.1016/j.jsps.2023.101794
Figure 4

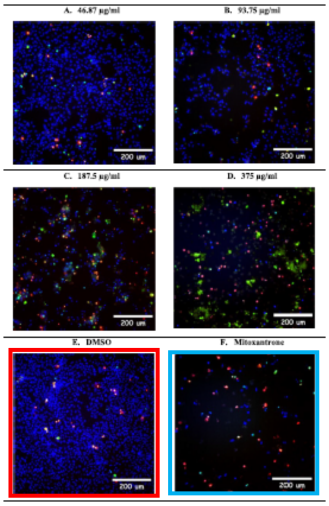


Sci. Rep. 2024; 14: 7202
DOI: 10.1038/s41598-024-57680-0
Figure 5

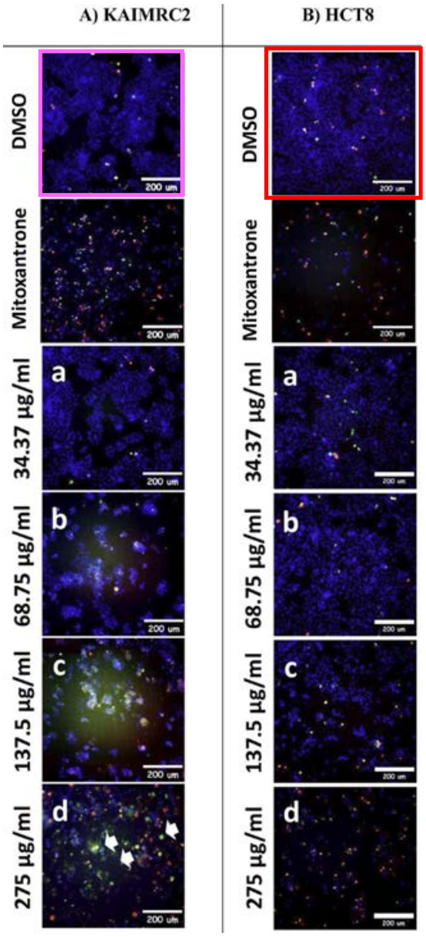

J. Cancer. 2025; 16(1): 44-54
DOI: 10.7150/jca.95266
Figure 1G/I

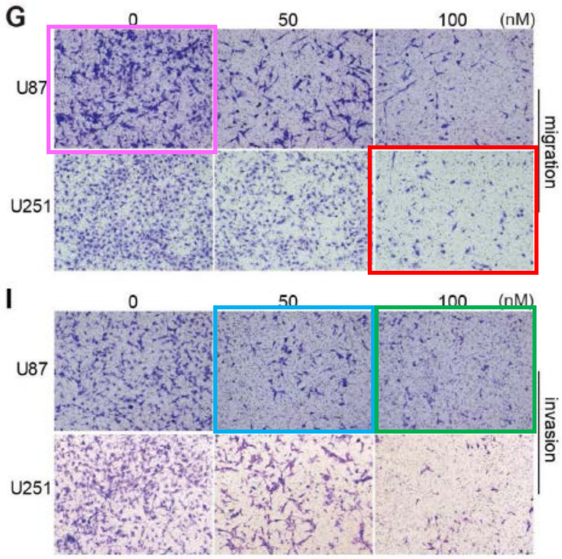


Heliyon (2024) e27722
DOI: 10.1016/j.heliyon.2024.227722
Figure 2C/E

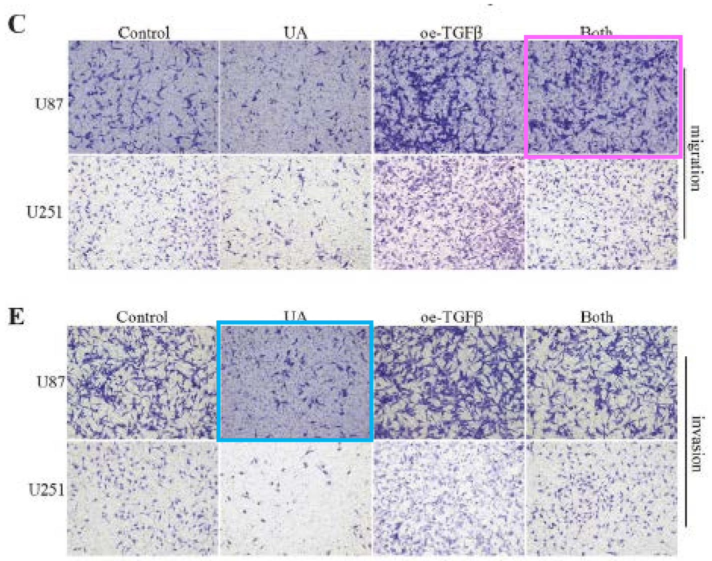


Heliyon (2024) e27722
DOI: 10.1016/j.heliyon.2024.227722
Figure 1F

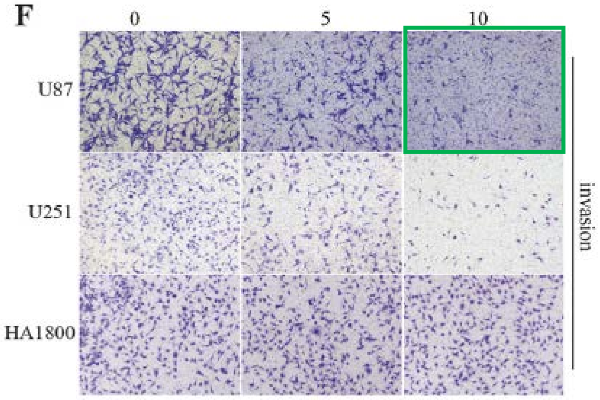


Sci Rep (2024) 14: 15758
DOI: 10.1038/s41598-024-66484-1
Figure 13F

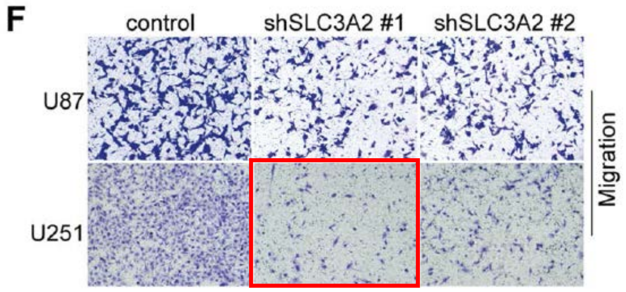

**J. Cancer. 2025; 16(1): 44-54**
**DOI: 10.7150/jca.95266**
**Figure 3C/E**

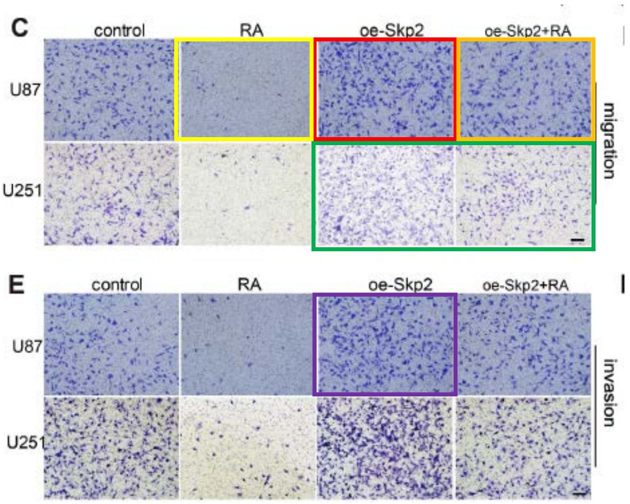


**Heliyon (2024) e27722**
**DOI: 10.1016/j.heliyon.2024.227722**
**Figure 2C/E**

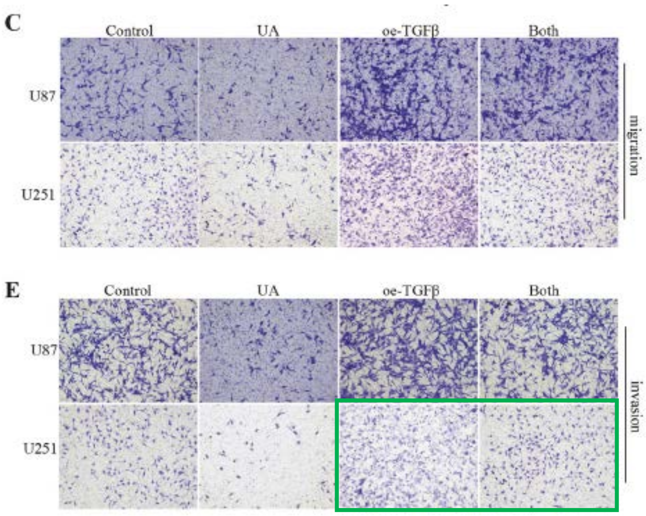


**Int J Med Sci. 2021; 18(7): 1609-1617**
**DOI: 10.7150/ijms.52206**
**Figure 2C/D**

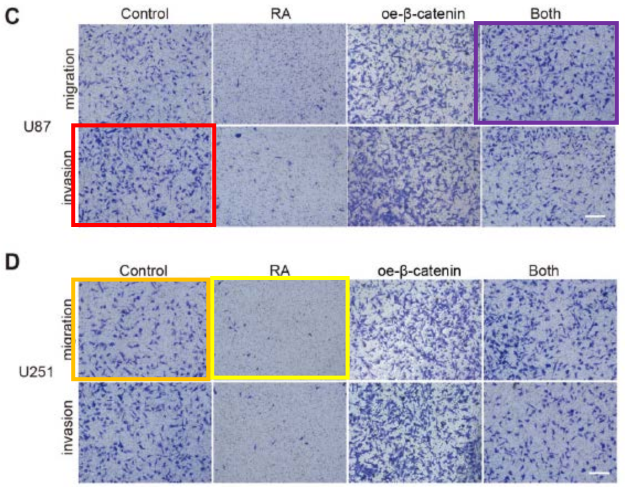

**J. Cancer. 2025; 16(2): 590-602**
**DOI: 10.7150/jca.97034**
**Figure 4B**

B

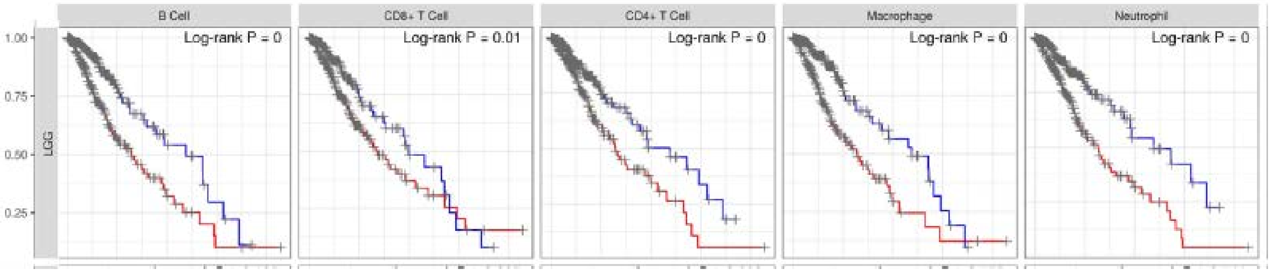


**Inflamm Immun Dis 2025; 13: e70224**
**DOI: 10.1002/iid3.70224**
**Figure 7B**

B

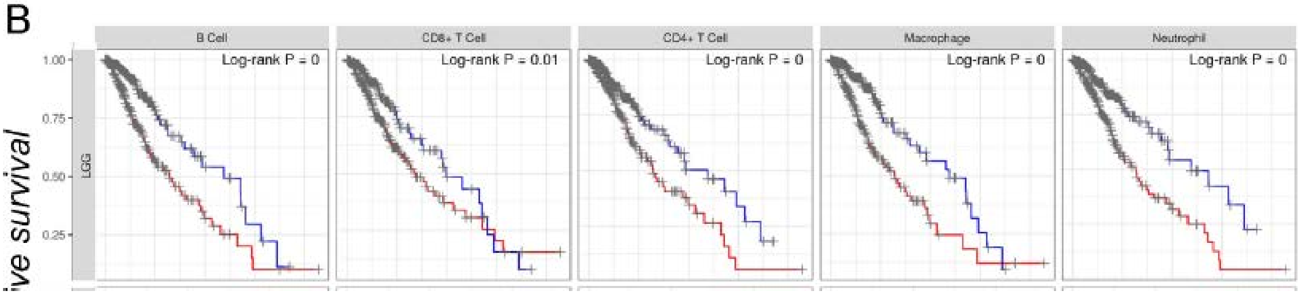


**Sci Rep (2023) 13: 3820**
**DOI: 10.1038/s41598-023-30413-5**
**Figure 9C**

C LGG

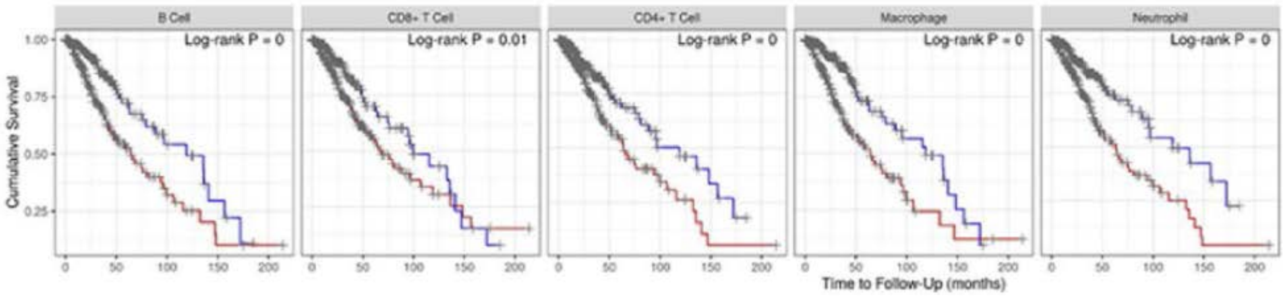

J. Cancer. 2025; 16(7): 2388-2400
DOI: 10.7150/jca.103286
Figure 7B

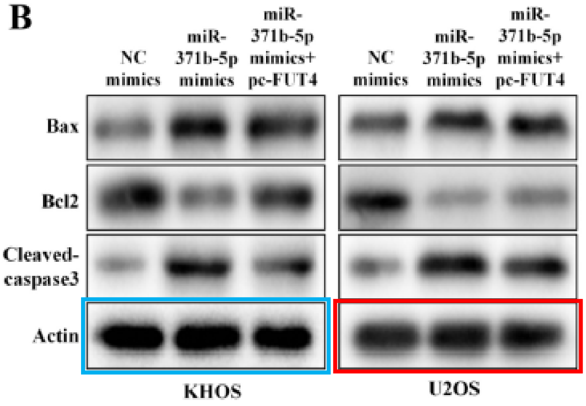


Biosci. Rep. 2021; 41: BSR20210138
DOI: 10.1042/BSR20210138
Figure 2F/H

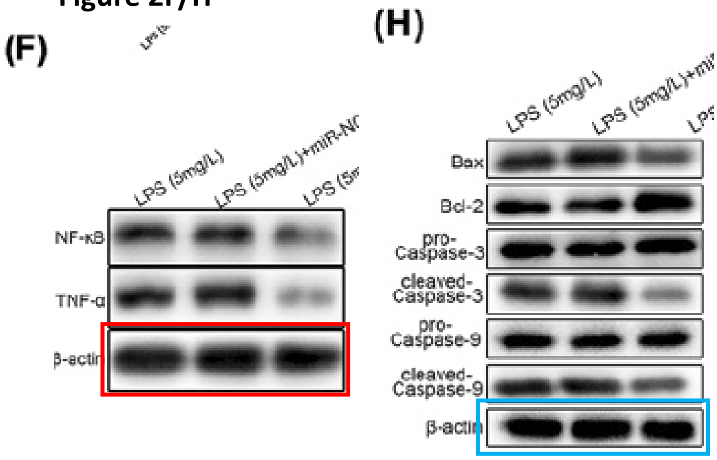


J. Cancer. 2025; 16(7): 2388-2400
DOI: 10.7150/jca.103286
Figure 7A

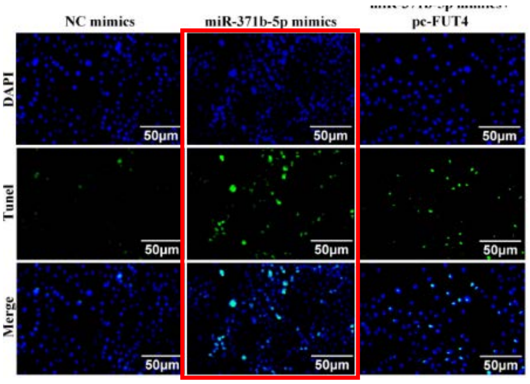


Comput. Math Methods Med. 2021; ID 3957738
DOI: 10.1155/2021/3957738 *RETRACTED*
Figure 5D

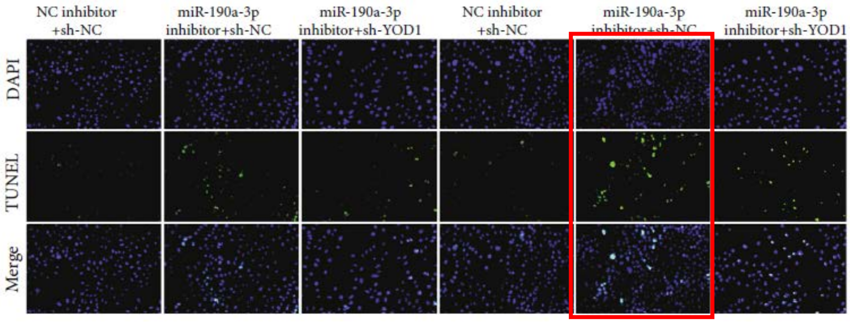

J. Cancer. 2025; 16(13): 3884-3896
DOI: 10.7150/jca.115525
Figure 7B

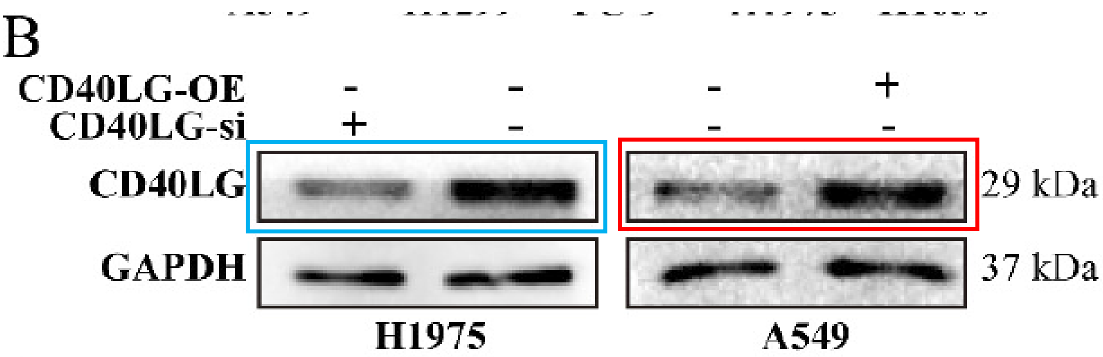


J. Cancer. 2024; 15(9): 2601-2612
DOI: 10.7150/jca.91955
Figure 2C

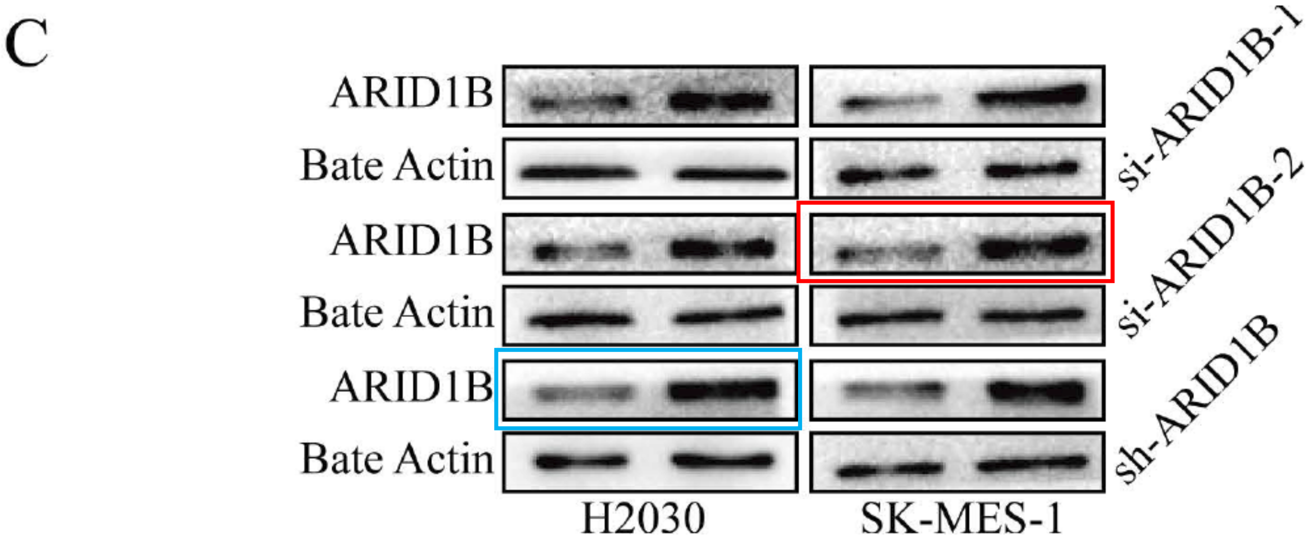

### Image problems within *J. Cancer* papers

J. Cancer. 2025; 16(1): 315-330
DOI: 10.7150/jca.102618

Figure 3B

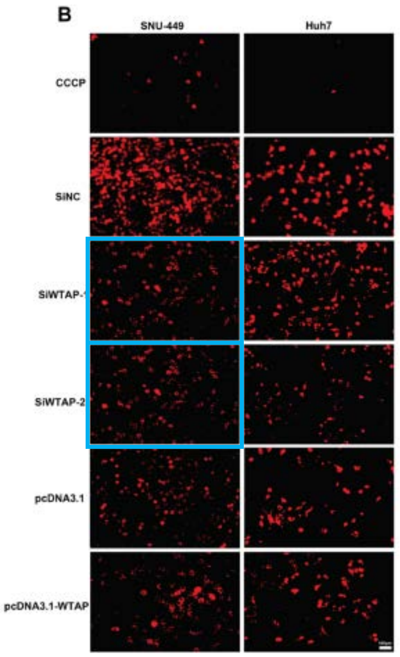


Figure 4A

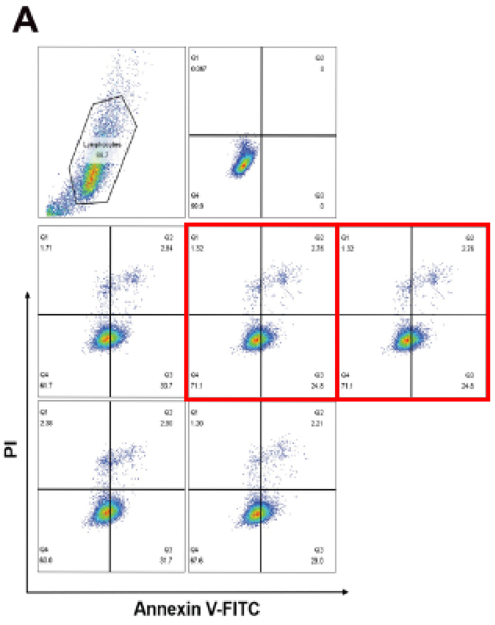


Figure 7A/D/G/J

Same b-actin blots are used, but clearly protein-of interest blots are not from the same gel

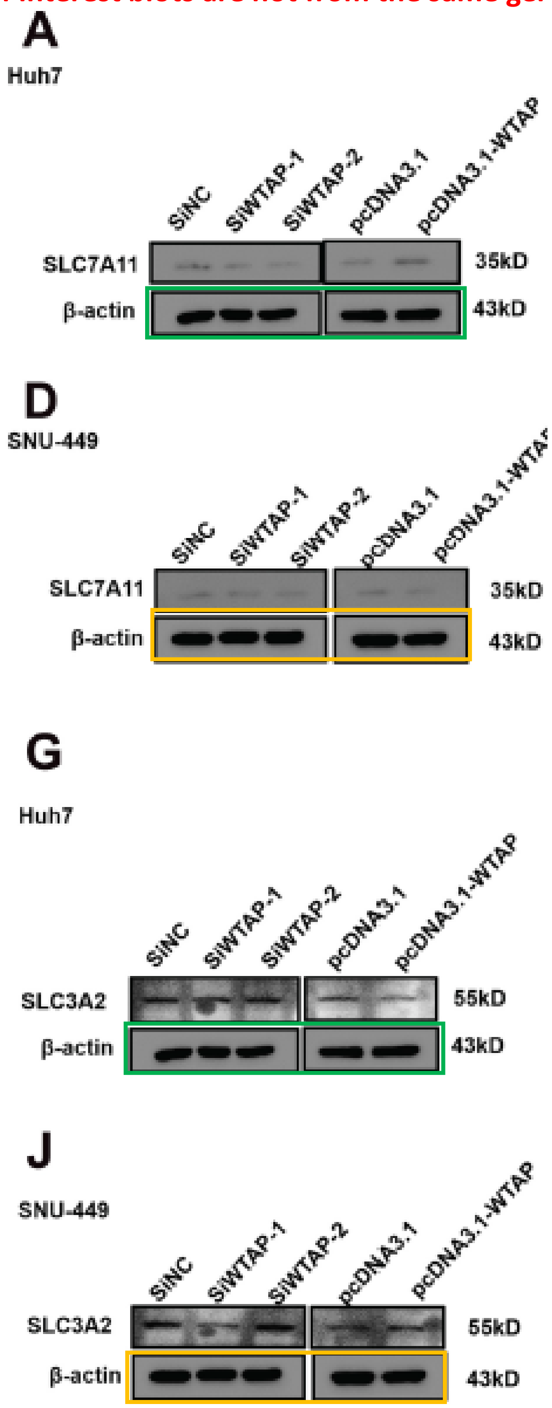

J. Cancer. 2025; 16(2): 406-416
DOI: 10.7150/jca.102360

#### Figure 2 vs 3, data repeated

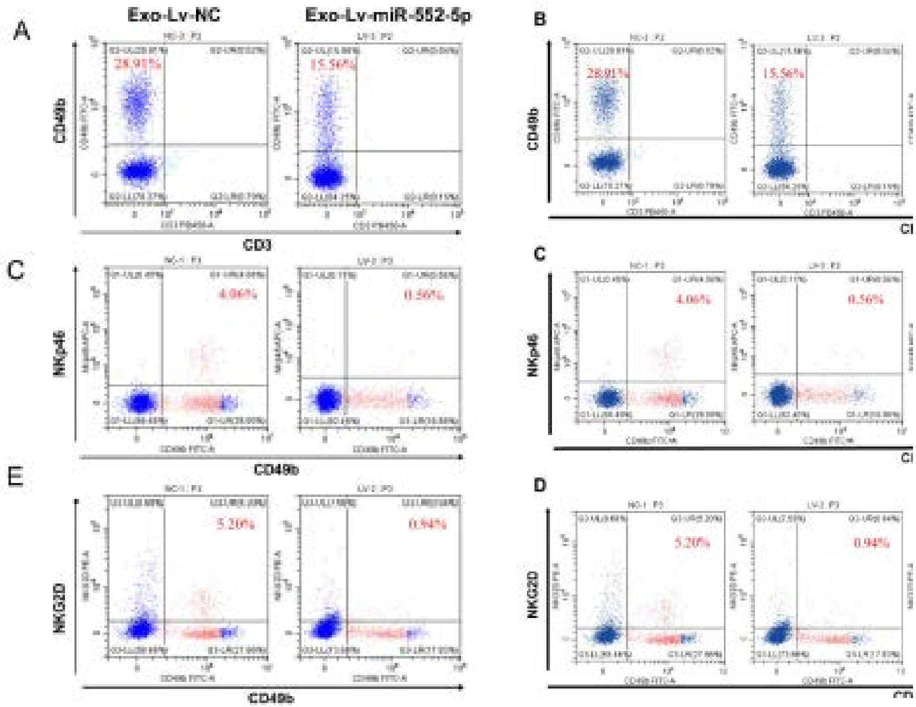


#### Figure 4G

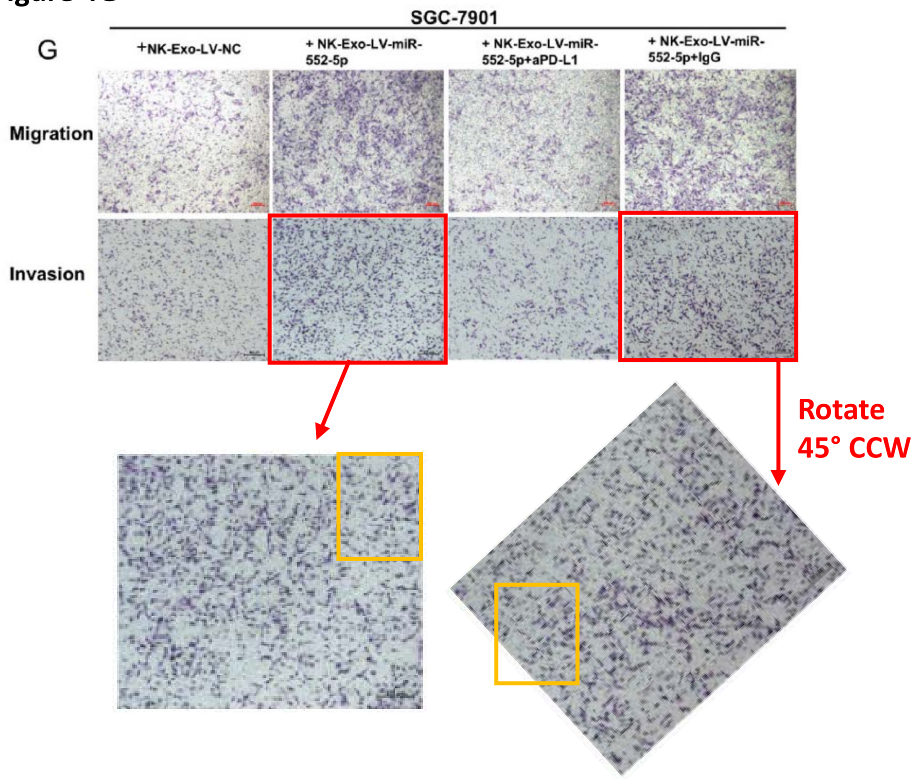

J. Cancer. 2025; 16(2): 445-459
DOI: 10.7150/jca.100951

**Figure 2G**

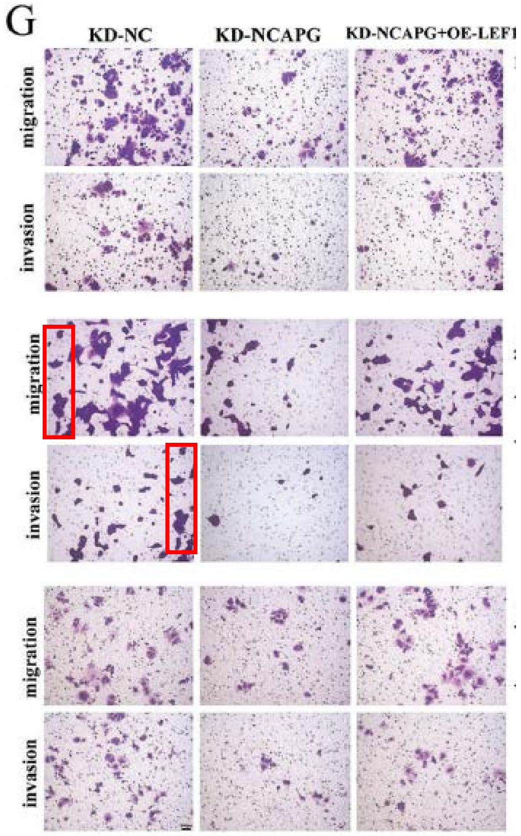


**Figure 3H**

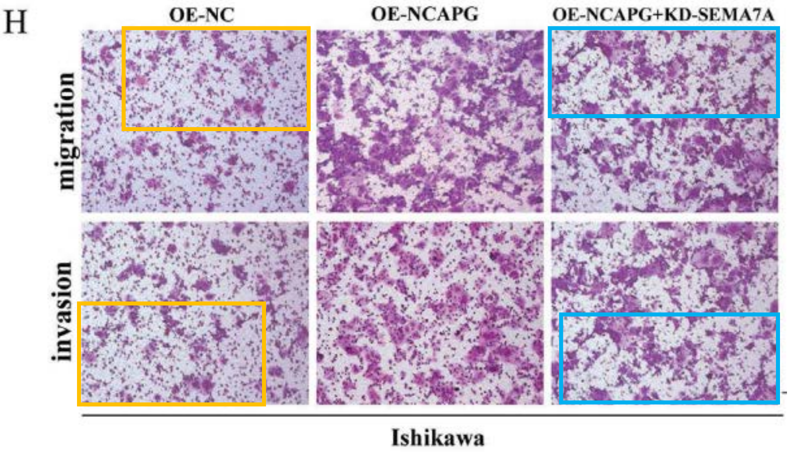

J. Cancer. 2025; 16(2): 521-532
DOI: 10.7150/jca.101285

Figure 4E/F

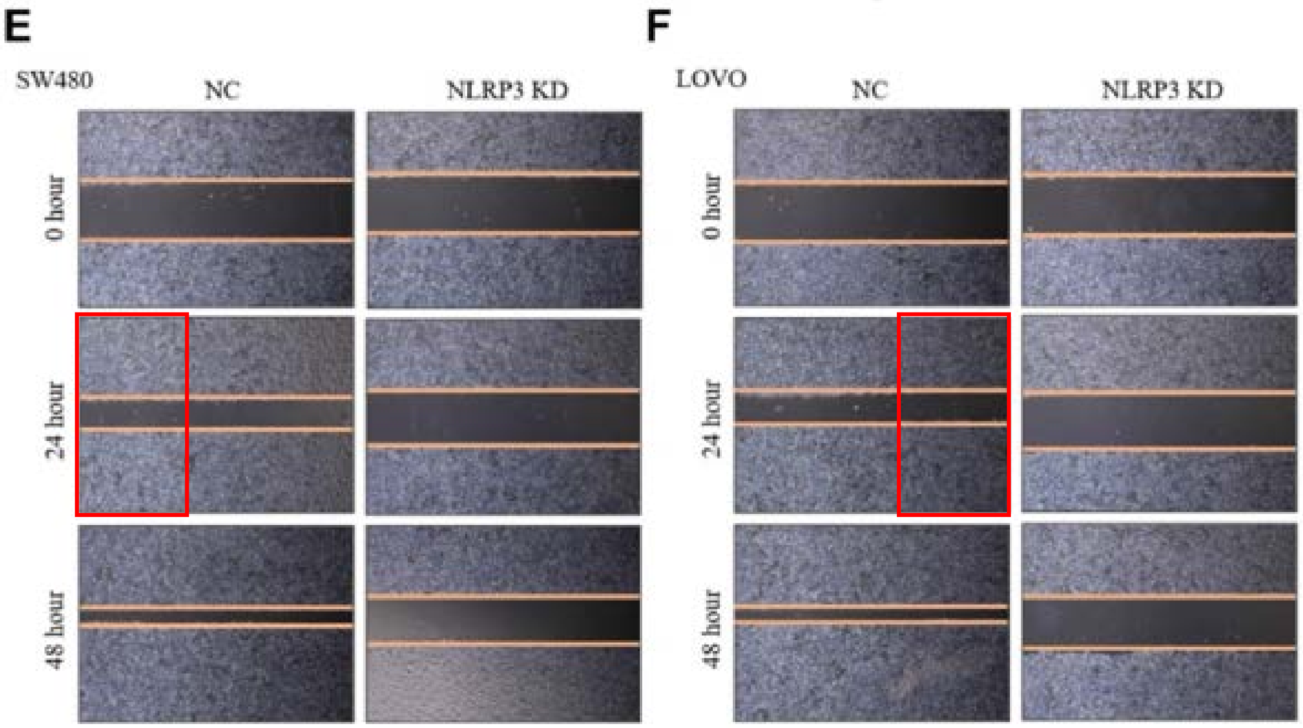


Figure 4 I/J

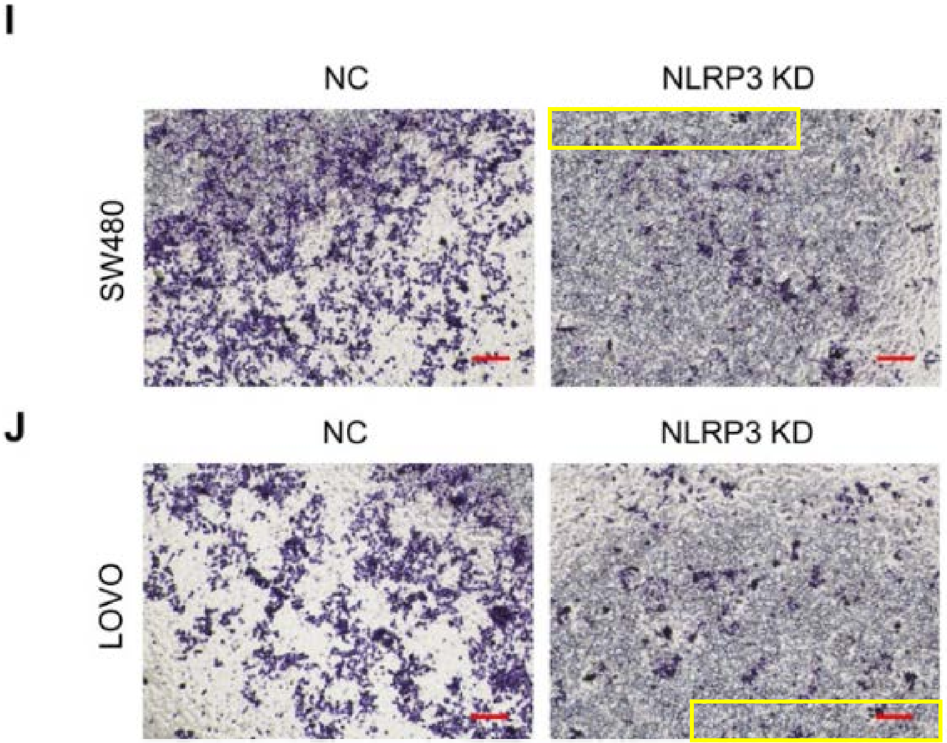

J. Cancer. 2025; 16(3): 765-783
DOI: 10.7150/jca.101798
Figure 6D / 7D

Different treatments

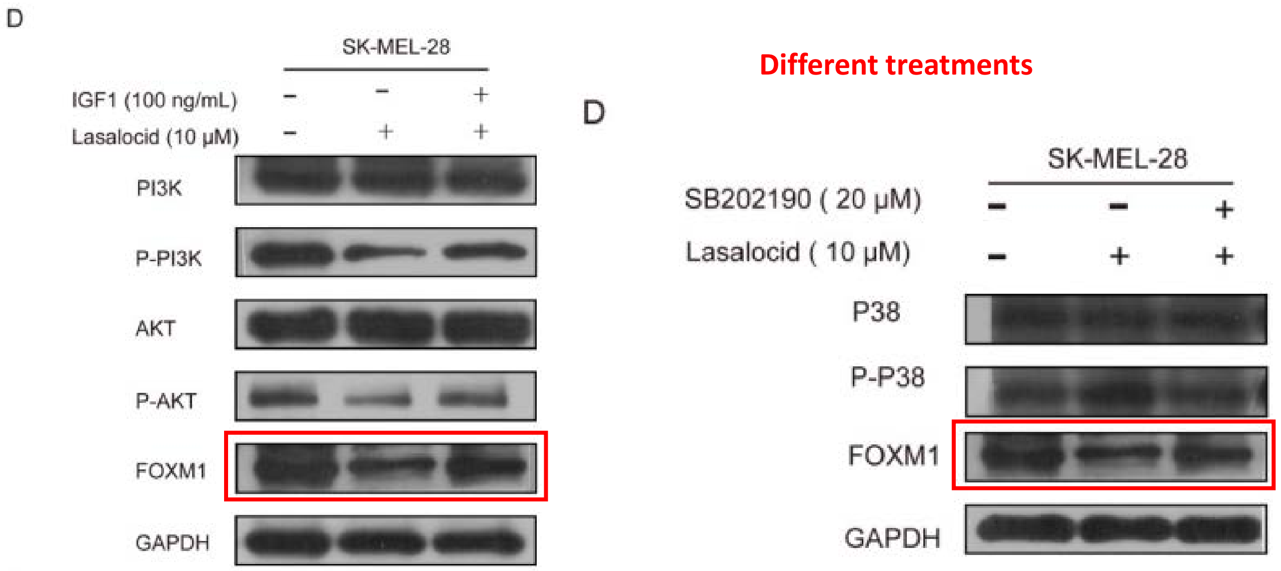

J. Cancer. 2025; 16(4): 1066-1077
DOI: 10.7150/jca.102201
Figure 4A

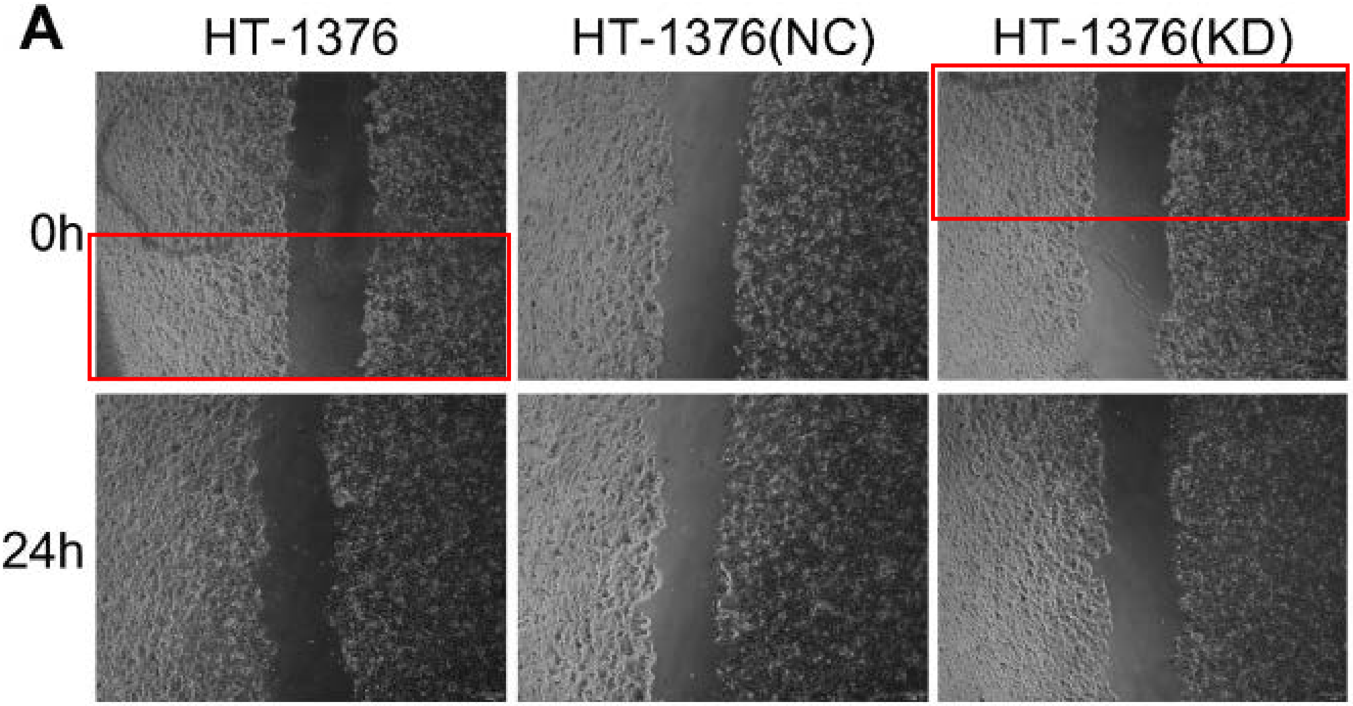

J. Cancer. 2025; 16(4): 1202-1214
DOI: 10.7150/jca.103482
Figure 2F / 4C

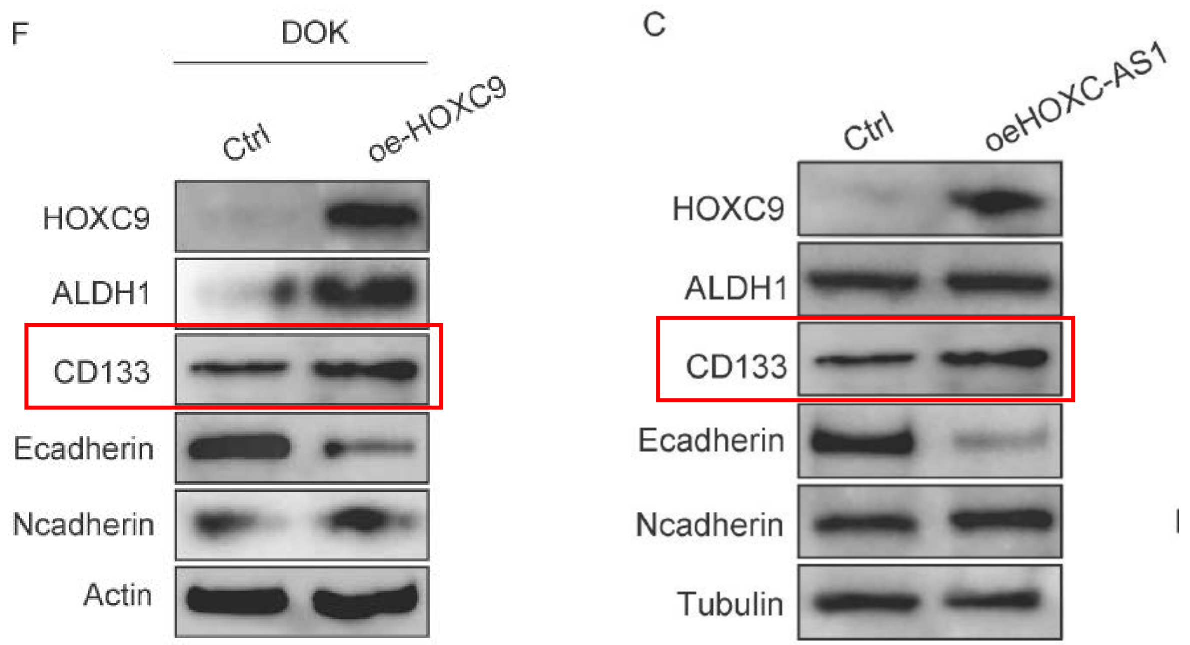


Figure 6I

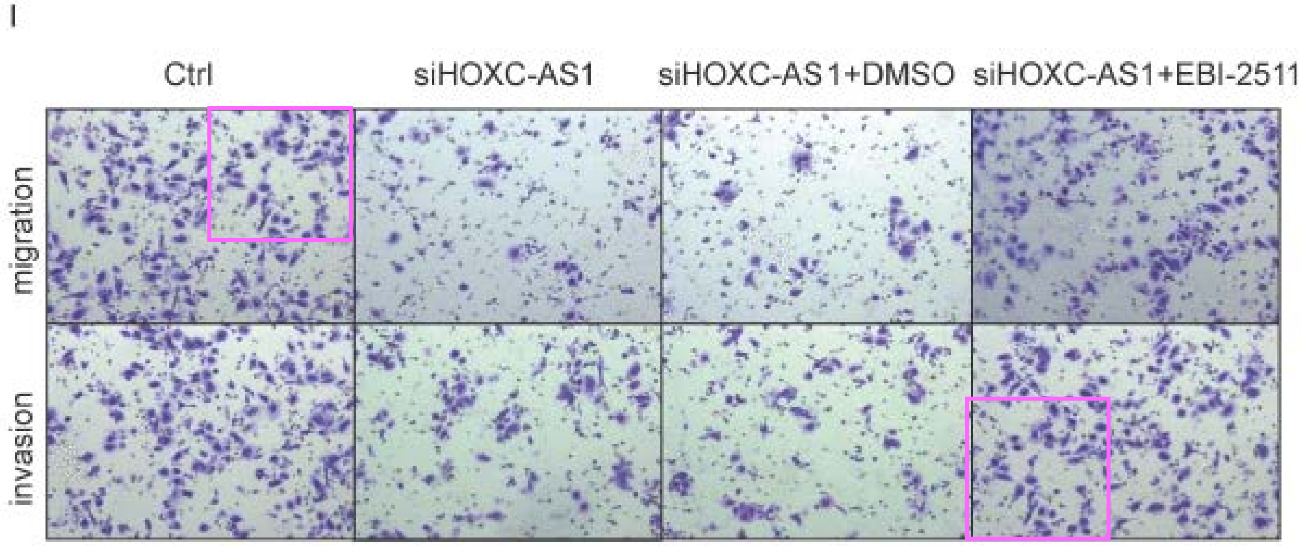

J. Cancer. 2025; 16(4): 1296-1309
DOI: 10.7150/jca.104000
Figure 4B / 6A

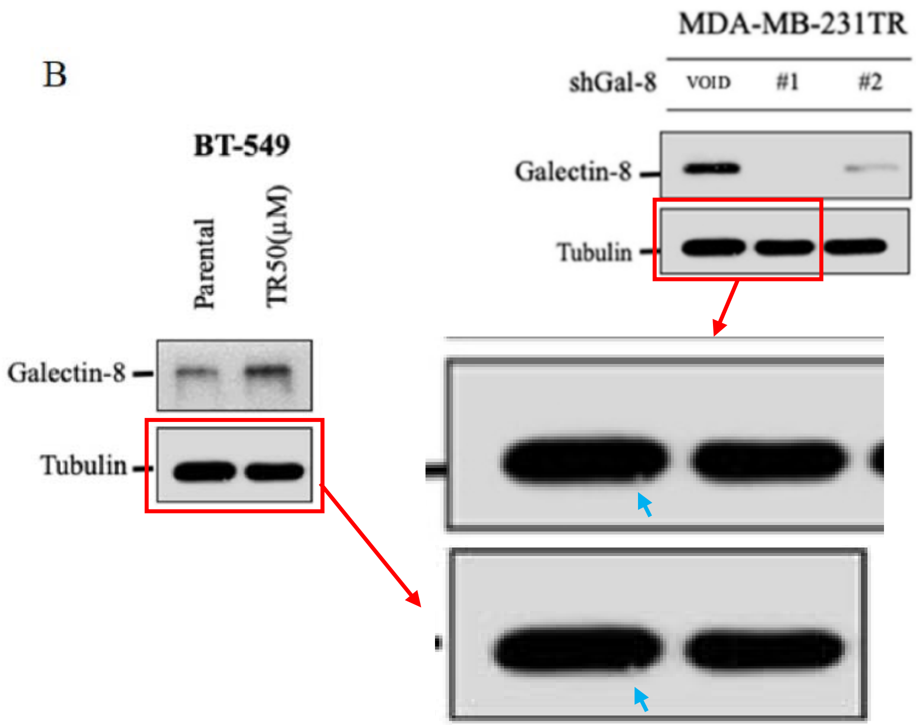


J. Cancer. 2025; 16(4): 1296-1309
DOI: 10.7150/jca.104000
Figure 5B / 6B

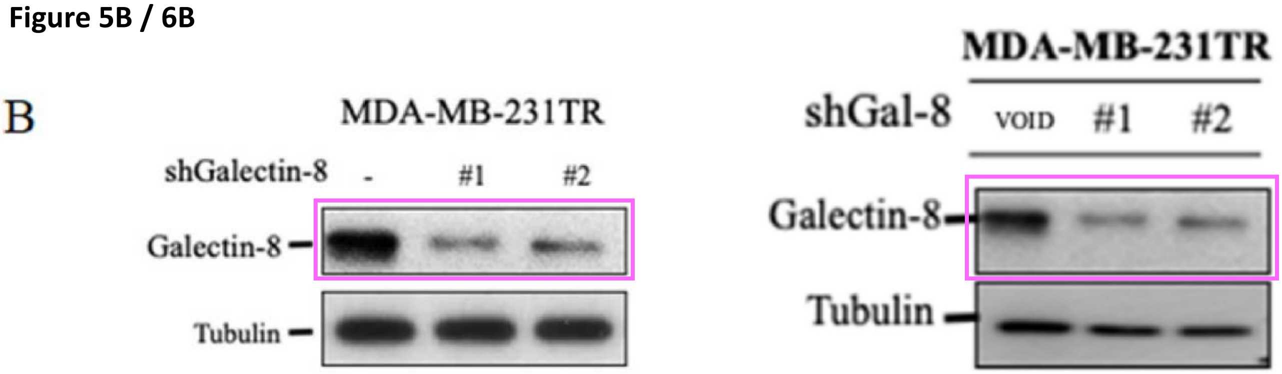

J. Cancer. 2025; 16(4): 1348-1362
DOI: 10.7150/jca.103229
Figure 2A / 5D

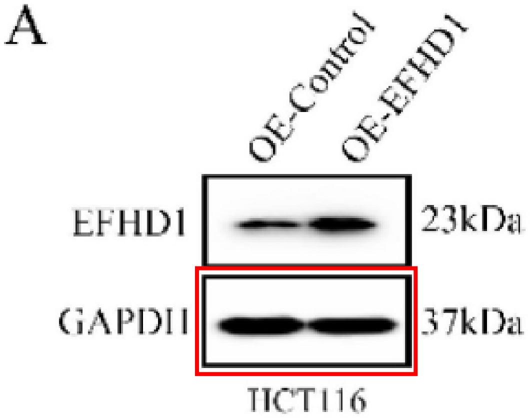


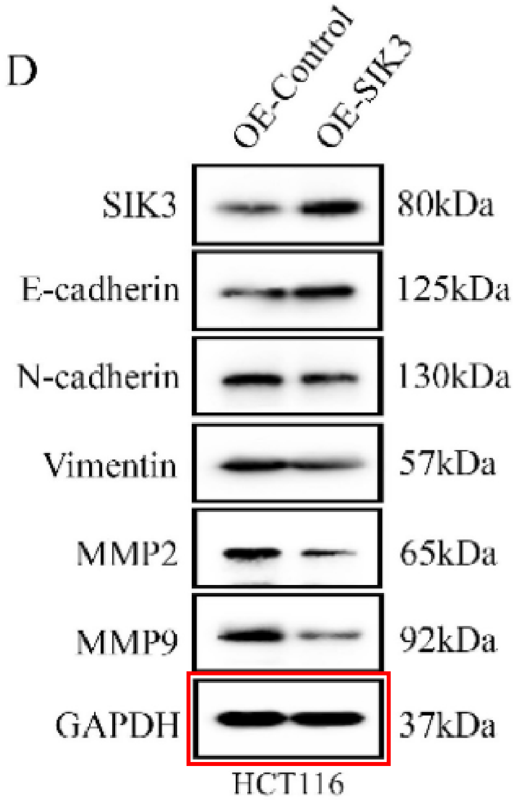


Figure 2E / 3C

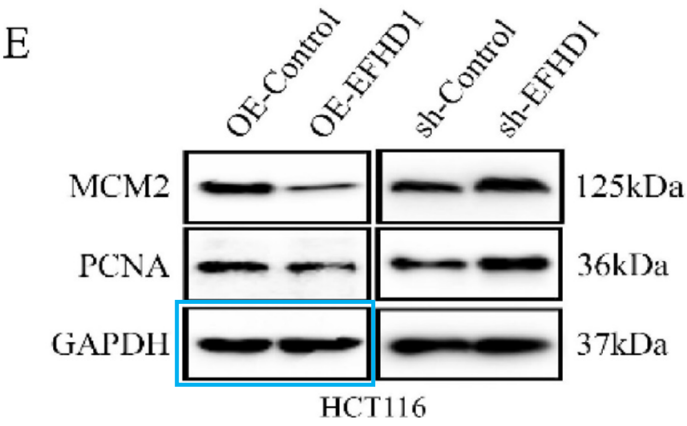


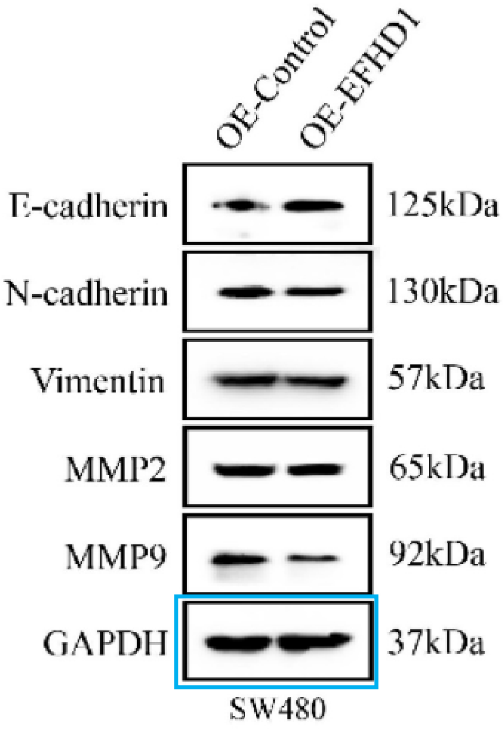

J. Cancer. 2025; 16(5): 1555-1562
DOI: 10.7150/jca.104142
Figure 2B

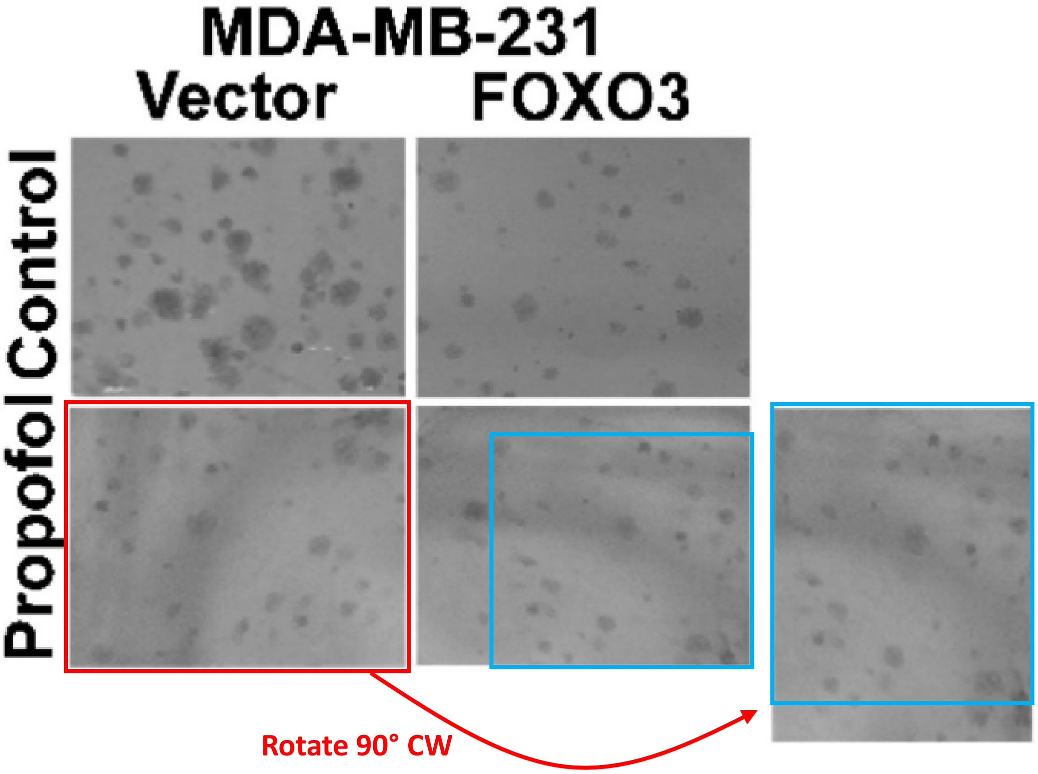

J. Cancer. 2025; 16(5): 1598-1617
DOI: 10.7150/jca.97784
Figure 2e / 2i

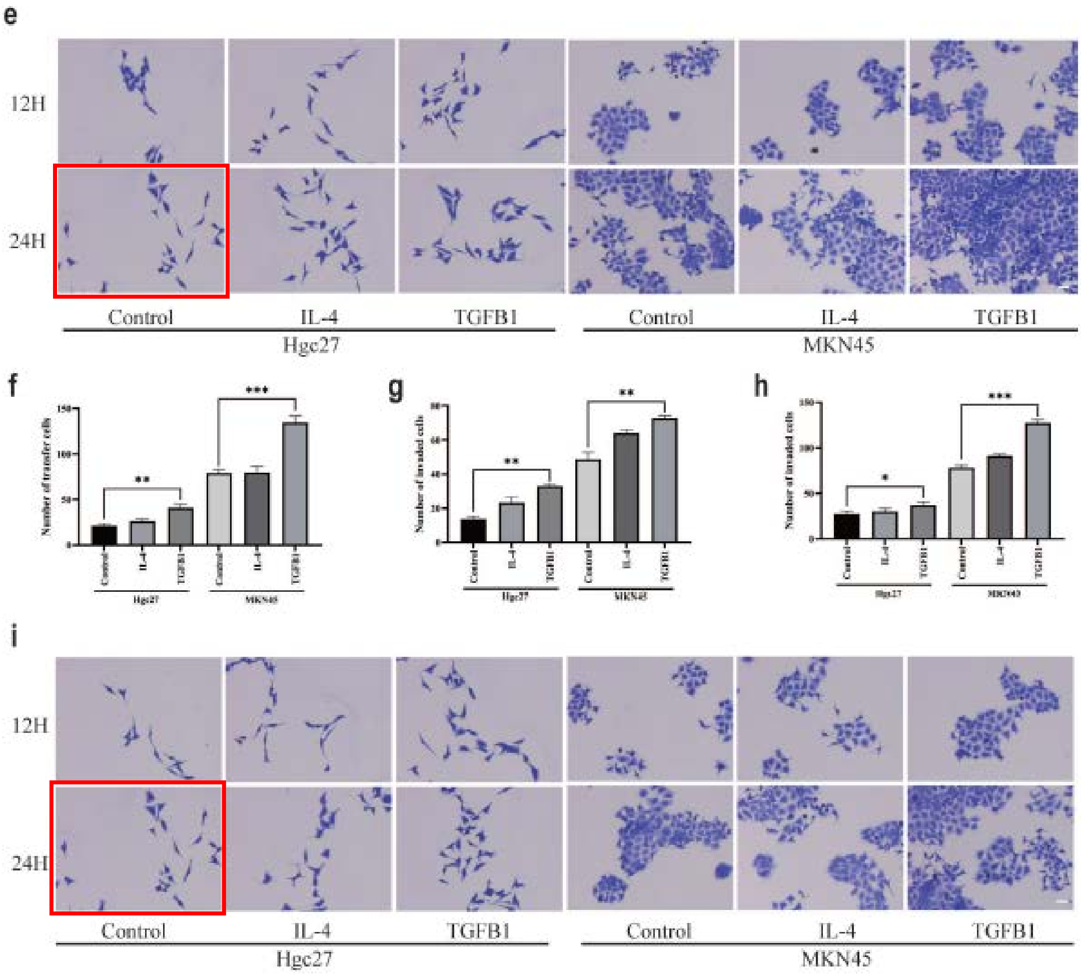

J. Cancer. 2025; 16(5): 1618-1630
DOI: 10.7150/jca.94822
Figure 2C / 5E

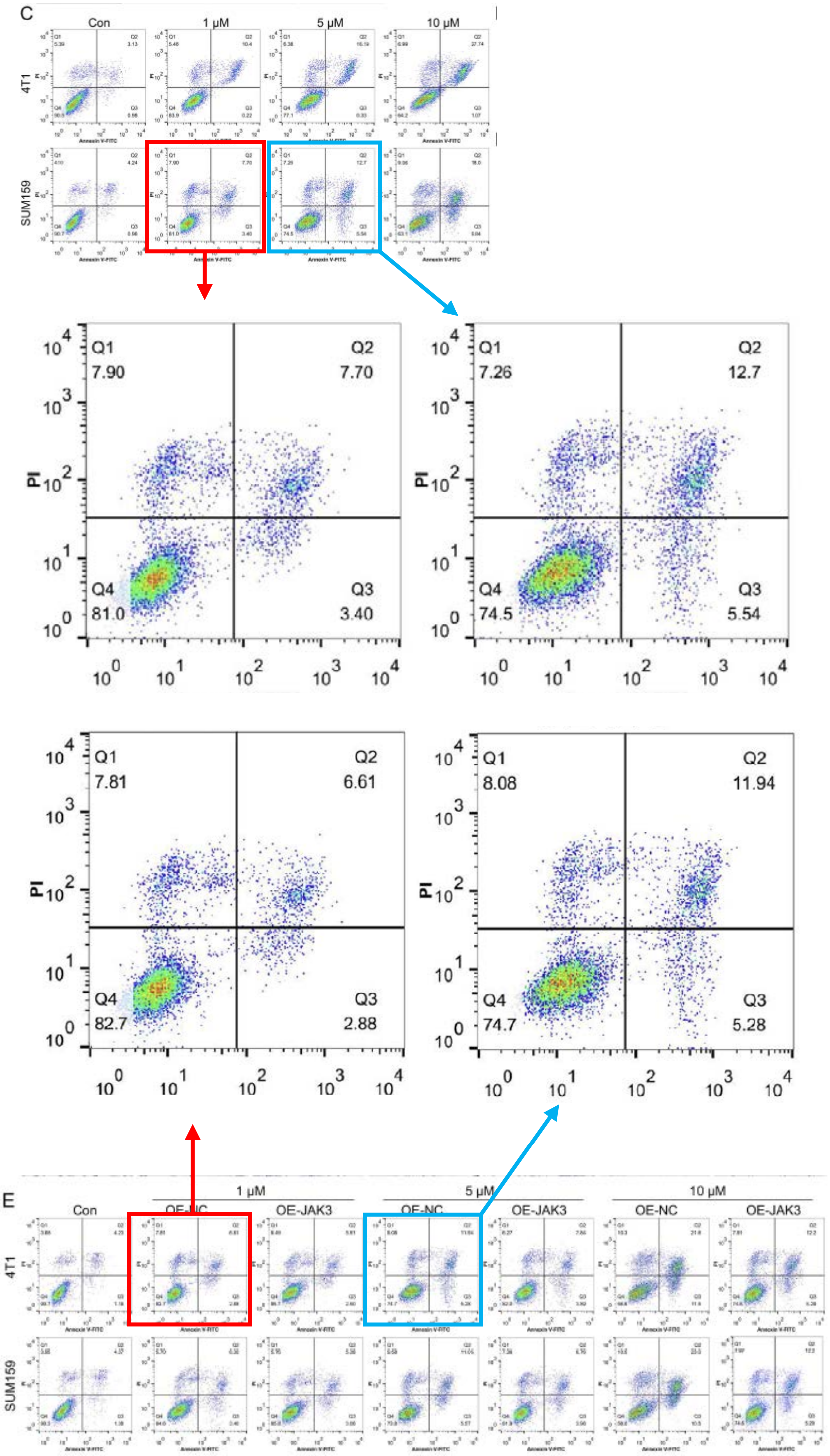

J. Cancer. 2025; 16(5): 1618-1630
DOI: 10.7150/jca.94822
Figure 5E

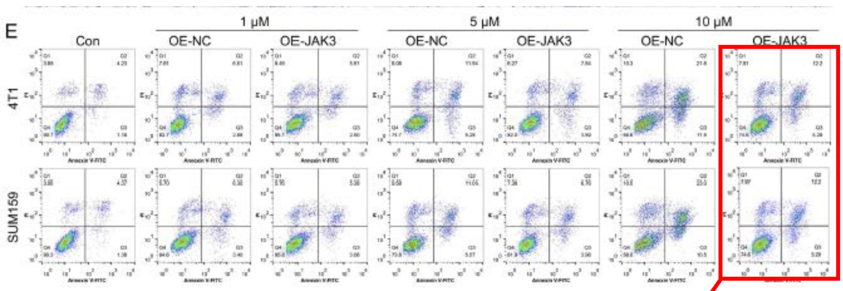


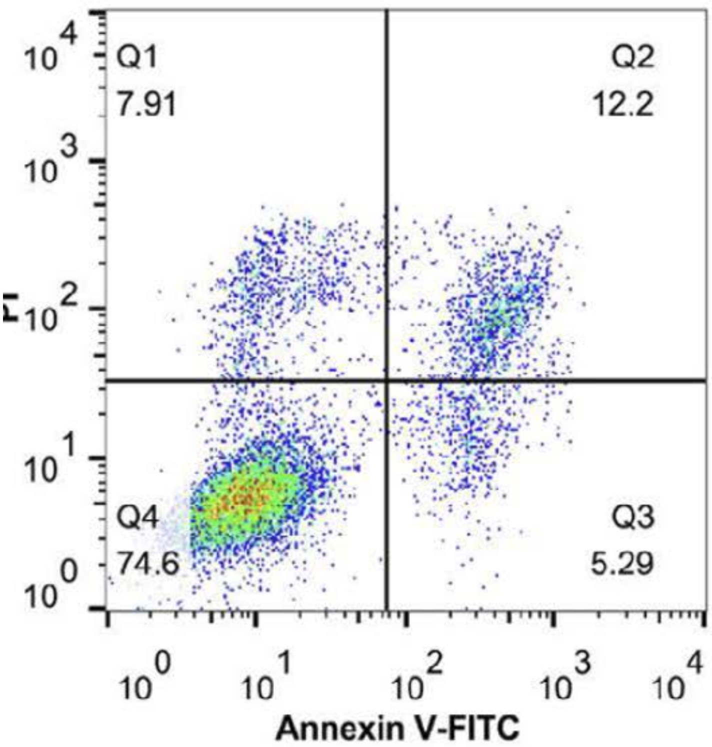


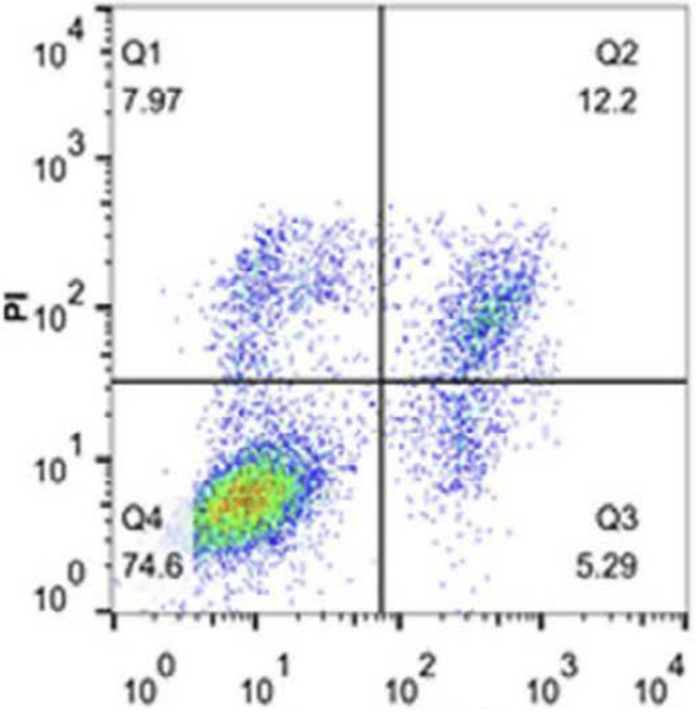

J. Cancer. 2025; 16(5): 1618-1630
DOI: 10.7150/jca.94822
Figure 2F / 4G

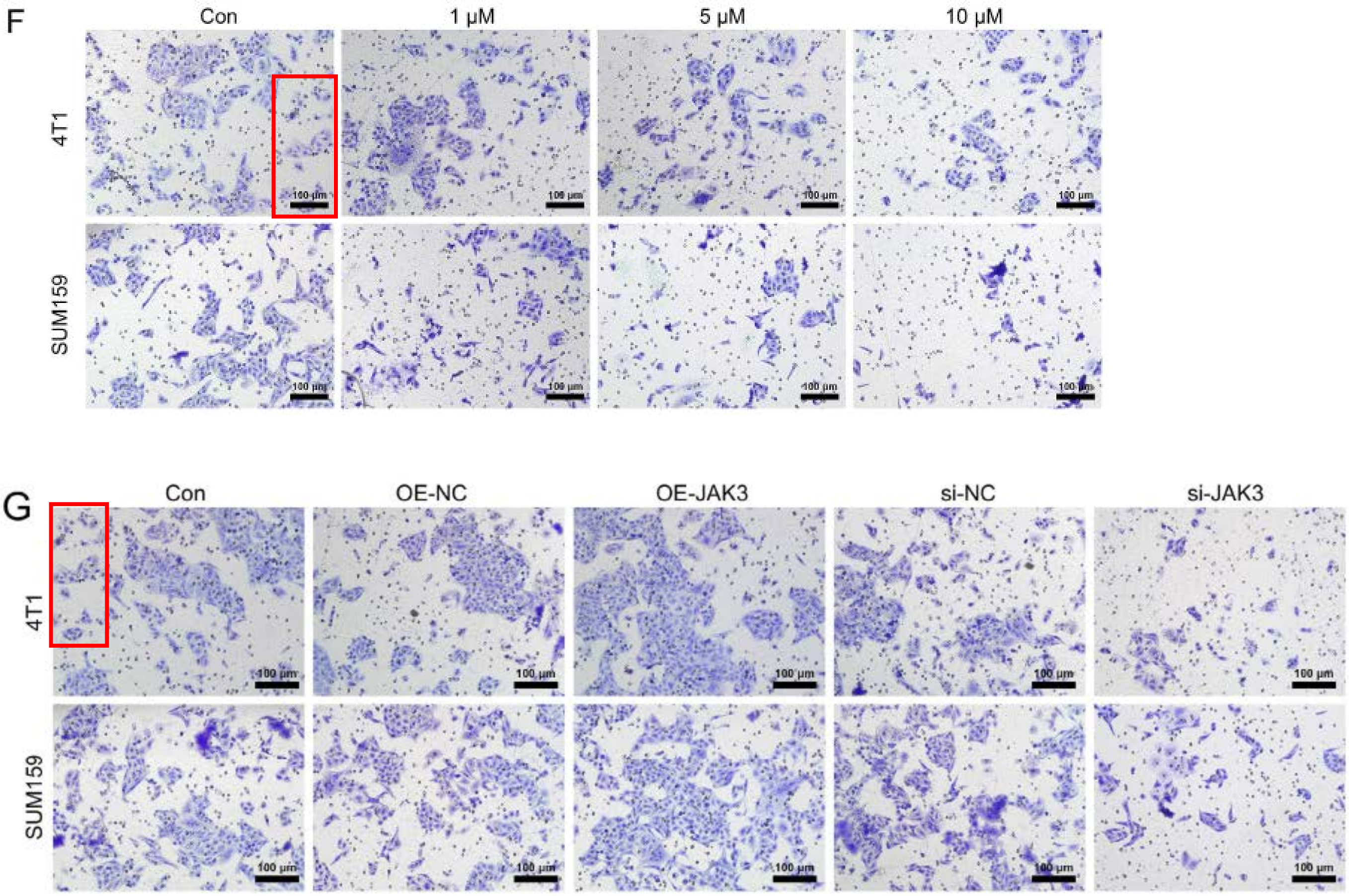

**J. Cancer. 2025; 16(5): 1694-1708**
**DOI: 10.7150/jca.105425**
**Figure 3A / 3C**

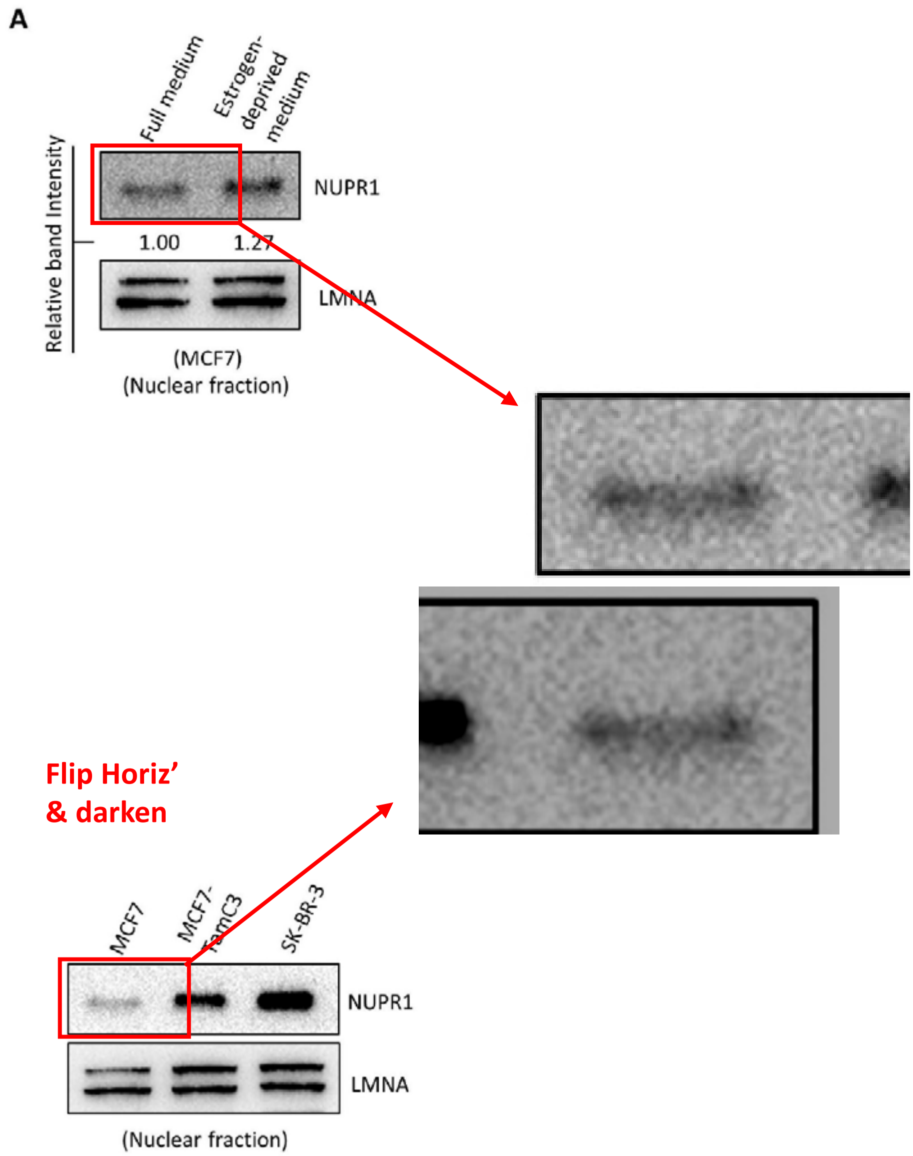

J. Cancer. 2025; 16(5): 1905-1917
DOI: 10.7150/jca.105557
Figure 9C / 10F

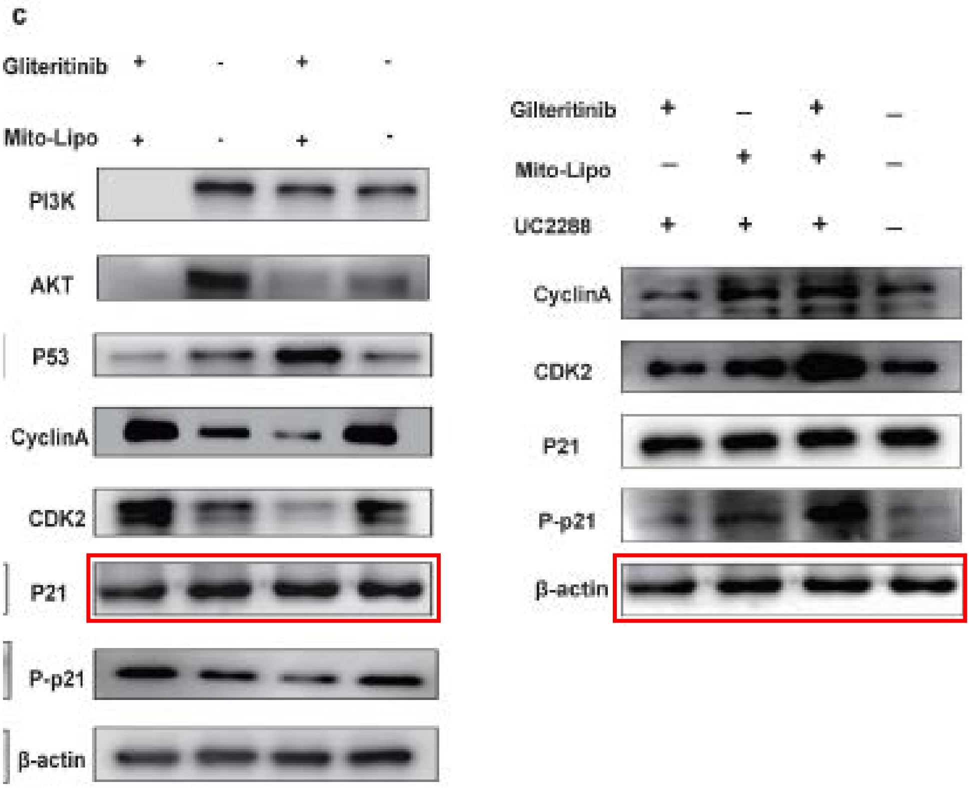

J. Cancer. 2025; 16(7): 2087-2102
DOI: 10.7150/jca.104389
Figure 10B

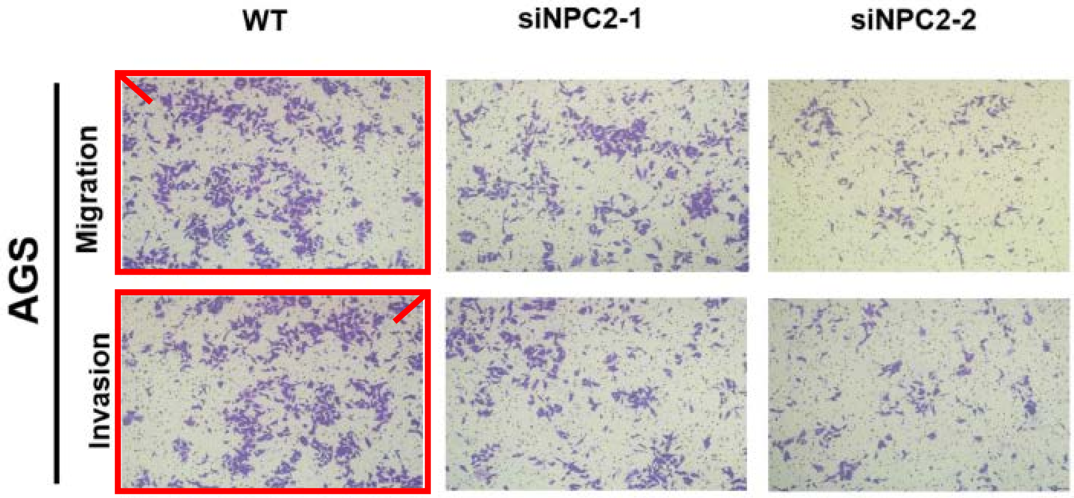

J. Cancer. 2025; 16(7): 2233-2249
DOI: 10.7150/jca.105120
Figure 3C/D

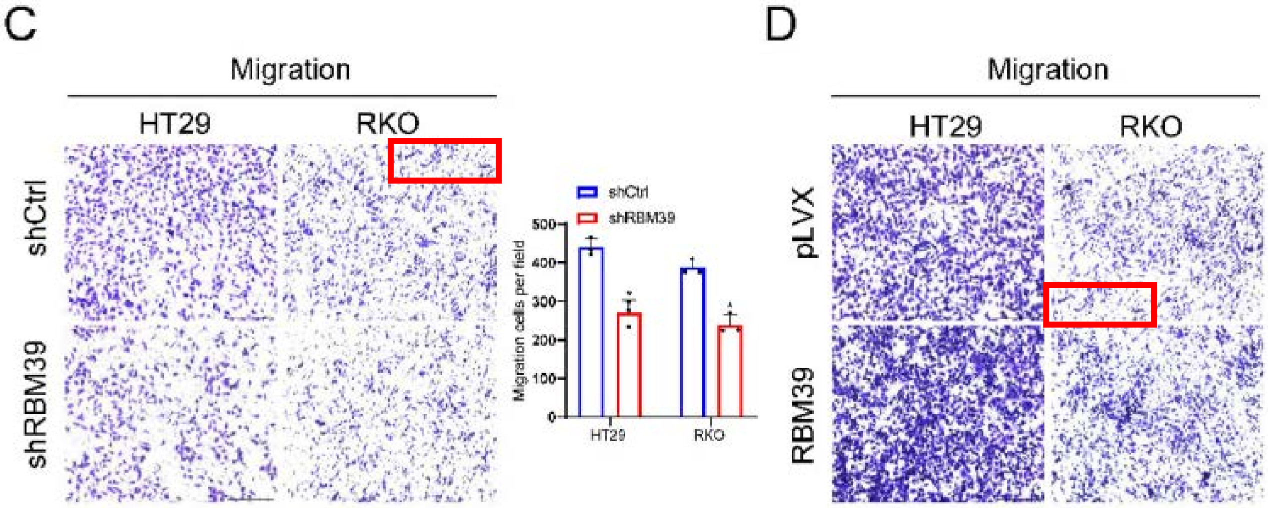


Figure 3E

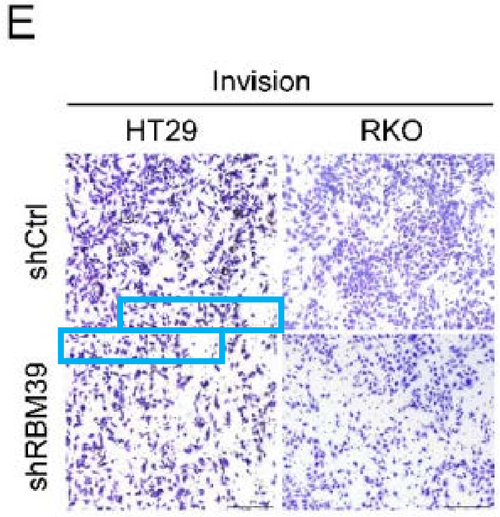

J. Cancer. 2025; 16(7): 2434-2448
DOI: 10.7150/jca.99599
Figure 3F

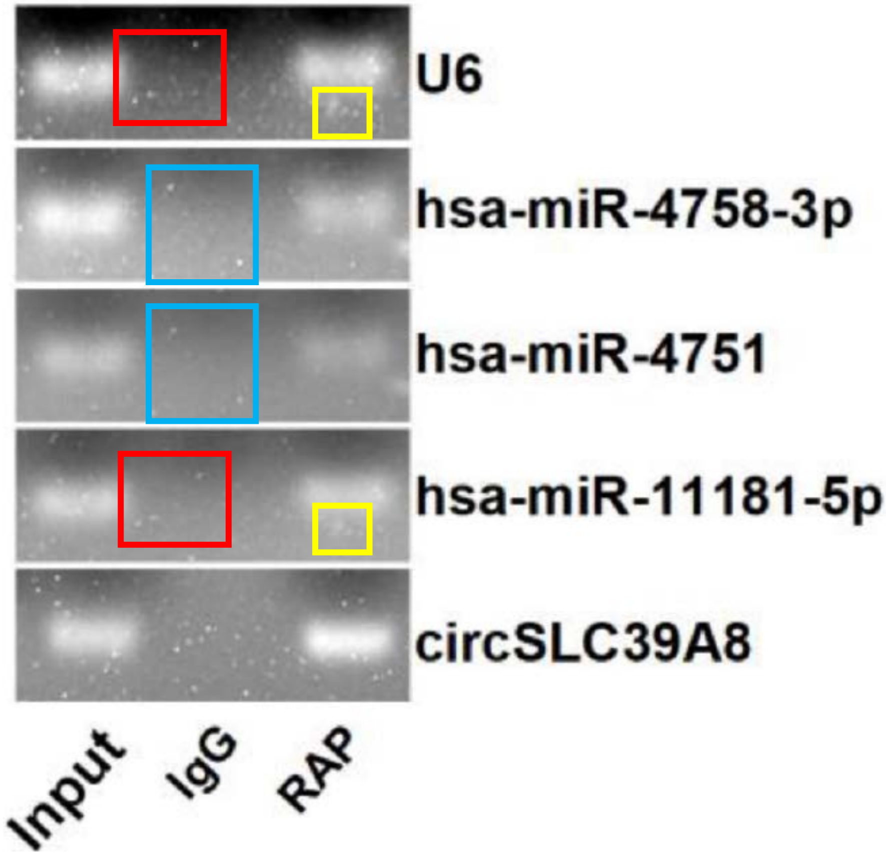


Figure 4D

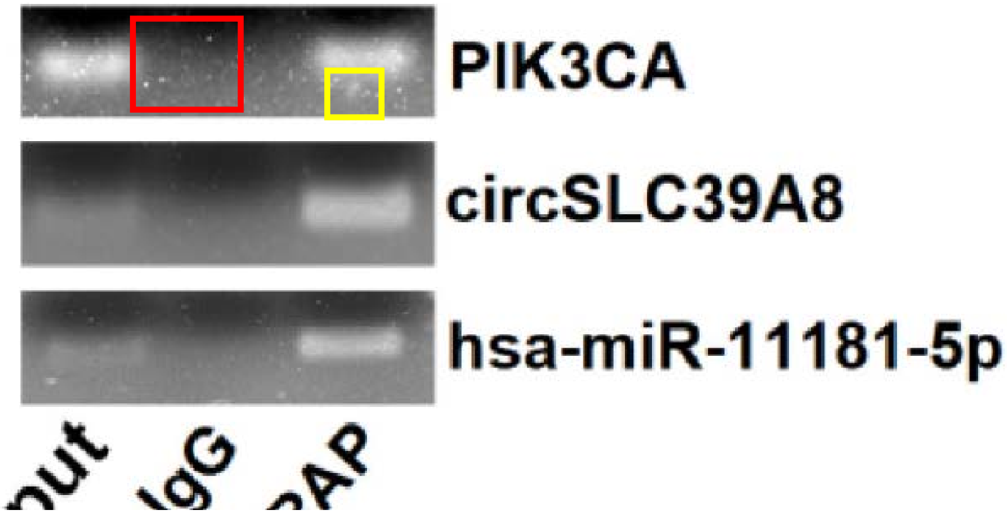

J. Cancer. 2025; 16(8): 2466-2475
DOI: 10.7150/jca.98431
Figure 3A / 5A

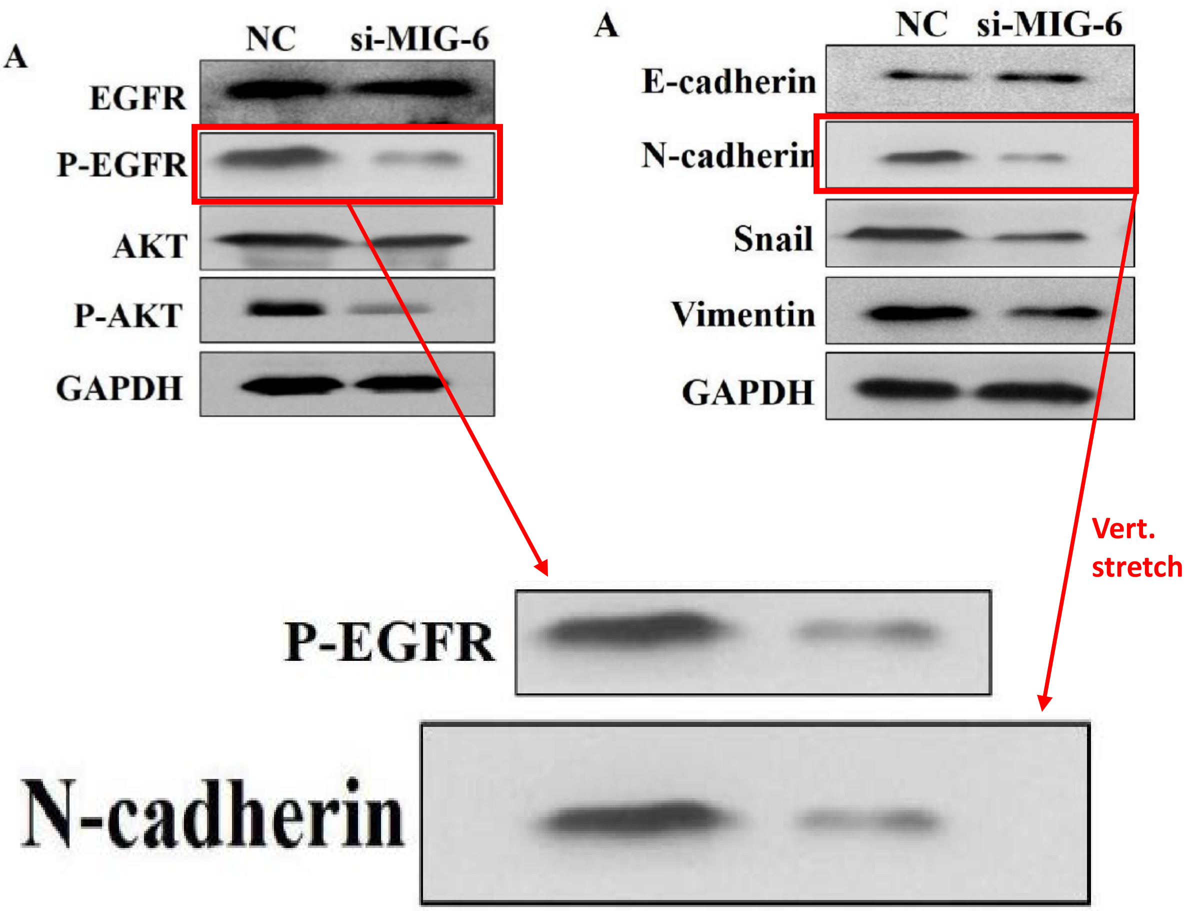

J. Cancer. 2025; 16(10): 3080-3093
DOI: 10.7150/jca.111459
Figure 2G / 7E

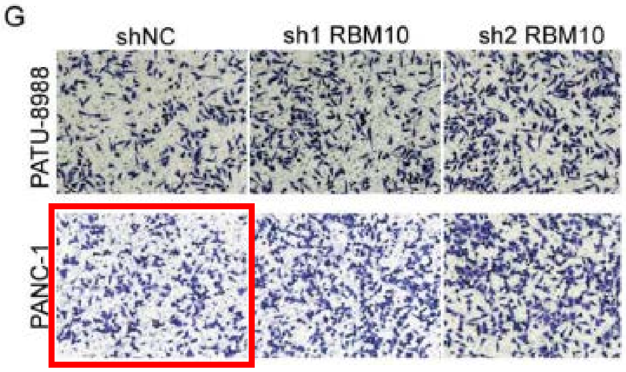


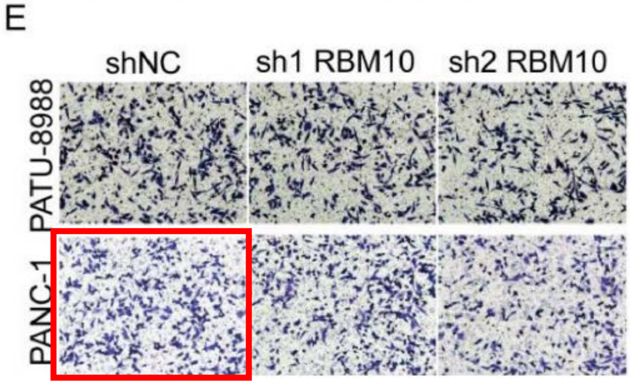


Different conditions in 7E (Rx w. AZD1480)

J. Cancer. 2025; 16(10): 3180-3191
DOI: 10.7150/jca.114495
Figure 4

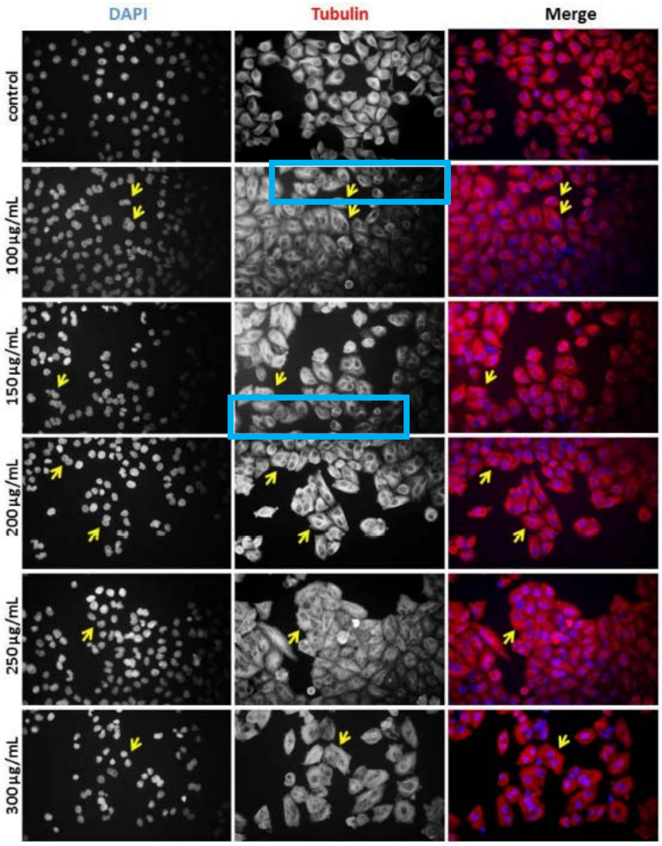

J. Cancer. 2025; 16(11): 3329-3342
DOI: 10.7150/jca.115892
Figure 3B/C

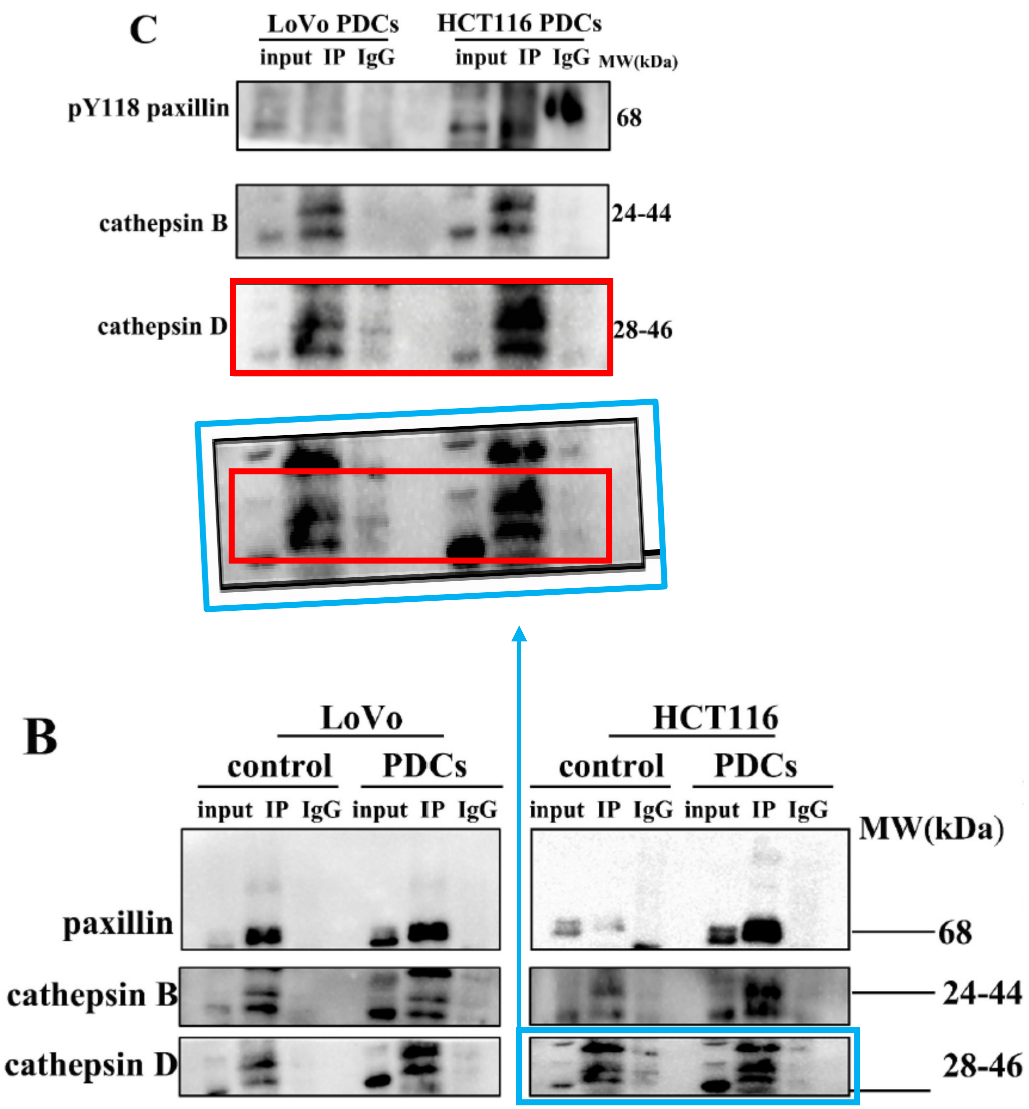

J. Cancer. 2025; 16(12): 3729-3745
DOI: 10.7150/jca.111383
Figure 1I

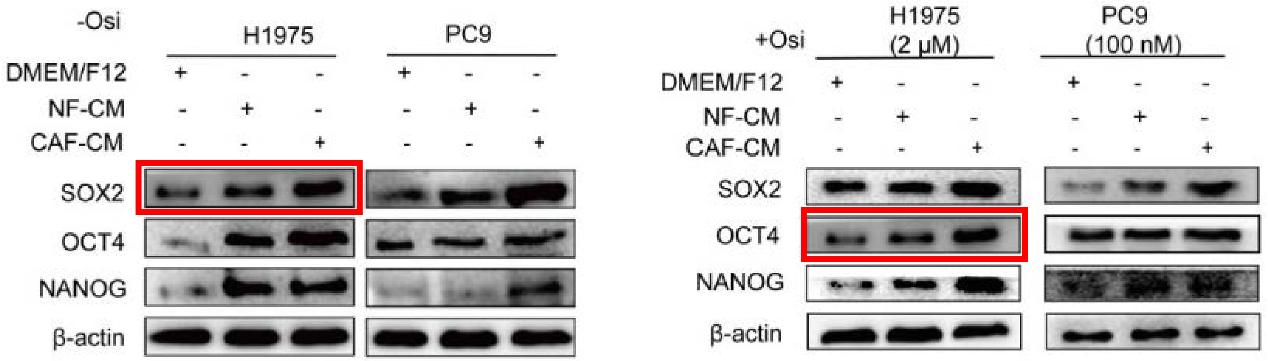


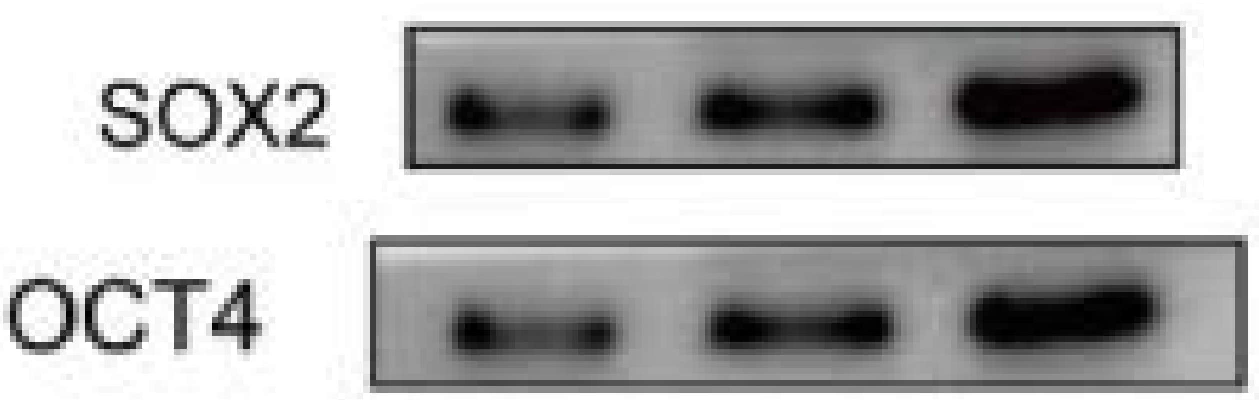

J. Cancer. 2025; 16(13): 3928-3941
DOI: 10.7150/jca.112791
Figures 7F/G vs. 8K/L

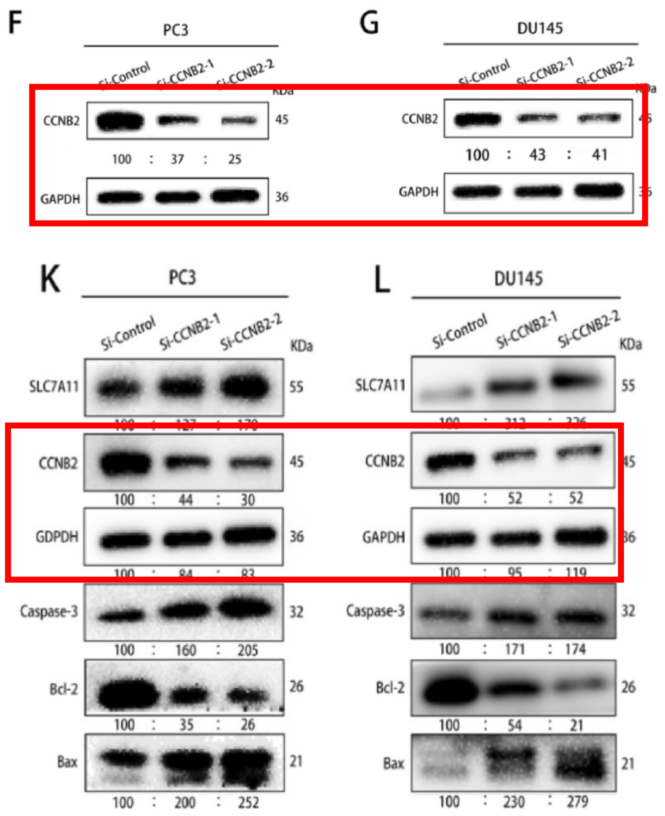


Same blots but faded and resized, quant' #s below are different

Figure 8E/F

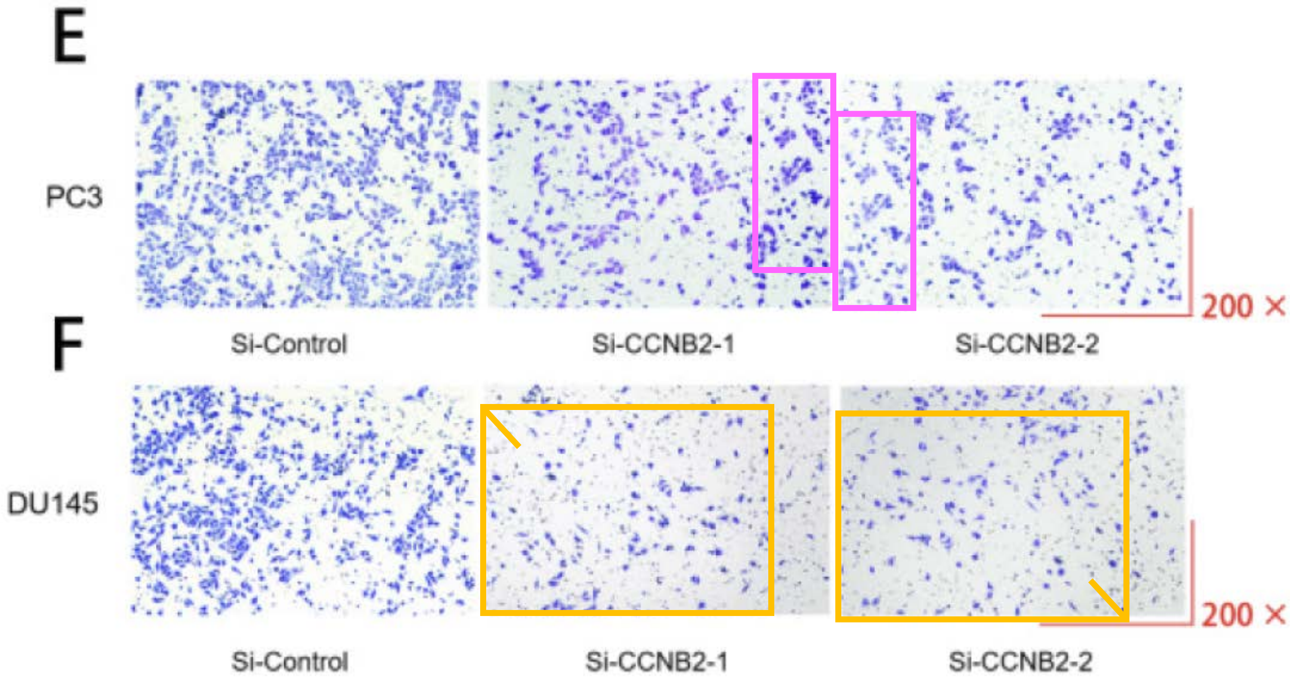

J. Cancer. 2025; 16(14): 4117-4129
DOI: 10.7150/jca.119549
**Figure 1 vs. 2**

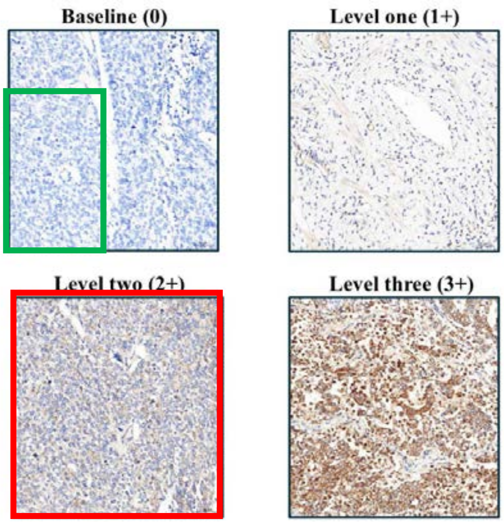


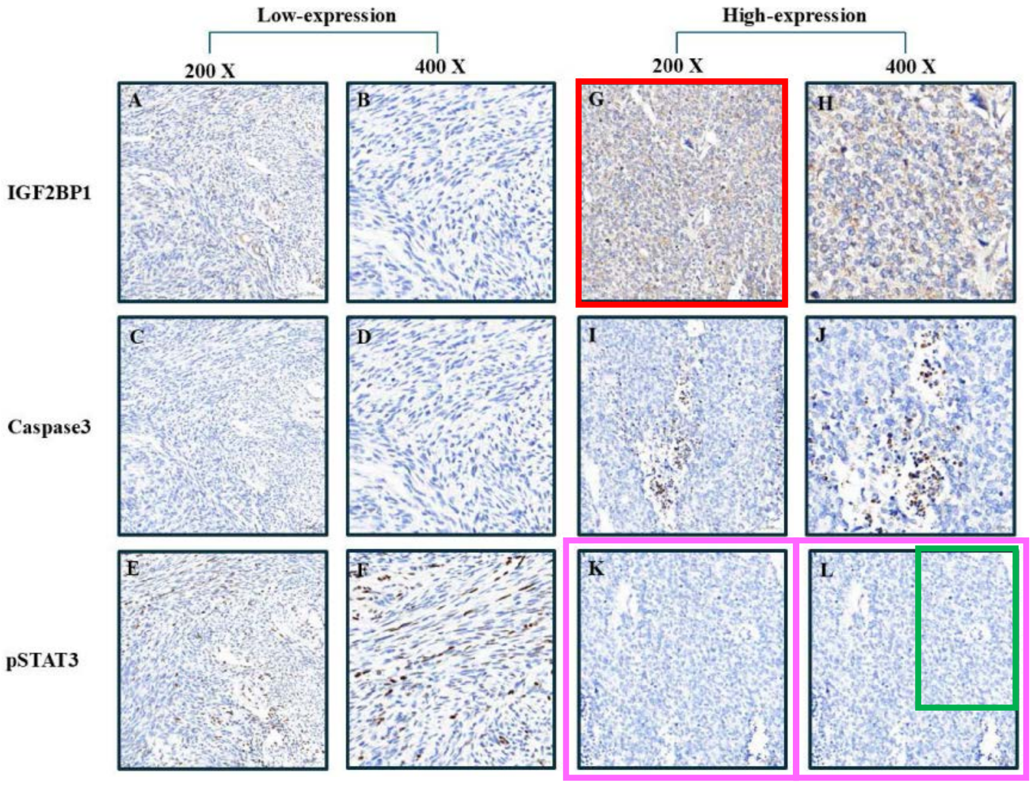

J. Cancer. 2025; 16(15): 4284-4291
DOI: 10.7150/jca.119706

Figure 5A/B

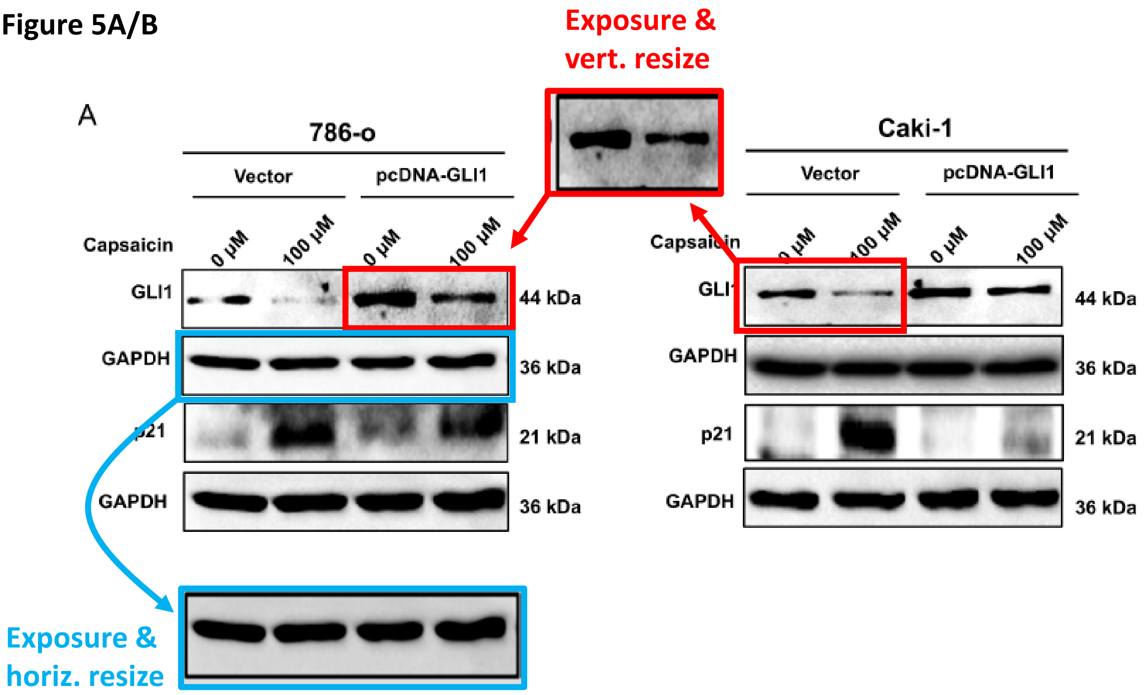
